# A dynamic displacement mechanism drives protein import into mitochondria

**DOI:** 10.64898/2026.01.14.699430

**Authors:** Iva Sučec, Undina Guillerm, Jakob Schneider, Iniyan Ganesan, Niklas Webling, Anna Kapitonova, Conny Steiert, Frank Schmelter, Antoine Feignier, Rajkumar Singh, Laura F. Fielden, Julia M. Dung, Doron Rapaport, Beate Bersch, Karin B. Busch, Francois Dehez, Nils Wiedemann, Paul Schanda

**Author notes:** Walter and Eliza Hall Institute of Medical Research, 1G Royal Parade, Parkville, VIC 3052, Australia; and Department of Medical Biology, University of Melbourne, Melbourne, VIC 3000, Australia. I.S., U.G. and J.S. contributed equally to this work. N.Wi., K.B.B. and F.D. contributed equally to this work.

## Abstract

Most mitochondrial proteins are produced in the cytosol and imported through the translocase of the outer mitochondrial membrane (TOM) to reach their final destination. Although this protein entry gate has been structurally characterized, it remains unclear how precursor proteins are handed off from the cytosolic receptor domains to the translocation pore. Here we show that the cytosolic domain of Tom22—traditionally viewed as the central TOM receptor—acts not as a structured scaffold but as a largely disordered, flexible segment that plays an active role in precursor transfer. Atomic-level structural techniques and in vivo experiments identified a conserved short linear motif that forms a transient α-helical element within this disordered domain. By binding to the canonical precursor protein binding sites of the receptors Tom20 and Tom70, this critical α-helical segment acts as a precursor protein displacement element (PPDE). This competitive interaction facilitates the release of preproteins directly above the import pore, and thereby drives translocation across the outer mitochondrial membrane. These findings reveal that flexibility, rather than rigid structure, underlies the central transfer step of mitochondrial outer-membrane protein translocation. Our results point to a versatile mechanism for ligand displacement in chaperone, receptor, and transport systems that must balance selective binding with efficient release.

## Introduction

Mitochondria are crucial organelles fundamental to numerous metabolic and signaling pathways within eukaryotic cells^1, 2^. Almost all mitochondrial proteins are produced in the cytosol, and among these, more than 80%^3, 4^ are imported post-translationally^5–12^. After their synthesis, precursor proteins (preproteins) are guided by cytosolic chaperones to the organelle surface, where they are recognized by receptor proteins^12–16^. Proteins destined for the mitochondrial matrix, inner membrane, intermembrane space, and some outer-membrane (OM) proteins are translocated through the TOM complex. Defects in these receptor–client interactions underlie a range of neuromuscular and metabolic disorders^17^.

The TOM_core_ adopts a dimeric structure comprising two Tom40 β-barrel translocation pores.^18–27^ Centrally positioned between the two Tom40 pores are two copies of Tom22, a single transmembrane (TM) helix protein. Accessory proteins Tom5, Tom6, and Tom7 flank each pore (Fig. 1A). Analysis of the hydrophobic and electrostatic properties of the barrel interior, combined with cross-linking, has provided valuable information regarding the translocation pathway^21, 28, 29^. The two receptor proteins Tom20 and Tom70 are loosely associated with the TOM_core_^30–38^ and can diffuse independently in the OM, as shown directly for Tom20^39^.

**Figure 1.**
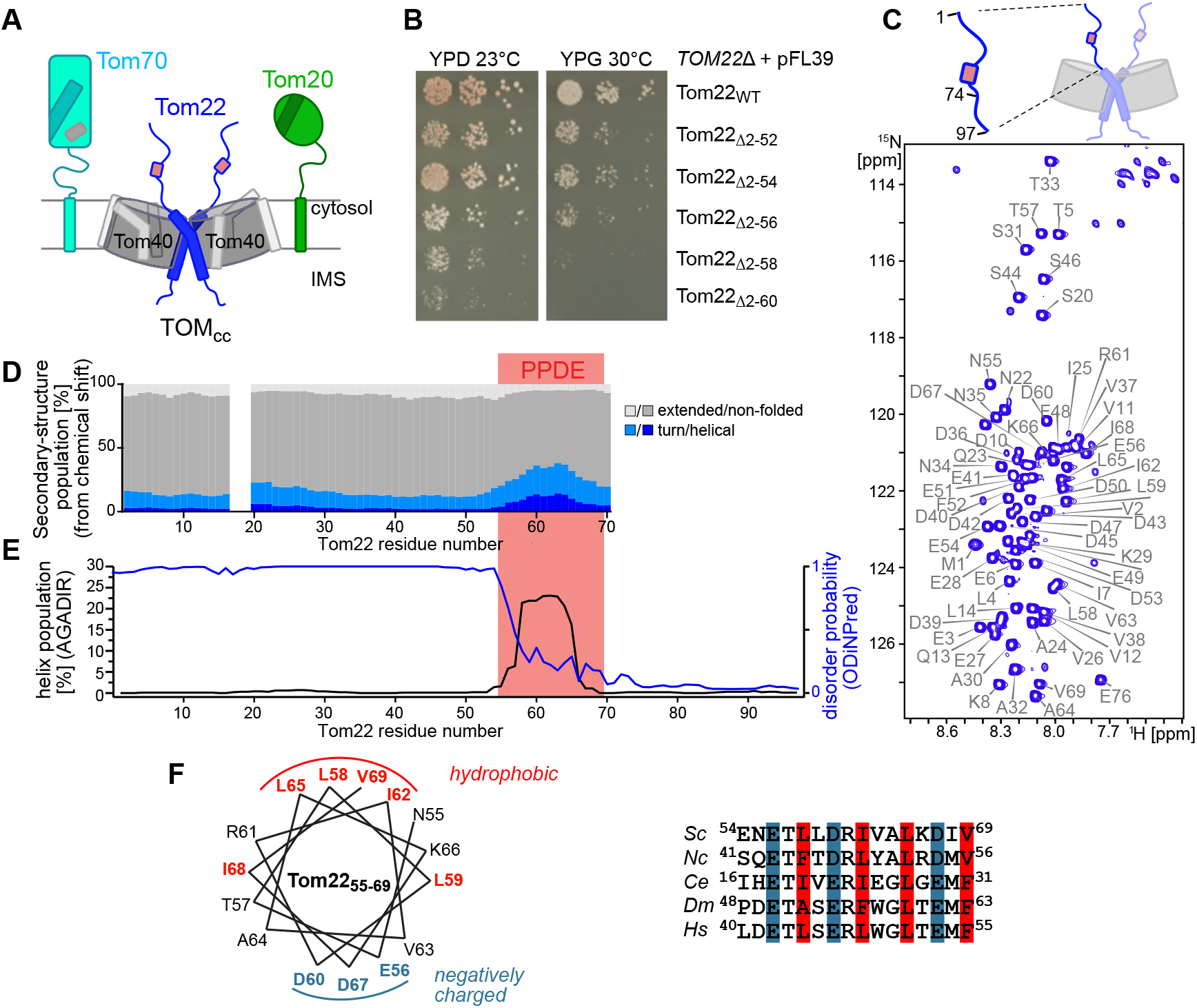
Tom22_cyt_ is an intrinsically disordered protein and only a short conserved transient α-helix is essential for function. **A**, Schematic representation of the TOM_core_ with two copies of Tom22 (in blue) at the pore interfaces and loosely associated Tom20 (green) and Tom70 (cyan) receptors. Preprotein binding sites on Tom20 and Tom70 are indicated in darker color and the Hsp70-binding pocket on Tom70 is indicated in grey. Tom22_PPDE_ comprising residues 55-69 is indicated in pink. **B**, Growth analysis of yeast strains expressing wild-type (WT) or N-terminal truncations of the cytosolic domain of Tom22. Deletion of residues further than residue 54 cause strong growth defects on fermentable (glucose, YPD) and non-fermentable (glycerol, YPG) medium. Experiments to verify protein levels are shown in Figure S2. **C**, ^1^H-^15^N correlation spectrum of *Sc*Tom22_1-74_ with residue-specific resonance assignments. The spectrum of *Sc*Tom22_1-97_ is shown in Fig. S5. **D**, Secondary structure of *Sc*Tom22_1-74_ determined from assigned ^1^H, ^15^N, ^13^C^α, 13^C^β, 13^CO chemical shifts using the CheSPI algorithm^54^ point to a transient helix, Tom22_PPDE_, highlighted in pink in the primary sequence. **E**, Sequence-based prediction of α-helical content (black) and disorder propensity (blue) in *Sc*Tom22 using AGADIR^55^ and ODiNPred^56^, respectively. **F**, Sequence alignment of the Tom22_PPDE_ reveals conserved hydrophobic and negatively charged residues. *Sc: Saccharomyces cerevisiae, Nc: Neurospora crassa, Ce: Caenorhabditis elegans, Dm: Drosophila melanogaster, Hs: Homo sapiens*. The helical wheel (left, *Sc*Tom22) shows the amphipathic nature of Tom22_PPDE_.

Tom20 and Tom70 have distinct preprotein specificities^37^: Tom20 primarily interacts with presequence-carrying preproteins. A binding groove of its globular cytosolic domain recognizes amphipathic, positively charged cleavable presequences, predominantly in an α-helical conformation^31, 40^. Tom70 is responsible for the import of hydrophobic, noncleavable preproteins with internal targeting signals like α-helical mitochondrial solute carriers of the inner membrane (IM), as well as multi-span OM proteins^10, 14, 32, 41–45^. Tom70 performs a dual role: it anchors preprotein-loaded chaperones (Hsp70 and Hsp90)^14, 44, 46^ and binds preproteins (primarily membrane proteins) at internal sequences^32, 45, 47^. No structural data of Tom70 bound to preproteins is available, however, structures in which the SARS-CoV-2 protein Orf9b is tightly bound to Tom70 point to the putative interaction pocket^48, 49^.

Unlike Tom20 and Tom70, Tom22 is essential for yeast viability under respiration conditions^35, 36, 38^, and has often been described as the central TOM receptor^36, 37, 50–52^. Its cytosolic domain (Tom22_cyt_) is hypothesized to interact with preproteins or facilitate their transfer to the import pore^37, 50–53^. However, previous pull-down experiments showed that while purified Tom22_cyt_ binds presequences at low salt concentrations, its interactions with internal targeting sequences appear weak^23, 37, 51^, leaving its precise mechanisms and functional relevance enigmatic. Furthermore, given that Tom22 interacts with Tom20 and Tom70^29, 51^, it is unclear whether Tom22 is an obligatory binding partner for preproteins or whether the receptors function synergistically when handling preproteins^38^.

Mechanistic understanding of precursor transfer requires atomic-level characterization, yet current structural information leaves key aspects unresolved. Cryo-EM structures have not consistently resolved the cytosolic receptors^18–27, 50^:

Tom70 was absent in all structures, an observation ascribed to its loose association with the TOM_core_. In most structures, neither the cytosolic part of Tom22 nor Tom20 was modeled, and in those structures where parts of Tom20 and Tom22_cyt_ were modeled^23–26^, the positions and conformations of both Tom20 and Tom22 differ largely (Fig. S1). It is unclear whether this structural variability points to different functional states, or whether the chemical cross-linkers used in one set of studies^23, 24^ explains the differences. Besides the discrepancies among these structures, the inter-atomic distances between Tom20 and Tom22 in these cryo-EM models are incompatible with in vivo crosslinking results^29^ (Fig. S1B,H).

A critical unresolved mechanistic question is how cytosolic chaperones and receptor domains handle and target preproteins before they enter the translocation pore. Like other protein transport, chaperone and receptor systems, the mitochondrial protein import machinery has to navigate a delicate balance between sufficient binding affinity and efficient preprotein release. While the two primary import receptors Tom20 and Tom70 are required for the import of virtually all mitochondrial precursor proteins, it is not understood how preproteins are released from them.

To uncover how preproteins are handled at the import gate, we performed integrated studies using atomic-level NMR, single-molecule tracking (SMT) in live cells, and yeast mutagenesis. Our results reveal that the cytosolic domain Tom22_cyt_ is intrinsically disordered, contradicting its role as a preprotein-binding receptor. A conserved transient helical element within this disordered protein competitively displaces precursor proteins from Tom20 and Tom70, thereby driving efficient translocation across the outer mitochondrial membrane. We refer to this element as Precursor Protein Displacement Element (Tom22_PPDE_). Our data point to a general protein-release mechanism that underlies the handover of client proteins at import machineries, which is based on a dynamic protein displacement mechanism.

## Results

### A short transient α-helix of the otherwise intrinsically disordered Tom22_cyt_ is essential for function

The removal of the first 74 residues of *S. cerevisiae* (*Sc*) Tom22 has previously been shown to have dramatic effects on preprotein import in yeast^51^. To obtain more precise information on the part of Tom22_1-74_ responsible for these defects, we analyzed the growth of yeast strains that express mutant forms of Tom22. While deletion of residues 2-54 (Tom22_Δ2-54_) caused only minor yeast growth defects on non-fermentable glycerol (YPG) medium, removing just a few more residues (Tom22_Δ2-58_) resulted in strong growth defects (Fig. 1B, Fig. S2A,B). Our data challenge the structural significance of the compact three-helix bundle proposed in recent cryo-EM models of chemically cross-linked samples^23^, as this motif is disrupted by the Δ2-54 truncation (Fig. S1A). To further assess whether the proposed three-helix bundle structure^23^ is stable, we subjected it to all-atom molecular dynamics (MD) simulations and found that it disintegrated within tens of nanoseconds, further questioning the functional relevance of this fold (Fig. S1C,D). In contrast, the cytosolic domain of Tom20_cyt_ (*Homo sapiens*) remains stable over a 0.8 µs long simulation (Fig. S1E, F), as expected for a folded protein domain.

We used NMR spectroscopy to experimentally characterize the structural properties of Tom22_cyt_. NMR spectra of constructs comprising the cytosolic domain of Tom22 (Tom22_1-74_ or Tom22_1-97_; the TM helix starts approximately at residue 97^18^) displayed the characteristic clustering of ^1^H-^15^N cross peaks within a narrow ^1^H frequency range, a hallmark of intrinsically disordered proteins (IDPs) (Fig. 1C). Quantitative analysis of backbone chemical shifts and spin-relaxation parameters demonstrated that the majority of Tom22_cyt_ exists in unfolded, extended conformations (Fig. 1D, Fig. S3A). Importantly, however, within this intrinsically disordered Tom22_cyt_, residues E55 to D68 transiently adopt an amphipathic α-helical conformation, at an estimated population of about 20-30% (Fig. 1D-F). The disordered nature of Tom22_cyt_ as well as the presence of a transient helical element appear to be a general feature, as revealed by NMR analysis of Tom22 from *Rattus norvegicus* (*Rn*) and *Yarrowia lipolytica*, sequence-based predictions, and AlphaFold2 modeling^57^ (Fig. S3B-I). Additionally, previous reports point to disorder in *Arabidopsis thaliana* Tom22^58^.

Collectively, NMR data on Tom22_cyt_ from three organisms, sequence-based structure predictions, and atomistic simulations revealed that Tom22_cyt_ is not a stably folded domain, but largely disordered with a functionally important short linear motif (SLiM)^59^ that adopts α-helical conformations ~20-30% of the time. Mutagenesis of the Tom22_PPDE_ segment is not compatible with growth on non-fermentable glycerol medium, demonstrating that this conserved ~14 residue short element is essential for Tom22 function.

### Electrostatic binding of preproteins by Tom22 is not relevant under physiological conditions

To reconcile the disordered nature revealed here with the previous reports suggesting direct binding of Tom22_cyt_ to preproteins^23, 37, 52^, we studied in vitro interactions of Tom22 with a 69-residue long presequence of the ATP synthase subunit 9 from *N. crassa* (pSu9_1-69_), a well-established model presequence^60–62^. Detailed NMR titration studies revealed that at low salt concentration (20 mM KCl) Tom22_cyt_ and pSu9_1-69_ interact with a dissociation constant in the single-digit micromolar range (Fig. 2A and Fig. S4A-F), mirroring previous reports from pull-down experiments^23^. Remarkably, both proteins retain their disordered nature within this dynamic complex, as revealed by their NMR chemical shifts. The interaction is driven by electrostatic charge complementarity, involving a negatively charged sequence stretch N-terminal to Tom22_PPDE_ (residues 35-53) and positive charges adjacent to the presequence of pSu9_1-69_ (Fig. 2B and Fig. S4G, H), similar to other charge-driven complexes of intrinsically disordered proteins^63^.

**Figure 2.**
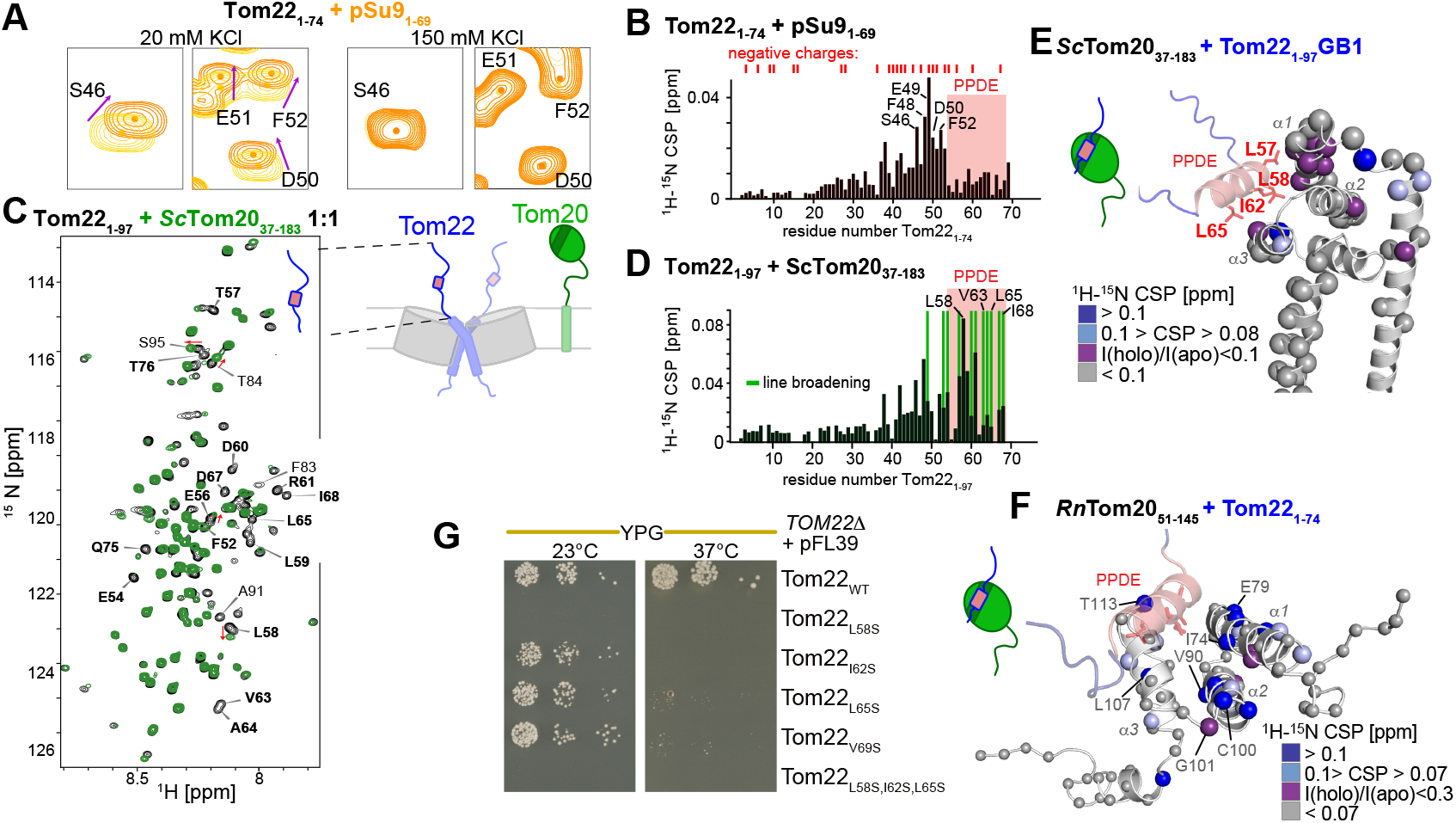
Tom22_cyt_ binds a presequence through electrostatics only at low salt, but binds the presequence-binding cleft of the Tom20 receptor at physiological conditions via its PPDE element. **A**, Interaction of *Sc*Tom22_1-74_ with pSu9_1-69_ probed by ^1^H-^15^N correlation spectra (yellow, orange), at 20 and 150 mM KCl. (No other salt concentrations were tested.) See Fig. S4B,C and E for more spectra and titration curves. **B**, Chemical-shift perturbation in Tom22_1-74_ induced by GB1-pSu9_1-69_ at 20 mM KCl (see spectra in panel A and Fig. S4A). Residues with largest effects, and the position of negatively charged residues (top) are indicated. **C**, ^1^H-^15^N correlation spectra of *Sc*Tom22_1-97_ apo (black) and in the presence of stoichiometric amount of *Sc*Tom20_37-183_ (green). Residues that show significant perturbations are annotated. **D**, Chemical-shift perturbation and line broadening effects in Tom22_1-97_ by *Sc*Tom20 (see spectra in panel C). Example residues with strong CSP or line-broadening effects are indicated. **E**, AlphaFold3 structural model of *Sc*Tom20 in complex with short cytosolic fragment of Tom22_45-72_. Spheres are shown at all Tom20 residues for which experimental ^1^H-^15^N chemical shift perturbation (CSP, i.e. peak-position changes) data in *Sc*Tom20_37-183_ induced by the addition of *Sc*Tom22_1-97_ (1:2) are available; the colors indicate the extent of the perturbation by Tom22. **F**, Chemical-shift perturbation and line broadening effects in *Rn*Tom20_51-145_ upon addition of *Sc*Tom22_1-74_ (molar ratio 1:2.2), plotted on an AlphaFold model. See Fig. S7C for details. **G**, Single point mutations affecting conserved hydrophobic residues from Tom22_PPDE_ cause severe yeast growth defects on non-fermentable/respiratory (YPG) medium.

At physiological salinity, however, the Tom22_cyt_–pSu9_1-69_ interaction is abolished, as revealed by the absence of detectable NMR binding signatures (Fig. 2A, Fig. S4C). We performed further experiments with three presequence peptides, an internal matrix-targeting signal, and a β-barrel targeting signal sequence at 100 mM KCl and 50 mM KP_i_ concentration and found that Tom22_cyt_ does not bind any of these targeting sequences with detectable affinity (Figs. S4J and S5H). In contrast, and consistently with its established role as a receptor protein, the Tom20_cyt_ binds all these presequences at the canonical presequence binding cleft (Figs. S4M,N and S6E-H).

The nature of the charge-stabilized intrinsically disordered complex of Tom22_cyt_ with a presequence explains why this interaction is salt-sensitive^37, 52^. The long-range electrostatic interactions required for this complex formation are quenched at physiological conditions, thereby suppressing this complex formation. Consistent with this finding, the deletion of the negatively charged Tom22_cyt_ region identified as the binding region does not result in a growth defect (Fig. 1B), in line with a previous report^64^. These findings show that direct preprotein binding is not the primary role of Tom22.

### The transient helix Tom22_PPDE_ binds to the presequence binding site of the import receptor Tom20

Previous reports used co-immunoprecipitation^65, 66^ and crosslinking in cells^29^ to establish that Tom20 and Tom22 interact, but structural details have remained controversial^24, 26^ (Fig. S1). NMR interaction experiments of Tom20_cyt_ and Tom22_cyt_ allowed us to identify the binding site accurately: chemical shift perturbations (CSPs), line broadening and spin-relaxation experiments revealed that the interaction requires the Tom22_PPDE_, which becomes more α-helical when binding Tom20, pointing to a conformational-selection mechanism (Fig. 2C,D and Figs. S6 and S7).

NMR experiments that detect Tom20_cyt_ showed that the Tom22_PPDE_ binds to the conserved hydrophobic preproteinbinding cleft formed by helices α1 and α3^31, 40^ in both *Rn*Tom20_cyt_ and *Sc*Tom20_cyt_ (Figs. 2E,F and Fig. S6D-H). We probed the binding of six different presequences to both *Rn*Tom20_cyt_ and *Sc*Tom20_cyt_ by NMR, and confirmed that Tom22_PPDE_ and presequences bind at the same binding site (Figs. S6 and S7). Independent experiments with rat Tom20 (*Rn*) that quantified CSPs and peak intensity changes established that the Tom22_PPDE_ binding site is conserved across species (Fig. 2F and S7C).

Collectively, this data shows that Tom22_PPDE_ and Tom20 interact via the hydrophobic presequence-binding site. We validated the functional importance of this hydrophobic interface by yeast growth assays: substituting conserved hydrophobic residues of Tom22_PPDE_ (L58, I62, L65) with a hydrophilic one (Ser) severely impaired or even abolished growth on respiratory medium (Fig. 2G; Figs. S8). This finding underscores the essential role of this interaction surface in mitochondrial function.

### Tom22_PPDE_ functions as a preprotein displacement element

The finding that Tom22_PPDE_ binds to the preprotein-binding site on Tom20 suggests that Tom22 is functionally positioned to displace precursor proteins. That such a displacement mechanism is indeed operational is supported by the binding affinities determined with the soluble domains: the *Sc*Tom20_cyt_-*Sc*Tom22_cyt_ complex (dissociation constant, K_d_ ~1.5 µM) is more tightly formed than complexes of *Sc*Tom20_cyt_ with presequences (e.g. K_d_ ~7-25 µM for pSu9_1-69_, at pH 6.5; Table S1). Because both Tom20 and Tom22 are membrane anchored, their effective affinity on the native membrane is expected to be even higher than the one determined in solution^67^. We investigated the interactions formed in the simultaneous presence of a presequence (pSu9_1-69_), Tom20 and Tom22 (Fig. S9). These experiments point to competition of Tom22 and the presequence for binding to Tom20, with fast binding and release, as expected. Interestingly, however, they also indicate the presence of ternary complexes, revealed by the fact that the bound-state population of Tom22 is increased when adding the presequence. As shown above, the pSu9-Tom22 interaction is electrostatic in nature. Long-range electrostatic interactions are quenched at high salt concentration, and therefore the binary pSu9-Tom22 has weak affinity; however, short-range electrostatic interactions are known to persist at high salt concentrations^68^. While we did not obtain data that unambiguously establish the role of charge interactions, we speculate that displacement of the presequence from Tom20 by Tom22_PPDE_ may occur through the concerted action of the hydrophobic Tom20–Tom22 interaction and the electro-static Tom22–preprotein contacts formed within this transient ternary complex.

### The transient helix Tom22_PPDE_ also binds the preprotein binding site of the second import receptor Tom70

We hypothesized that Tom22_PPDE_ may also interact with Tom70, i.e. that the preprotein displacement mechanism may operate similarly with this second receptor protein. Because Tom70_cyt_ is large (molecular mass >60 kDa) and difficult to study by conventional NMR, we utilized methyl-TROSY spectroscopy^69^ in conjunction with selective methyl labeling of methionine, isoleucine, and threonine (MIT) residues in a deuterated background. We obtained sequence-specific assignments of the methyl groups using 55 single-point mutants, in which we mutated, one by one, the Met, Ile or Thr residues, which opens the possibility to probe interactions and dynamics of Tom70 in a residue-resolved manner. To date, no structural data of preprotein–Tom70 complexes have been reported. To directly probe the interactions of different preproteins with Tom70, we prepared stable complexes of Tom70_39-617_ with hydrophobic carrier fragments (Aac2 TM helix 4 and PiC TM helix 6) with two approaches, using a pull-down method or cleavable fusion constructs (see Methods). The resulting complexes were stable for several weeks at concentrations exceeding 150 µM, confirming Tom70’s function as a holdase chaperone of these aggregation-prone inner-membrane proteins (Fig. S10A,B).

We additionally investigated binding of a different class of preproteins using a fragment of the outer-membrane voltage-dependent anion channel (VDAC_257–279_). Changes in Tom70 methyl-TROSY NMR spectra induced by these preprotein clients (Aac, VDAC) revealed a continuous binding surface on Tom70, primarily within the conserved hydrophobic groove (Fig. 3A-C and Figs. S10 and S11). This groove is in good agreement with information obtained earlier by cross linking^47^, and the binding site we identified overlaps with the interaction site of the SARS-CoV-2 protein Orf9b^48, 49^ (Fig. S10F). Interestingly, the cyclic form of the VDAC peptide which forms a β-turn, binds more efficiently than the more flexible linear form. This can be rationalized by the fact that the β-turn forming version exhibits a well defined hydrophobic side^41^, more favorable to interact with the Tom70 binding groove (Fig. 3C and Fig. S11B).

**Figure 3.**
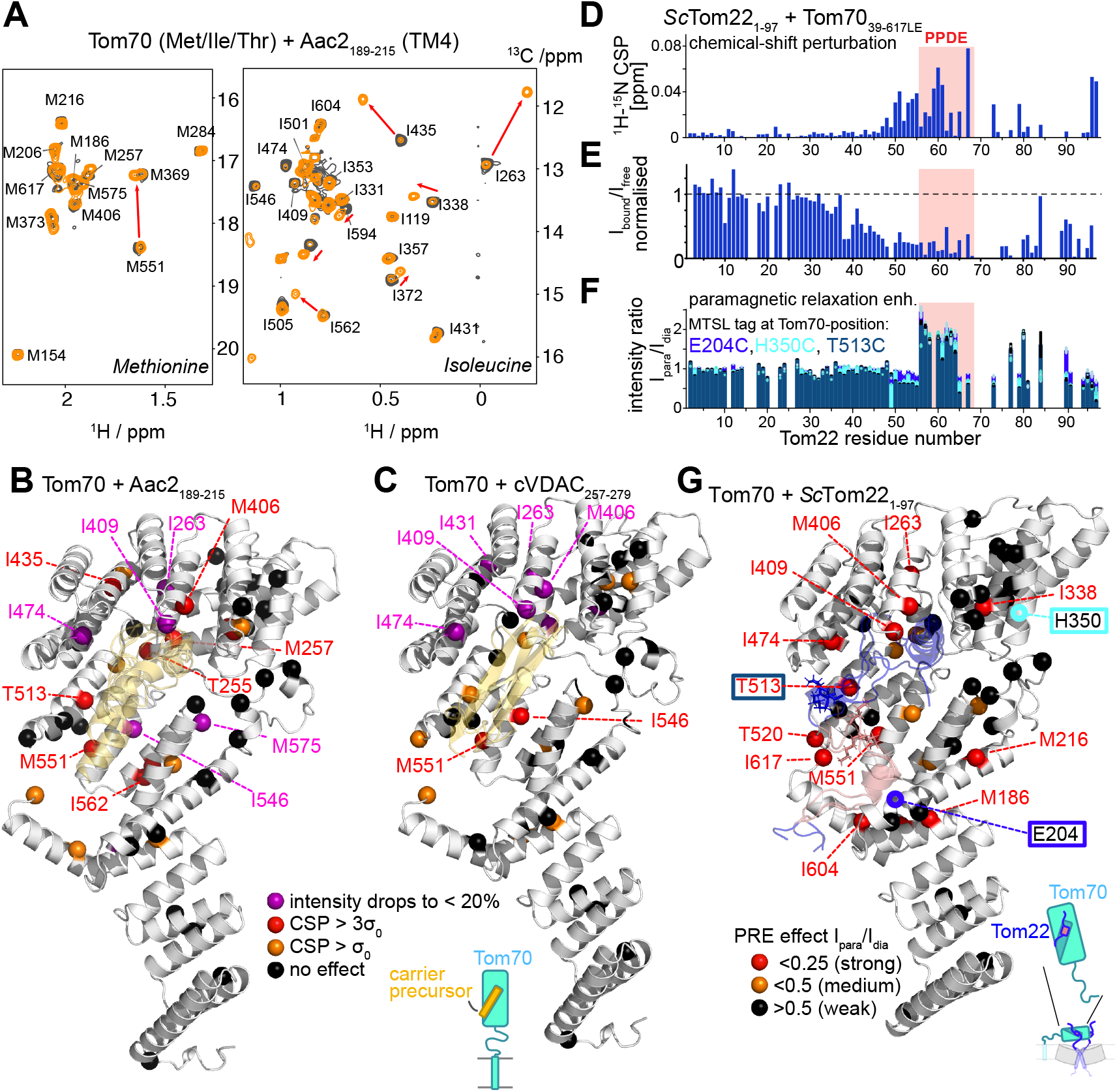
The Tom22_PPDE_ element engages the preprotein-binding cleft of Tom70. **A**, ^1^H-^13^C spectra of [^15^N,^2^H],Met-#-[^13^CH_3_],Ile-∃1-[^13^CH_3_],Thr-%2-[^13^CH_3_]-labeled (henceforth: MIT-labeled) Tom70_39-617_ (apo, black) and of the MIT-labeled Tom70_39-617_–Aac2_189-215_ sample (after TEV cleavage, orange). Methyl resonance assignments are indicated. Signals with particularly large CSP due to Aac2_189-215_ are indicated. See Figs. S10 and S11 for additional spectra and analyses. **B**, CSP and broadening effects due to Aac2_189-215_ plotted onto an AlphaFold model of the Tom70_39-617_–Aac2_189-215_ complex. **C**, CSP and broadening effects due to cVDAC_257-279_ plotted onto a complex structure obtained using the docking software HADDOCK and the NMR interaction restraints. Three binding poses of cVDAC, obtained from HADDOCK, are shown. **D**, ^1^H-^15^N CSP in Tom22_cyt_ upon addition of Tom70_cyt_. **E**, Intensity change of amide signals in Tom22_1-97_ upon addition of unlabeled Tom70. The reduced intensity of the PPDE and adjacent parts is ascribed to the slow tumbling of the large Tom70 which leads to rapid signal decay. **F**, Paramagnetic effects in Tom22_1-97_ upon addition of Tom70 with an MTSL label at either position 204, 350 or 513 (also indicated on the structure of panel G). Plotted is the ratio of ^1^H-^15^N peak intensities in *Sc*Tom22_1-97_ in samples containing PRE-labeled Tom70_39-617_ over intensities in the diamagnetic Tom22-Tom70 sample. A short repetition delay (0.5 s) was used, therefore generating intensity increases for residues close to the paramagnetic site due to the enhanced longitudinal relaxation. **G**, Tom70-Tom22 structural model from molecular-dynamics simulations. The colored spheres indicate the PRE effects detected for Tom70 methyl groups in the presence of Tom22_1-74_ tagged paramagnetically on lysine residues. Additionally, the positions of the MTSL labels (204, 350, 513) of the PRE experiment shown in panel F are indicated on the structure.

We then investigated how Tom22_cyt_ binds the Tom70 receptor. A suite of experiments monitored binding either by detecting Tom22’s backbone amides or the 51 Met, Ile, Thr methyl groups of Tom70 for which data were obtained (Figure S11C). These data unambiguously demonstrated that Tom22 binds to the substrate-binding groove of Tom70 identified above (Fig. 3D-F and Fig. S10, S11). On the side of Tom70, this was observed most prominently in experiments with MIT-labeled Tom70 bound to paramagnetically tagged Tom22 (Fig. S10D, S11C(iii)). Additionally, Tom22-induced CSPs, and intensity changes upon binding the (nonparamagnetic) Tom22 points to the same binding site (Fig. S11C(i,ii), and Fig. 3G).

Observing Tom22, CSPs induced by Tom70 clearly identify the PPDE as interaction motif. Additionally, strong reduction of Tom22 NMR signals (without PRE tagging) was observed for the PPDE as well as parts adjacent to it (Fig. 3E); this is typical of the binding of a highly flexible protein to a large partner^70^, and suggests that additional parts around the Tom22_PPDE_ get slowed down in their dynamics, either by direct binding to Tom70 or through their attachment to the bound PPDE. Furthermore, paramagnetic relaxation enhancement (PRE)^71^ experiments of Tom22 in the presence of Tom70 carrying a PRE tag at three different sites also highlights that the interaction chiefly involves the Tom22_PPDE_ (Fig. 3D-F).

We built structural models of the Tom70_cyt_-Tom22_cyt_ complex using AlphaFold2 and subjected these models to molecular-dynamics (MD) simulations to gain some insights into their stability. AlphaFold2 reported two distinct sets of conformations, with a different orientation of the PPDE in Tom70’s binding pocket. Both binding poses are stable throughout the 0.5 µs simulation, suggesting that both represent plausible conformations of this complex. We used the Tom70-detected PRE data to explore if one of these conformations is in better agreement with experiments (Figure S12). One of these two conformations is in better, although not perfect, agreement with the PRE data, and a representative conformation is shown in Fig. 3G. We note that this is not an experimental atomic-level structure. However, the functional mechanism we propose does not rely on the exact binding pose but only on the fact that the PPDE binds the preprotein-binding pocket. The fact that the CSPs induced by Tom22 are smaller than those induced by Aac2 (Figure S11A,C) suggests that Tom22 is more dynamically bound than Aac2.

Taken together, these results demonstrate that Tom22 binds primarily via Tom22_PPDE_ to the preprotein-binding groove not only of Tom20, but also of Tom70, implying that a similar preprotein displacement mechanism is operational for the translocation of the vast majority of mitochondrial precursor proteins with a presequence or an internal targeting signal.

### Tom22_PPDE_ is crucial to recruit Tom20 to the TOM complex in living cells

The Tom20 and Tom70 receptors are not permanently bound to TOM_core_. Instead, bound and free species are thought to coexist, which has been shown directly for the case of Tom20 ^39^. Which parts of TOM_core_ are involved in recruiting the receptors is under debate. We reasoned that the interaction of Tom22_PPDE_ with the cytosolic domains, demonstrated above, must contribute to the attachment of these receptors to the TOM_core_. Our model implies that if the Tom22_PPDE_–receptor interaction is weakened, the equilibrium would be shifted towards the freely diffusing species. This hypothesis can be tested in vivo in intact mitochondria by quantifying the fraction of TOM_core_-associated versus freely diffusing receptor domains in cells that harbor wild-type Tom22 or variants with reduced binding affinity. To weaken this interaction, we designed hydrophobic-to-charged mutants in human (*Hs*) Tom22_PPDE_ (L44E, L48E and L48E,L51E) which correspond to the residues L54, L58 and L61 in baker’s yeast (Fig. 4A). NMR experiments confirmed that mutations at these hydrophobic positions in *Sc*Tom22, either to charged residues, or to the helix-breaking Pro, reduced the complex population to ~50% of the WT level (K_d_ 1.5 µM for WT); the estimated change in K_d_ is around two orders of magnitude (Fig. 4B, C).

**Figure 4.**
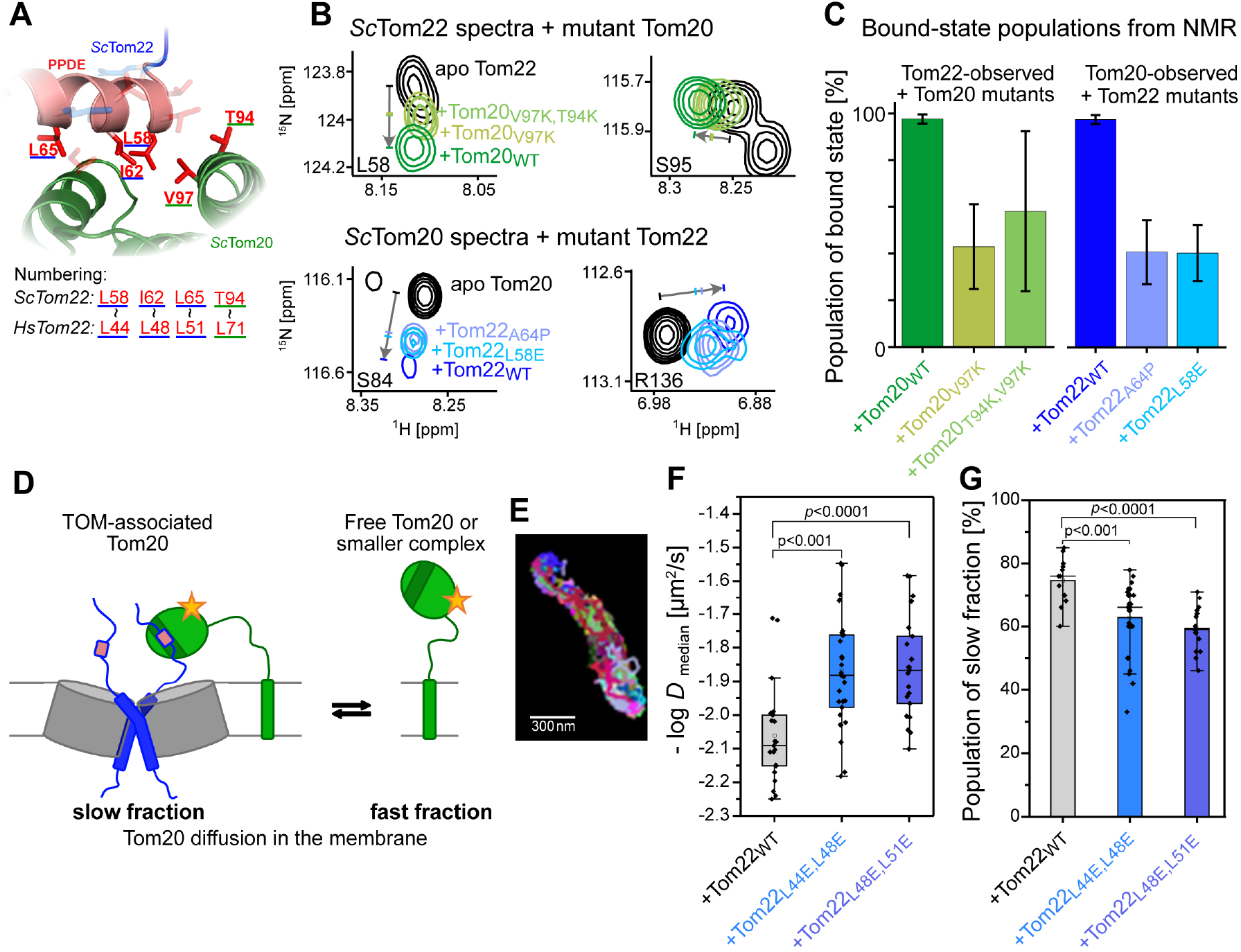
The Tom22_PPDE_-Tom20 interaction is important for binding Tom20 to the TOM complex in living cells. **A**, Zoomed in view of the *Sc*Tom20-*Sc*Tom22 binding site (AlphaFold model), highlighting important residues also identified in the NMR CSP data. **B**, Examples of NMR interaction spectra of *Sc*Tom22_1-97_ with *Sc*Tom20. Spectra of isotope-labeled Tom22 (top) and Tom20 (bottom) are shown in interaction with unlabeled binding partners. The isotope-labeled protein was in each case at a concentration of 60 µM, and the unlabeled protein at 120 µM. Experiments were performed at pH 6.5, 50 mM KPi buffer, 20 °C. All *Sc*Tom20_37-183_ : *Sc*Tom22_1-97_ and *Sc*Tom22_1-97_ : *Sc*Tom20_37-183_ samples shown are at 1:2 molar ratios. (See Figs. S7E and S5L for full spectra of Tom20 and Tom22, respectively.) **C**, Populations of the bound state of Tom22 (left) and Tom20 (right), obtained from the interaction experiments of panel B. The population in the case of WT interaction partner was calculated from K_d_ = 1.5 µM. We estimated the population of the bound state by comparing the CSP in the experiments with the mutant and WT interaction partners. The average over the five residues with the largest reduction of CSP was used. **D**, Schematic representation of the equilibrium between a TOM-bound state of Tom20, which diffuses slowly, and a free more rapidly diffusing species. Single particle tracking (SPT) experiments allow for diffusion analysis of fluorescence-labeled Tom20 (asterisk). **E**, Representative mitochondrion in the periphery of a HeLa cell, showing the cumulative trajectories (4000 frames). See Fig. S13 for details of the SPT workflow. **F**, Median diffusion coefficients (−log *D*) obtained from SPT experiments. **G**, Population of the slowly-diffusing Tom20 fraction, analyzed by fitting distinct Gaussians to the probability density function (PDF) of diffusion coefficients, *D*, of Tom20 in cells expressing different Tom22 mutants, illustrated in Fig. S13F. Statistics (F-G): Mann-Whitney-Test. Box-and-whisker plots, box: 25-75 percentile; open square indicates mean, line indicates median. Analysis of protein expression levels (Figure S13G) shows that approximately 85% of the Tom22 incorporated in the TOM complex is the tagged version, suggesting that a full incorporation may result in an even more pronounced change in the diffusion coefficient.

To investigate how interactions between the cytosolic domains influence Tom20 association with TOM_core_, we performed Single Particle Tracking (SPT) microscopy on fluorescently tagged Tom20 in live HeLa cells (Fig. 4D, E and Fig. S13). Previous SPT analyses established that there are two populations of Tom20 with different mobility in mitochondria: a slowly-diffusing TOM_core_-bound state and a rapidly-diffusing state that corresponds to unbound Tom20^39^. We found that in HeLa cells expressing Tom22 variants with reduced Tom20-binding affinity, Tom20 exhibited enhanced mobility, evidenced by a significantly increased mean-square displacement (Fig. S13I) and a shift in diffusion coefficients towards higher values (Fig. 4F, and Fig. S13J). Statistical analysis confirmed a decrease in the population of slowly-diffusing Tom20 species when a Tom22_PPDE_ mutant with lower binding affinity is expressed (Fig. 4G). These findings demonstrate that Tom22-Tom20 binding stability is critically driven by the hydrophobic properties of Tom22_PPDE_, contradicting previous models of electrostatic binding interfaces^26^ (Fig. S1G, H). Our single molecule tracking results show that the hydrophobic interaction surface of Tom22_PPDE_ is crucial to enhance the interaction between the cytosolic domains of the presequence receptor Tom20 and Tom22, and thus for recruiting the primary receptor to the TOM_core_ complex in vivo.

### Tom22_PPDE_ is crucial for mitochondrial biogenesis

To further demonstrate the in vivo requirement of the Tom22_PPDE_, we generated yeast strains expressing Tom22 variants with a TEV protease Cleavage Site (TCS) inserted either immediately N-terminal or C-terminal to the PPDE (Tom22_1-53_TCS_54-152_ or Tom22_1-74_TCS_75-152_, respectively). These strains also expressed cytosolic TEV protease under the control of the pGAL promoter. Induction of TEV protease by addition of 0.25% galactose in the growth medium efficiently cleaved the N-termini of the Tom22-TCS-variants, in contrast to WT Tom22 and the Tom22-TCS-variants on medium without galactose (Fig. 5A). Removal of the N-terminus including the PPDE caused a severe growth defect on YPG in contrast to the N-terminus removal before the PPDE which grew similar to WT (Fig. 5B). This indicates that the Tom22_PPDE_ is specifically essential to maintain respiratory yeast growth similar to full-length Tom22^36^. Moreover, we selected three Tom22 single point mutants Tom22_I62S_, Tom22_L65S_, Tom22_V69S_ (Fig. 2G) and three Tom22ΔN mutants Tom22_Δ2-54_ (minor phenotype), Tom22_Δ2-56_ (medium phenotype) and Tom22_Δ2-58_ (strong phenotype on YPG – Fig. 1B) and cultured these yeast strains to check for in vivo import defects via the accumulation of mitochondrial precursor protein in the cytosol^72^. The precursor of Mdj1 (pMdj1) clearly accumulates more in the Tom22_I62S_ point mutant compared to wild type (Fig. 5C). In addition, we observed an increasing level of precursor Mdj1 accumulation from Tom22_Δ2-54_ to Tom22_Δ2-56_ to Tom22_Δ2-58_ (Fig. 5C), which directly correlates with the Tom22ΔN-mutant growth phenotypes (Fig. 1B). This indicates that the Tom22_PPDE_ is crucial to maintain mitochondrial precursor protein import. Given the large flexibility of Tom22_cyt_, we reasoned that the precise attachment site of the precursor-protein displacement element may not be crucial, as long as the displacement occurs in the vicinity of the Tom40 import pore. We therefore hypothesized that a TOM_core_ complex in which the Tom22_PPDE_ element is attached at a different location, rather than to the Tom22_TM_ helix, may still be functional. To test this hypothesis, we first generated a yeast strain expressing Tom22 lacking its cytosolic domain (Tom22_Δ2-75_). This strain exhibited a growth defect comparable to a Tom22 deletion strain (Fig. 5D), confirming the essential role of the cytosolic domain for Tom22 function. We then engineered a strain in which the first 75 residues of Tom22_cyt_ were removed and instead fused to Tom7 (Tom22_1-75_-Tom7, Tom22_76-152_) (Fig. 5E, F). This attachment of the Tom22 cytosolic domain to Tom7 rescued the growth phenotype (Fig. 5D; Tom22_1-75_-Tom7 + Tom22_76-152_). In contrast, selective removal of the Tom22_PPDE_ element from this construct (Tom22_1-52_-Tom7 + Tom22_76-152_) again led to severe growth defects, highlighting the functional importance of the Tom22_PPDE_.

**Figure 5.**
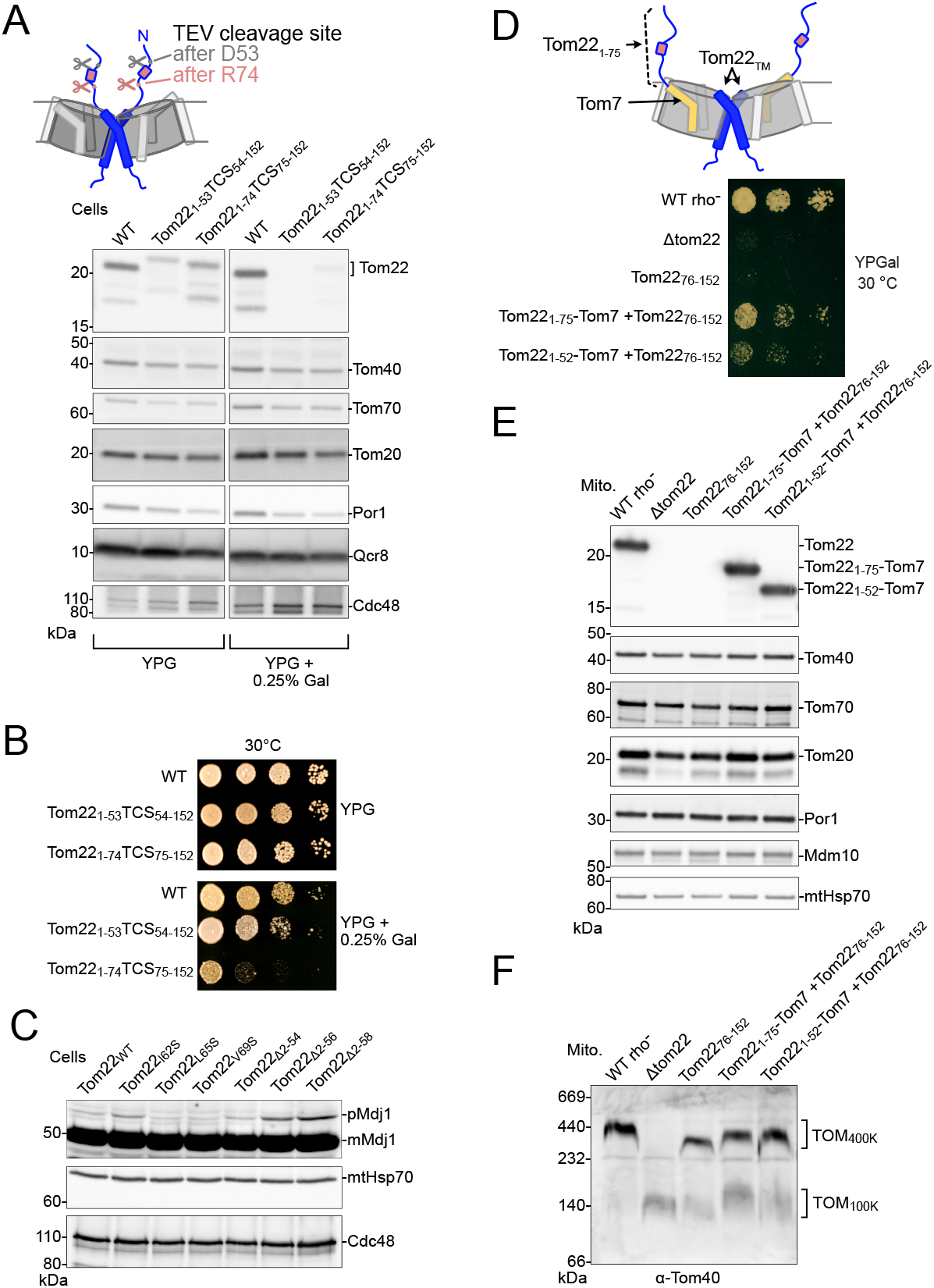
Importance of the Tom22_PPDE_ for mitochondrial biogenesis. **A**, Top, scheme of TEV protease cleavage sites (TCS) introduced N- and C-terminal of the Tom22_PPDE_. Bottom, whole cell extract/immunodecoration. Upon addition of 0.25% galactose (Gal) to the medium the N-termini of both Tom22_1-53_TCS_54-152_ and Tom22_1-74_TCS_75-152_ are efficiently cleaved, while the other Tom proteins remain unaffected. **B**, In contrast to Tom22_1-53_TCS_54-152_ and WT, Tom22_1-74_TCS_75-152_ has a severe growth defect on respiratory glycerol (YPG) medium upon TEV protease induction. **C**, In vivo accumulation of Mdj1 precursor protein (pMdj1) in Tom22_PPDE_-mutant whole cell extracts. **D**, Top, scheme of yeast strain with N-terminal truncation of Tom22_76-152_ lacking the PPDE expressing a fusion construct of Tom22_1-75_-Tom7. Bottom, fusion of Tom22_1-75_-Tom7 containing the Tom22_PPDE_ rescues the yeast growth defect of Tom22_76-152_ lacking the PPDE. **E**, Whole cell extract/immunodecoration reveals a similar expression of Tom22_cyt_-Tom7 fusions compared to WT Tom22. **F**, Blue native-PAGE and immunodecoration of isolated mitochondria expressing Tom22_cyt_-Tom7 fusions show a comparable assembly of the TOM_400K_ complex.

Collectively, the severe growth defect upon in vivo cleavage of the Tom22_PPDE_ (Figs. 5A, B), the in vivo accumulation of mitochondrial precursor protein in Tom22_PPDE_-mutants (Fig. 5C), and the domain swapping rescue experiments (Figs. 5D–F) demonstrate that the precursor protein displacement element of Tom22 is crucial for mitochondrial function.

### Tom22_PPDE_-bound import receptors dynamically sample the space above the translocation pore

Having shown the highly dynamic nature of the cytosolic domain of Tom22 and the importance of Tom22_PPDE_ in receptor binding and preprotein displacement, we integrated all available data into a structural model, and used MD simulations to explore its properties. We generated an explicit all-atom model of *Sc*TOM_core_ based on the cryo-EM structure^18^, added the cytosolic region of Tom22 using a disordered-ensemble generator^73^, embedded the complex in a lipid bilayer with explicit water and performed a 1 µs MD simulation. In this simulation, the cytosolic domain of Tom22 samples a broad conformational landscape (Fig. 6A), with both Tom22_PPDE_ elements able to essentially reach the entire space above the Tom40 translocation pores.

**Figure 6.**
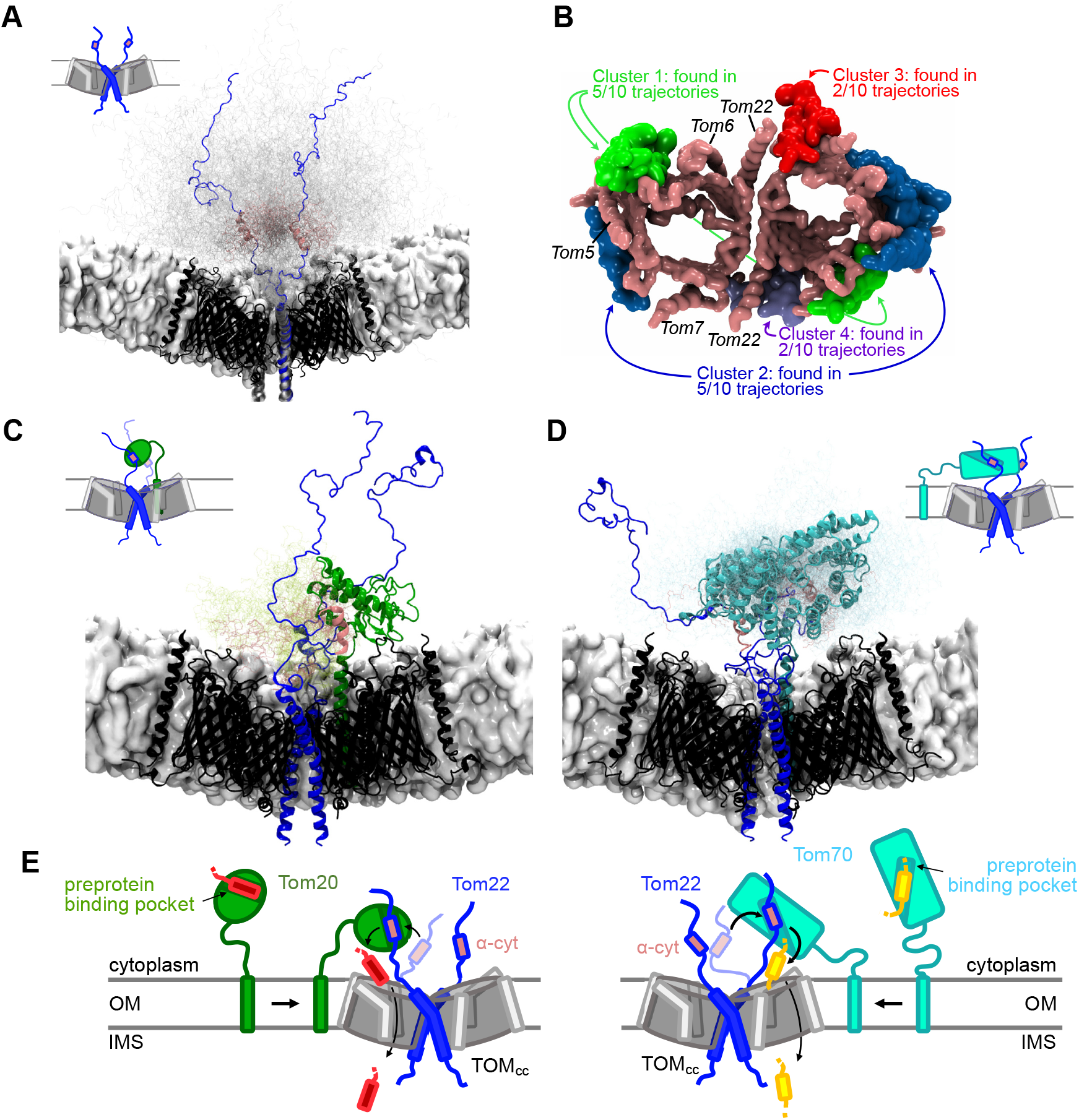
Dynamic model of the receptor domains at the TOM gate. **A**, Dynamic amplitude available to Tom20 from all-atom MD simulations of the entire TOM_core_ and ensemble generation (see Methods). Two chains of Tom22 are shown in blue and pink (for Tom22_PPDE_), and other conformations of Tom22 are shown in grey. **B**, The positions of the Tom20 TM helix found in coarse-grained MD simulations cluster into 4 groups. Symmetry-equivalent positions have the same color. The position of the Tom20 TM helix in a cryo-EM model obtained with chemical cross-linkers^23, 24^ corresponds to cluster 4. Ten simulations were performed, starting from various initial locations (Fig. S1K). **C**, A plausible conformation of Tom20 bound to the Tom22_PPDE_ helix, positioned above the translocation pore and attached to its TM helix in a position corresponding to cluster 1. 99 additional conformations of the folded domain of Tom20 and of Tom22_PPDE_ are shown as sticks. See Supplementary movie 1. **D**, A plausible conformation of Tom70, bound to Tom22_PPDE_, above the translocation pore, with its TM helix in a position corresponding to cluster 1 of panel B. See Supplementary movie 2. **E**, Schematic model of precursor protein translocation, assisted by Tom22_PPDE_, acting as a preprotein displacement element. A preprotein (red, yellow) bound to Tom20 (left) or Tom70 (right) is displaced by Tom22_PPDE_ (pink) above the TOM translocation pore, positioning it for translocation.

The positions of the Tom20 TM helix differ largely in available TOM cryo-EM structures^24, 25^ (Fig. S1). Given this controversy, we investigated the possible positions using coarse-grained MD simulations of freely diffusing Tom20 transmembrane helix in the membrane. Amongst multiple simulations corresponding to 80 different trajectories of Tom20_TM_, we found multiple binding events at different sites of TOM_core_. Stable association events clustered into four distinct binding sites at the TOM_core_ (Fig. 6B). The finding of multiple possible binding sites mirrors the structural variability observed in cryo-EM structures and suggests that the overall Tom20–TOM_core_ association is not dictated by specific contacts between the transmembrane segments. This finding implies that the affinity between the cytosolic domains is crucial for the receptor association, and our SPT experiments have highlighted that Tom22_PPDE_ plays the crucial role in this recruitment.

Using one of these bound Tom20_TM_ positions, we built a TOM_core_-Tom20 model in an explicit membrane and solvent, and subjected it to an all-atom MD simulation. The Tom20_cyt_-Tom22_PPDE_ complex connected to the two respective TM helices by their long flexible linkers sampled a wide range of conformations above the TOM pore (Fig. 6C and Supplementary Movie 1). The Tom20_TM_ segment can adopt multiple positions, but the cytosolic receptor domain of Tom20 is consistently located above the TOM translocation pore in all these conformations. This localization enables a direct preprotein transfer into one of the two Tom40 translocation pores. While being very mobile overall, the folded Tom20 domain occasionally formed transient contacts with the rim of the Tom40 barrel (see Supplementary Movie 1).

Using the same strategy, we constructed a model of the Tom22_PPDE_-Tom70 complex (starting with the AlphaFold/MD-derived model shown in Fig. 3G), linked to the TM helices, and performed all-atom MD simulations (Fig. 6D, Supplementary Movie 2). Tom70 remains above the Tom40 pore, without making substantial contacts to Tom40 or the small Toms, in positions that allow a direct preprotein transfer into the respective TOM translocation pores.

The structural modeling, together with the yeast experiments, demonstrates that the essential precursor-protein displacement function of Tom22 does not depend on a specific architecture, but only requires that the precursor protein displacement element is positioned dynamically above the Tom40 translocation pores.

## Discussion

### Tom22 ensures controlled preprotein release by a precursor-protein displacement mechanism

Efficient protein transfer in chaperone and import systems requires a fine balance: interactions must be stable to prevent premature release, yet dynamic enough to enable timely handover. Our data reveal that Tom22 employs a sophisticated mechanism to promote preprotein release specifically above the Tom40 translocation pore. Rather than acting primarily as a preprotein-binding receptor domain, Tom22 uses a transient α-helical molecular recognition element, Tom22_PPDE_, to compete with incoming precursors for their binding sites on Tom20 and Tom70. NMR experiments show that Tom22 engages the precursor-binding regions of both receptors, consistent with in vivo cross-linking results^29^; our data suggest that this mechanism is operational in a similar manner on both Tom20 and Tom70. We propose a mechanism in which Tom22_PPDE_ engages the hydrophobic interaction site on the receptor, possibly progressively, thereby displacing the preprotein (Fig. 6E). We hypothesize that during this process negatively charged residues of Tom22 may interact with the positively charged face of the presequence to facilitate the release. Electrostatic interactions may also bias the direction of precursor movement and prevent backward diffusion toward the cytosol^64^. Supporting such an electrostatically driven engagement, our in vitro data show that pSu9 forms a highly dynamic, charge-mediated complex with Tom22. It should be noted, however, that previous reports in which negative charges of Tom22 were deleted suggested that these negative charges are not essential^64^, and the present study did not find direct evidence for the role of the electrostatics in a physiological setting.

Spatially controlled displacement of preproteins is of central importance along the entire preprotein translocation pathway through each step of mitochondrial protein import. While this study has focused on the central step at the mitochondrial protein import gate, the same principle is likely operational in other parts of the import pathway. In the case described here the displacement element (Tom22_PPDE_) and the holdase function (Tom20, Tom70) are on different proteins; this arrangement ensures spatially controlled release right at the import gate. A functionally similar displacement mechanism may, however, also occur when the binding site and displacement element are on the same protein, such as in holdase chaperones^74, 75^.

### The Tom22_PPDE_ increases the local concentration of receptor domains at the translocation pore

In addition to its role in displacing precursor proteins at the translocation pore, binding of Tom22_PPDE_ to Tom20 and Tom70 increases the local receptor concentration at the TOM_core_ complex (Fig. 4). We propose that receptor enrichment in the vicinity of TOM_core_ facilitates efficient handover of precursor proteins from cytosolic chaperones to the translocation pore. For efficient preprotein relay, a well-adjusted affinity of the binding of Tom22_PPDE_ to the preprotein-binding sites of the receptors is crucial: if the binding was too strong, the receptors would essentially be blocked. The affinity we measured (K_d_ in the micromolar range) implies continuous binding and release. Overall, this process leads to a higher probability of finding the receptors close to TOM_core_, but their preprotein-binding sites are not permanently blocked.

Our yeast experiments challenge the significance of a recently proposed Tom20-Tom22 binding interface^24, 26^ (Fig. S1A,G). More than one interaction mode may exist between Tom22_PPDE_ and the receptor domains; we speculate that one Tom22 may bind Tom20 close to the membrane, as found in two recent cryo-EM structures^26, 27^, while the second Tom22_PPDE_ acts in displacing a preprotein. The two Tom22_cyt_ domains may thereby act synergistically in receptor recruitment and preprotein release.

### Tom22 as a general interaction platform on the TOM complex

Beyond its interactions with Tom20 and Tom70, Tom22_cyt_ likely serves a broader function as a versatile interaction platform for additional components of the import machinery. The assembly of the TOM–SAM supercomplex depends on contacts between Tom22_cyt_ and Sam37^76^. Interestingly, the N-terminal part of Tom22, up to residue 54, is dispensable for this interaction, whereas residues surrounding the Tom22_PPDE_ can be cross-linked to Sam37^77^. This suggests that Tom22_PPDE_ contributes directly to Sam37 recognition and binding. Furthermore, Tom22_cyt_ has been implicated in binding the Hsp40 co-chaperone Xdj1^15^, with the client-binding cleft of Xdj1 contributing to this interaction. Although the molecular details of this interface remain unresolved, we speculate that Tom22 may promote the release of client proteins from Xdj1, analogously to the preprotein-displacement mechanism described in this study. Our data highlight that intrinsically disordered soluble domains, often invisible in cryo-EM structures, can play key functional roles in membrane-protein machineries.^78^

Taken together, our findings reconcile previously conflicting structural interpretations of Tom22 and establish its role as a highly dynamic interaction hub of the TOM complex coordinating not only co-chaperones and translocase supercomplex formation, but also import receptor recruitment and precursor protein release. Key to its function is the precursor protein displacement element of Tom22. While we have not shown all aspects of the mechanism for both yeast and human, the conservation of the PPDE suggests that the mechanism is present across eukaryotes. It has previously been reported that human Tom22 can complement the absence of yeast Tom22^79^, further supporting the idea of this conserved mechanism. Our experiments, using yeast and human proteins and live cells explain how the Tom22_PPDE_, a conserved ~14 residue segment with α-helical propensity within the intrinsically disordered cytosolic domain, can be essential for Tom22 function to drive mitochondrial protein import.

## STAR⋆METHODS

Detailed methods are provided in the online version of this paper and include the following:

- KEY RESOURCES TABLE
- METHODS DETAILS

## SUPPLEMENTAL INFORMATION

Supplemental information includes 13 figures, one table and two movies.

## Data availability

Backbone chemical-shift assignments of *Sc*Tom22_1-74_, GB1-pSu9_1-69_, *Sc*Tom20 and MIT methyl assignments of *Sc*Tom70 have been deposited in the Biological Magnetic Resonance Databank (https://bmrb.io) under the accession numbers 51381, 51093, 53318, and 54075, respectively.

## Acknowledgements

This research was supported by the Scientific Service Units (SSU) of Institute of Science and Technology Austria (ISTA) through resources provided by the Nuclear Magnetic Resonance Facility and the Lab Support Facility. Part of this work has been performed at the platforms at the Grenoble Instruct-ERIC Center (ISBG; UMS 3518 CNRS CEA-UGA-EMBL) with support from the French Infrastructure for Integrated Structural Biology (FRISBI; ANR-10-INSB-05-02) and GRAL, a project of the University Grenoble Alpes graduate school (Ecoles Universitaires de Recherche) CBH-EUR-GS (ANR-17-EURE-0003) within the Grenoble Partnership for Structural Biology. We thank Petra Rovo, Margarita Valhondo, Adrien Favier and Alicia Vallet for excellent NMR support and Olena Tsvyetkova for assistance with protein production. This research was funded in whole or in part by the Austrian Science Fund (FWF) 10.55776/PAT1647625 and 10.55776/I6223 to P.S and the French Agence Nationale de la Recherche (ANR-18-CE92-0032) to P.S., and the German Deutsche Forschungsgemein-schaft, through project CRC944 (INST190/1672) to K.B.B., SFB 1381 project ID 403222702 to N.W., and WI4506/1-1 project ID 406757425 to N. W. We acknowledge support by Germany’s Excellence Strategy (CIBSS EXC-2189 project ID 390939984) to N.W. This study made use of NMRbox: National Center for Biomolecular NMR Data Processing and Analysis, a Biomedical Technology Research Resource (BTRR), which is supported by NIH grant P41GM111135 (NIGMS).

## Author contributions

Conceptualization: I.S., P.S., J. S., U. G., N. Wi.; methodology, P.S., F.D., N.Wi., K.B.B.; investigation, I.S., U.G., J.S., I.G., N.We., A.K., C.S., F.S., A.F., R.S., L.F.F., B.B., J.M.D., F.D., P.S.; writing – original draft, I.S., P.S.; Writing - Review & Editing, I.S., U.G., J.S., K.B.B., N.Wi., F.D., P.S.; visualization, I.S., U.G., J.S., K.B.B., I.G., C.S., L.F.F., J.M.D., K.B.B., F.D., P.S.; resources, D.R.; supervision, P.S., N.Wi., K.B.B.

## Methods

### Protein production and purification for in vitro experiments

#### Protein Constructs and their Cloning

The gene coding for the cytosolic domain, residues 39-617, of *S. cerevisiae* Tom70 protein (Uniprot ID P07213) was inserted into a bacterial expression vector pET-28a(+) carrying kanamycin resistance. Cloning of the gene of interest between NdeI and XhoI restriction sites yields a protein with an N-terminal His_6_-tag followed by a TEV protease cleavage site. After TEV cleavage, the protein sequence comprised the residues GHM at the N-terminus, followed by residues 39 to 617 (i.e. GHMGKK) and two residues, LE, at the C-terminus (i.e. GLMLE). This construct was used as starting point for the mutants used for assignment, and for the Tom22-detected experiment of Figure 3D-F. For the interaction experiments with Tom22 (Fig. 3G, S10D, S11C) a construct with the native C-terminus (devoid of the LE) was used (reference 10G1 in the key resource table). As a reference for the interaction experiment with Aac2 (Figure 3B, Figure S10B, S11A) a construct with a TEV-His6 at the C-terminus was used, and the C-terminus was cleaved before NMR (reference 11C7).

Similarly, genes encoding the *Sc*Tom22 protein constructs (residues 1-97 and residues 1-74, Uniprot ID P49334) were inserted into pET47b-TEV-GB1 expression vector. Here, Tom22 proteins are tagged at the C-terminus with the GB1 and His_6_-affinity tag, and the TEV protease cleaving site was inserted to allow the tag removal. The *Sc*Tom22_1-97_ protein sequence was followed by the TEV cleavage tag (LENLYFQ), MVELTEIKDDVVQLDEPQFSRNQAIVEEKASATNNDVVDDEDDSDSDFED EFDENETLLDRIVALKDIVPPGKRQTISNFFGFTSSFVRNAFTKSGNLENLYFQ. The rationale for the two constructs, comprising residues 1-74 and 1-97, respectively, is the following. Residue 97 is the last residue assigned to the cytoplasmic domain, according to UniProt ID P49334, making *Sc*Tom22_1-97_ a natural choice. Construct *Sc*Tom22_1-74_, which has much higher production yield, comprises the PPDE (which ends approximately at residue 69); it has a higher yield after purification than *Sc*Tom22_1-97_. NMR comparisons of both constructs indicated that the differences in terms of (residual) structure are minute. The cytosolic domain of *Rattus norvegicus* Tom20 (Uniprot ID Q62760) comprising residues 51-145 was chosen based on previous structural biology studies^31^. The construct comprised a hexahistidine tag, followed by a GB1 sequence and a TEV cleavage site, N-terminal to the Tom20_51-145_ sequence.

The *Sc*Tom20 (Uniprot ID P35180) cytosolic domain constructs were designed based on multiple sequence alignment of Tom20 proteins, to ensure retention of the conserved residues within the cytosolic domain. The *Sc*Tom20_37-183_ begins after the N-terminal conserved transmembrane helix (9-28) and after the helix breaker (Pro).The gene coding for the cytosolic domain of yeast Tom20_37-183_ was inserted into an expression vector pET-GB1-TEV-stop, carrying kanamycin resistance. Tom20_37-183_ protein was preceded by a His_6_-affinity tag, GB1 and a TEV protease cleavage site, akin to the Tom22 sequence. The cleaved final Tom20_37-183_ protein construct comprised 3 additional residues at the N-terminus (GHM) and two additional residues (LE) at the C-terminus due to XhoI restriction site, in addition to the native sequence 37-183 resulting it the sequenceGHMPQFRK VLRQRAKEQAKMEEQAKTHAKEVKLQKVTEFLSMELAKDPIPSDPSEREATFTTNVENGERLSMQQGKELEAASKF YKALTVYPQPADLLGIYQRSIPEAIYEYIILMIAILPPANVASFVKGVVGSKAESDAVAEANDIDDLE.

The mitochondrial subunit 9 of the F0 ATP synthase of *Neurospora crassa* was chosen as a representative of a precursor or client protein. The first 69 residues of this precursor protein carry the mitochondrial matrix targeting sequence, termed pSu9. pSu9 has been extensively used in in vitro mitochondrial import assays^51, 52, 85^. The gene coding for residues 1-69 of pSu9 was cloned in a customized pET-GB1-TEV stop vector for bacterial protein expression, resulting in an N-terminal His_6_-GB1-tag, cleavable by the TEV protease. The pSu9_1-69_ protein sequence with 3 foreign residues (Gly-His) at the N-terminus as remaining after cleavage: GHMASTRVLASRLASQMAASAKVARPAVRVAQVSKRTIQTGSPLQTLKRTQMTSIVNATTRQAFQK RAYSSLE. For data shown in Figure S4, the construct was used without cleavage resulting in a protein with the sequence MG HHHHHHPMKQYKLILNGKTLKGETTTEAVDAATAEKVFKQYANDNGVDGEWTYDDATKTFTVTEGSGSGSENLY FQ prior to the sequence shown in the previous sentence.

The cloning, production, and quality control of the plasmids were performed by GeneCust (Boynes, France; https://genecust.org/).

#### Protein Production

Optimizing recombinant protein production in *Escherichia coli* revealed distinct optimal conditions for each construct. Tom70 protein was overexpressed in *E. coli* BL21(DE3) cells by incubating shaking cultures overnight at 20 °C after induction with 0.75 mM isopropyl β-D-1-thiogalactopyranoside (IPTG). In contrast, the Tom22 protein (both 1-74 and 1-97 constructs) produced better results when overexpressed for 3-4 hours at 37 °C after induction with 0.75 mM IPTG.

The *Sc*Tom20 protein construct was produced in *E. coli* BL21(DE3) cells, while *Rn*Tom20 was produced in BL21(DE3)-RIL+ cells. Shaking cultures for *Sc*Tom20 were incubated for 3-4 hours at 30 °C following induction with 0.75 mM IPTG, whereas *Rn*Tom20 was overexpressed overnight at 20 °C after induction with 0.5 mM IPTG.

Unlabeled proteins were produced in lysogeny broth (LB) growth medium. Isotopically labeled Tom70 proteins used for detection by NMR experiments were expressed in D_2_O M9 minimal medium and specifically labeled on threonine, methionine and isoleucine side chains using TLAM-I^*ω*1^M^*ε*^T^*γ*^ kit from NMR-Bio (https://www.nmr-bio.com) according to the manufacturer’s instructions. Isotopically labeled protein constructs were produced in M9 minimal medium with ^15^NH_4_Cl (1 g/L) and ^13^C D-glucose (2 g/L). The growth medium for deuterated (^2^H) proteins was additionally made in D_2_O.

#### Protein Purification

Bacterial cells expressing Tom70 were harvested at 5500 g and resuspended in buffer **A** (50 mM Tris, 300 mM NaCl, pH 8, 10% glycerol, 5 mM 2-mercaptoethanol (β-ME) with addition of cOmplete - EDTA free protease inhibitor cocktail). Cells were lysed by sonication in an ice-bath and the soluble protein fraction was loaded on a gravity-flow Ni-NTA agarose resin. The column was first washed with 5 column volumes (CV) of buffer **A**, followed by 5 CV of buffer **A** supplemented with 10 mM of imidazole. Tom70 protein was eluted from the Ni-NTA resin with buffer **A** supplemented with 300 mM imidazole. The elution fractions were concentrated and dialyzed for 2 h at 4 °C against buffer containing 50 mM Tris pH 7.6 and 50 mM NaCl. In the second round of dialysis, fresh buffer was supplemented with 2 mM of dithiothreitol (DTT) and 1 mg of TEV protease was added to the dialysis bag per every 20 mg of Tom70. Cleaving reaction was incubated for 16 hours at 4 °C. Cleaved Tom70 protein was recovered in the flow-through fraction from a second Ni-NTA affinity chromatography, while the TEV protease and cleaved His-tag were retained on the resin. Tom70 was further purified by size-exclusion chromatography (SEC) on a HiLoad 26/600 Superdex 200 pg column equilibrated in 20 mM Tris pH 7.6 and 50 mM NaCl.

*Sc* and *Rn*Tom20 protein constructs were purified as described for Tom70. After TEV cleavage, Tom20 proteins were purified on a HiLoad 26/600 Superdex 75 pg column in buffer B containing 20 mM Tris pH 7, 100 mM NaCl buffer.

Tom22 was expressed with a C-terminal His_6_-GB1-tag. After elution from a Ni-NTA gravity flow column, the fusion protein was concentrated and further purified by SEC on a HiLoad 26/200 Superdex 75 pg column in buffer B. The removal of the tag was achieved in the presence of 2 mM DTT and 1 mg of TEV protease per 20 mg of Tom22 at 4 °C for 16 hours. Cleaved Tom22 protein was recovered in the flow-through fraction during the gravity-flow Ni-NTA affinity chromatography. Unlike a previous report^58^, we found that the samples were stable over several weeks, as monitored by NMR and SDS-PAGE.

Protein concentrations were determined from absorption at 280 nm using theoretical extinction coefficients, except for cleaved Tom22 and *Rn*Tom20. For these proteins concentration was determined by the BCA Protein Assay Kit (ThermoFisher Scientific).

GB1-pSu9_1-69_ protein construct was purified from inclusion bodies. The insoluble protein extract after the cell lysis was solubilized in 6 M guanidinium-chloride (Gdn-HCl). After 16 h incubation at 4 °C with agitation, the supernatant was purified on a gravity-flow Ni-NTA column in presence of 4 M Gdn-HCl. Protein was eluted from the affinity column with buffer containing 4 M Gdn-HCl, 50 mM Tris at pH 7.4, 150 mM NaCl and 300 mM imidazole. Purified protein was concentrated to ≤1.7 mg/mL and was subjected to refolding by dialysis in 50 mM Tris pH 7, 150 mM NaCl for 16 hours at 4 °C. Remaining insoluble protein was removed by centrifugation and the supernatant was purified by SEC on a HiLoad 16/600 Superdex 75 pg column in buffer B. After refolding from denaturing conditions, the protein showed instability at low temperatures (−20 °C). For this reason, the refolding was performed just before using the sample.

#### Peptides

All the presequence (matrix targeting sequences of the mitochondrial precursor proteins) peptides were purchased from GeneCust (https://www.genecust.com/). The purity of the peptides is > 95%.

The presequence of the mitochondrial β-subunit of F1-ATPase from *S. cerevisiae* (UniProtKB - P00830) pF1β_1-31_: MVL-PRLYTATSRAAFKAAKQSAPLLSTSWKR.

The presequence of the mitochondrial Cytochrome c oxidase subunit 4 from *S. cerevisiae* (UniProtKB - P04037) pCox4_1-25_: MLSLRQSIRFFKPATRTLCSSRYLL.

The presequence of the rat aldehyde dehydrogenase ALDH2 split in two parts (UniProtID-P11884) pALDH_1-22_: MLRAAL-STARRGPRLSRLLSAA and pALDH_12-22_: PRLSRLLSAA.

The presequence of the subunit 9 of the mitochondrial ATP synthase from *N. crassa* (UniProtKB - P00842) pSu9_1-25_: MASTRVLASRLASQMAASAKVARPA.

The presequence of the β-subunit of the mitochondrial ATP synthase (UniProtKB - P07251) pAtp1_1-35_: MLAR-TAAIRSLSRTLINSTKAARPAAAALASTRRL, and the internal matrix targeting signal of the same precursor protein Atp1_306-324_: DDLSKQAVAYRQLSLLLRR (iMTS-L).

The fragments of human VDAC1 peptide (cyclic or linear VDAC_257-279_) were prepared by solid-phase synthesis as described previously ^41^.

All peptides were received lyophilized. Consequently, they were resolubilized in 100% DMSO and step-wise diluted into interaction-compatible buffer resulting in less than 2.5% DMSO in the final protein sample.

The peptide used for photo-induced cross-linking is a modified version of the peptide used for NMR studies due to amino acid substitution L263 for a Bpa side chain ^41, 86^.

#### Preparation of Tom70-preprotein complexes

##### Pull-down approach

To prepare Tom70–preprotein complexes, we adapted an approach previously proposed for complexes of the same family of preproteins with small TIM chaperones^86^. The His_6_-tagged preproteins (Tim23^87^ or Ggc1^86^) were expressed recombinantly as inclusion bodies, purified under denaturing conditions, and immobilized on a His-affinity resin in the presence of 4 M guanidine-HCl, as described^86, 87^. The column was washed with five column volumes (CV) of buffer A supplemented with 4 M guanidine-HCl and five CVs of buffer to remove the denaturant. Subsequently, a two-fold excess of Tom70_39–617_ was added to the column (passing through the column twice), followed by washing with buffer (10 CVs). The complex was eluted in 5 CVs of buffer A supplemented with 300 mM imidazole, immediately subjected to dialysis and analyzed by SDS-PAGE (Fig. S10A).

##### Cleavable fusion constructs

To generate complexes of Tom70 with fragments of mitochondrial carriers, we inserted the Tom70 sequence fused to either the ADP/ATP carrier Aac2 (TM4) or the phosphate carrier PiC (TM6) between NdeI and XhoI restriction sites of the pET30c(+) vector. The resulting fusion proteins comprised *Sc*Tom70_39-617_, followed either by the linker GPGP**ENLYFQ**↑**G**GVAG (which contains the TEV protease cleavage site (in bold)) and residues 189-215 of Aac2: LYRGFLPSVVGIVVYRGLYFGMYDSLK (Uniprot P18239) carrying a His_6_ tag on its C-terminus, or by the linker GS**ENLYFQ**↑**G**GSGSG and residues 251-299 of PiC: QSTVGLLAQLAKQLGFFGSFAGLPTRLVMVGTLTSLQFGIYGSLKSTLG (Uniprot P23641) tagged C-terminally with a His_6_-tag. The arrow indicates the position of the TEV cleavage. The fusion proteins were expressed and purified as Tom70 (see above). After the first Ni-NTA affinity chromatography, the fusion proteins were subjected to SEC and cleaved for 16 hours at 4 °C by the TEV protease. The cleavage was verified by SDS-PAGE (Fig. S10A).

### Biophysical methods for protein and binding characterization

#### Isothermal Titration Calorimetry experiments

Calorimetric binding experiments with Tom receptors were performed using a MicroCal ITC200 instrument (GE Healthcare). Ten successive 2.5 µl aliquots of titrant were injected into a sample cell containing the analyte. The spacing between the injections was set to 180 seconds and the stirring speed was 750 rpm. All ITC data were acquired in 20 mM Tris pH 7 buffer with 100 mM NaCl at 20 °C. Control experiments included titration of the titrant into the buffer. The enthalpy accompanying each injection was calculated by integrating the resultant exotherm, which corresponds to the released heat as a function of ligand concentration added at each titration point. ITC data were analyzed using the MicroCal Origin software using a single site binding model and non-linear least squares fit of the thermodynamic binding parameters (ΔH, K, and n).

#### Size-exclusion chromatography coupled to Multi-angle Laser Scattering (SEC-MALS)

SEC-MALS experiments were performed on the Biophysical platform (AUC-PAOL) in Grenoble. The experimental setup comprised an HPLC (Shimadzu, Kyoto, Japan) consisting of a degasser DGU-20AD, an LC-20AD pump, an autosampler SIL20-ACHT, a column oven XL-Therm (WynSep, Sainte Foy d’Aigrefeuille, France), a communication interface CBM-20A, a UV-Vis detector SPD-M20A, a static light scattering detector miniDawn Treos (Wyatt, Santa-Barbara, USA), a dynamic light scattering detector DynaPro NANOSTAR, a refractive index detector Optilab rEX. The samples were stored at 4 °C, and a volume of 20, 40, 50 or 90 µl was injected on a Superdex 10/300 200 pg, equilibrated at 4 °C; the buffer was 50 mM Tris, 150 mM NaCl filtered at 0.1 µm, at a flow rate of 0.5 ml/min. Bovine serum albumin was used for calibration. Two independent sets of experiments conducted with two different batches of protein samples were highly similar.

#### NMR spectroscopy

All NMR experiments were performed on Bruker Avance-III spectrometers operating at 600, 700, 800, 850 or 950 MHz ^1^H Larmor frequency. The samples were in NMR buffer (20 mM K-Pi pH 6.5 and 20 mM KCl) with 10% (v/v) D_2_O. All multidimensional NMR data were analyzed with CCPN V3 (Vranken et al., 2005).

#### Resonance assignment experiments

For the resonance assignment of Tom22 and Tom20 proteins from yeast, the following experiments were performed : 2D ^15^N-^1^H-BEST-TROSY, 3D BEST-TROSY HNCO, 3D BEST-TROSY HNcaCO, 3D BEST-TROSY HNCA, 3D BEST-TROSY HNcoCA, 3D BEST-TROSY HNcoCACB, 3D BEST-TROSY HNCACB, 3D BEST-TROSY hNcocaNH and a 3D BEST-TROSY hNcacoNH^88, 89^. The experiments with Tom22 were performed with a 0.6-0.9 mM [^15^N,^13^C]-labeled Tom22, at 298K and pH 6. The Tom20 experiments were performed with 0.5-0.6 mM [^2^H,^15^N,^13^C]-labeled Tom20, at 293K and pH 7. We monitored the integrity of the samples with 2D BEST-TROSY experiments between 3D experiments. Tom22 samples were stable over weeks; in the case of *Sc*Tom20, we found that the sample stability at 500 µM protein concentration was limited to 1-2 days, requiring several samples for the series of assignment spectra. We noted that addition of 50 mM Arg and 50 mM Glu and 200 mM NaCl extends the stability to several days even at 500 µM protein concentration. Additional experiments under these conditions, however, did not extend the amount of assignment further.

Resonance assignments of rat Tom20 (but not of yeast Tom20) had been reported before^31^. We collected BEST-TROSY HN, HNCA, HNcoCA, HNCO and HNcaCO experiments to confirm the sequence-specific resonance assignment of the [^15^N,^13^C]-labeled rat Tom20, aided by previously reported assignment efforts by^31^.

For the resonance assignment of methyl groups of Tom70, 2D ^1^H-^13^C-SOFAST HMQC experiments were recorded for the wild-type and all the 57 mutants in which single Met, Ile or Thr containing amino acids were mutated (see table 1). The experiments were performed with a 0.2-0.8 mM [U-^2^H,U-^12^C],[^13^C,^1^H]-Ile^*δ*1^Met^*ε*^Thr^*γ*^ labeled Tom70, at 298K and pH 7.4. We have assigned 55 methyls (13 Met, 25 Ile, 17 Thr; see BMRB entry 54075). We obtained quantitative analyses for 51 of them (e.g. because partial overlap hampered accurate analysis).

**Table 1.**
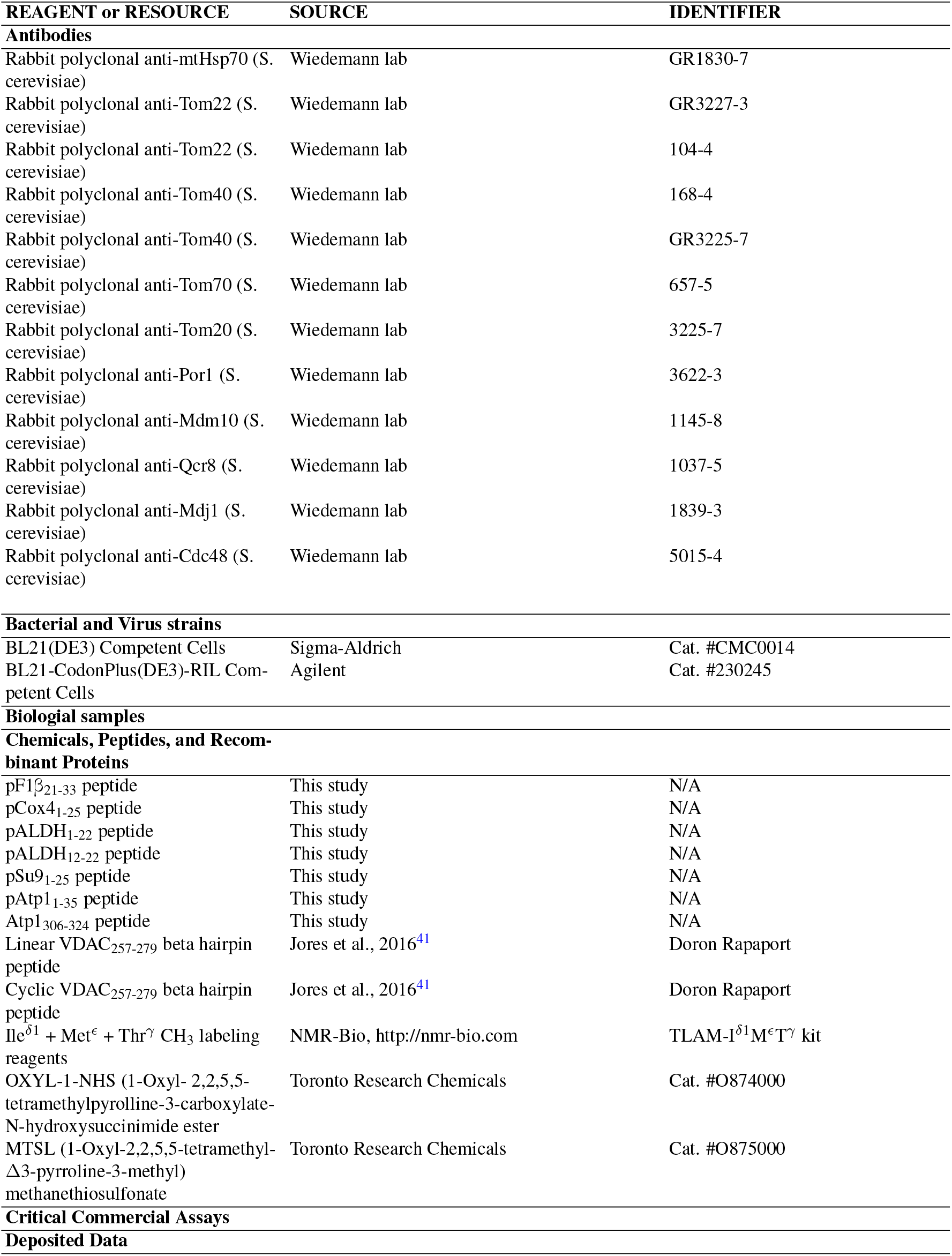
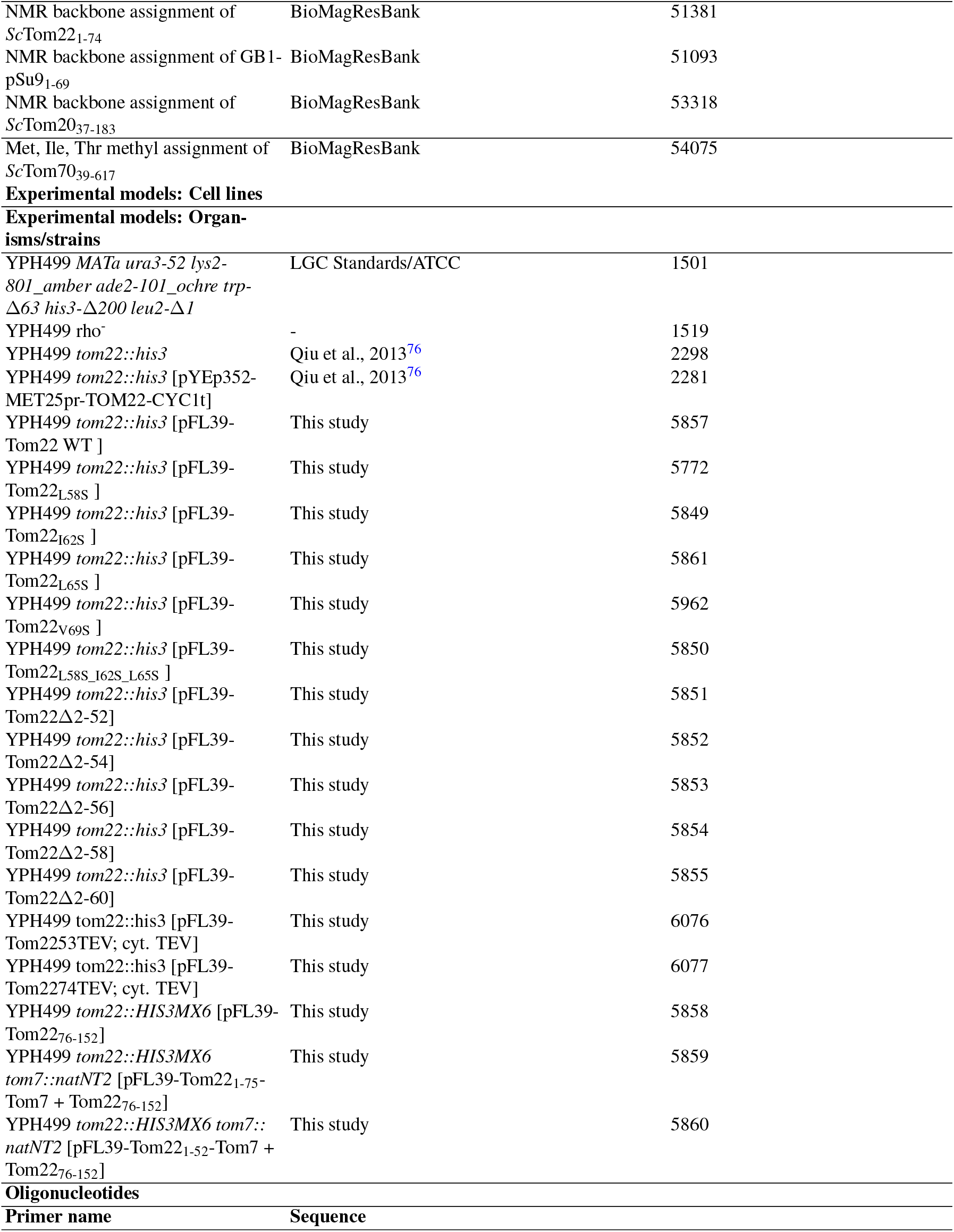
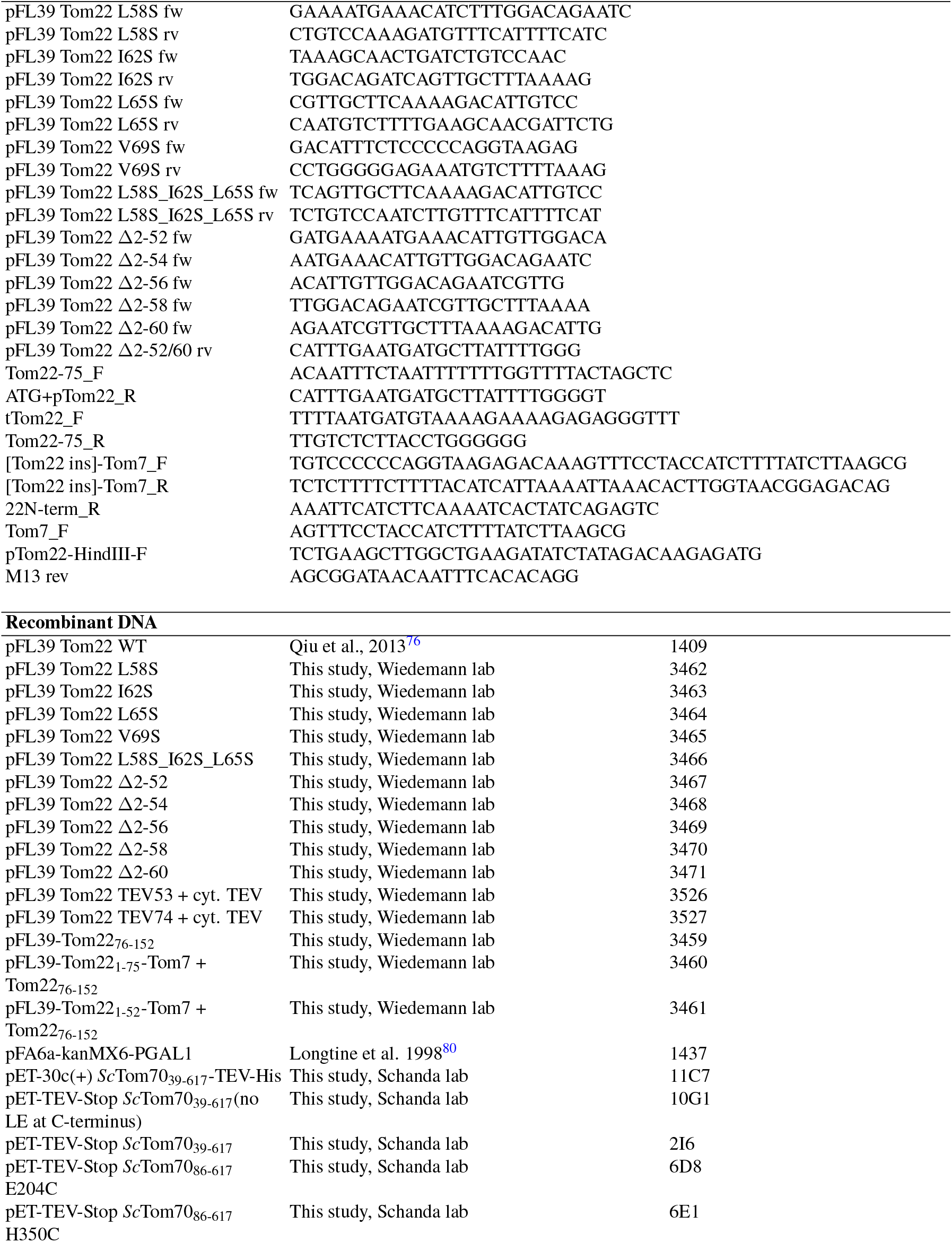
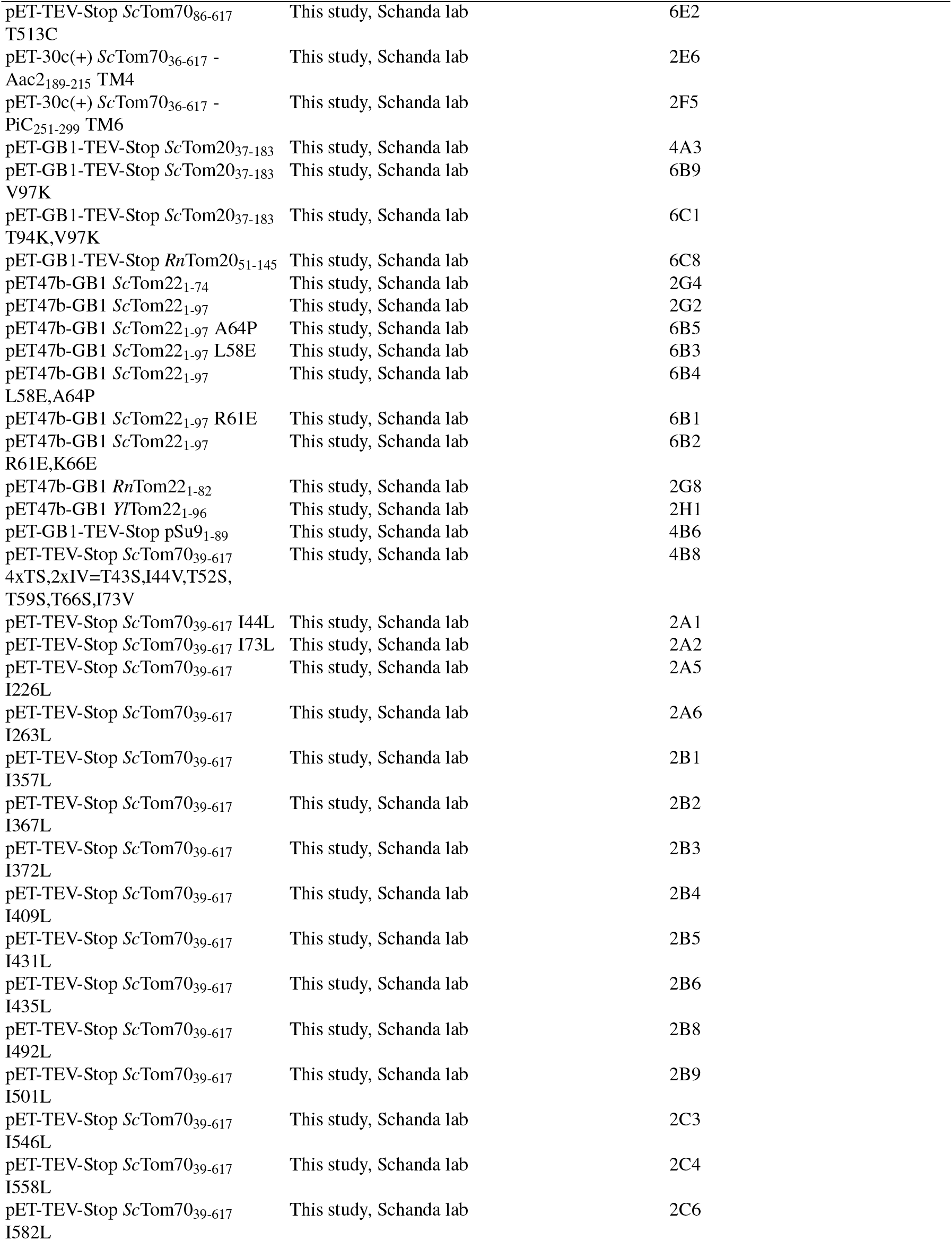
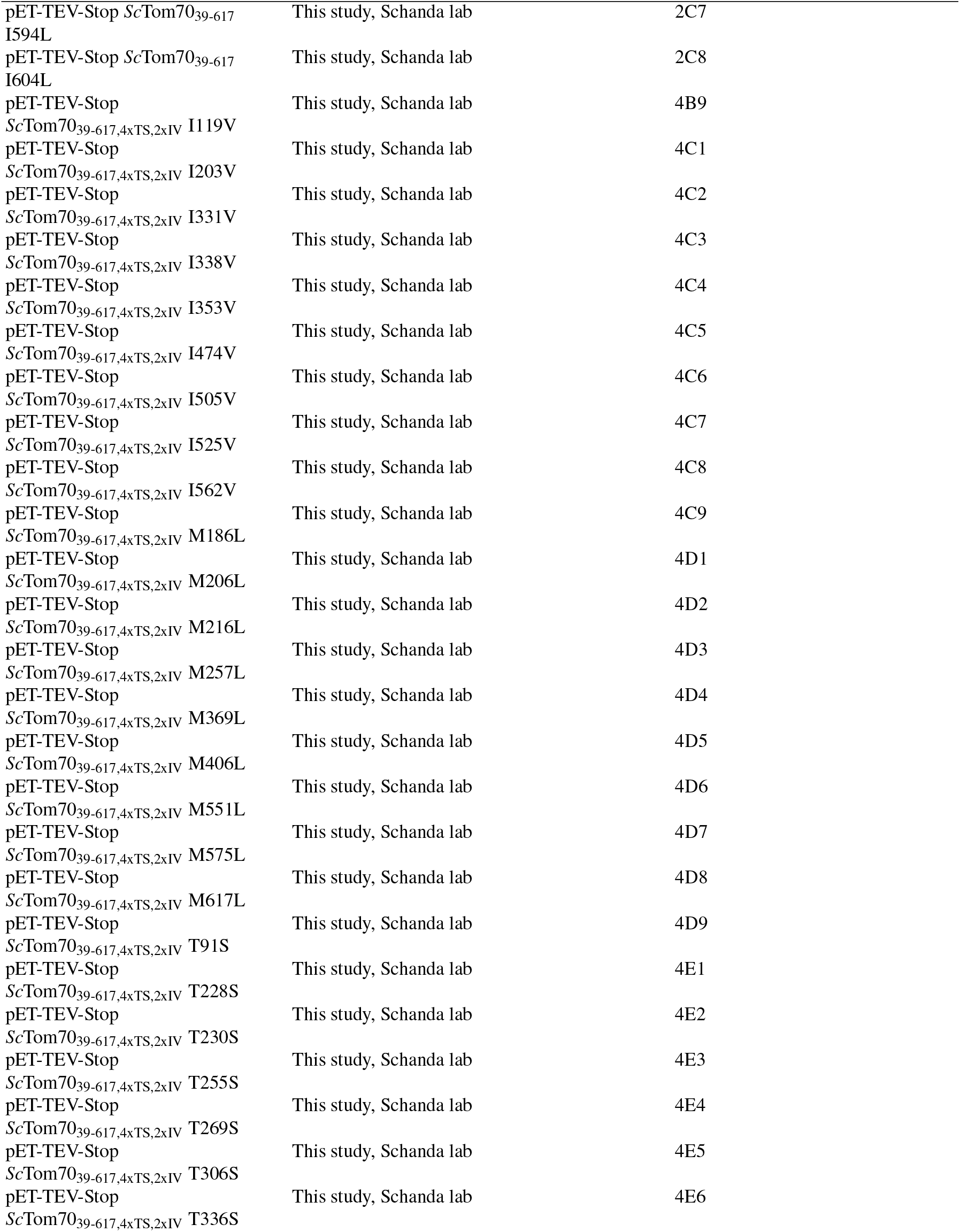
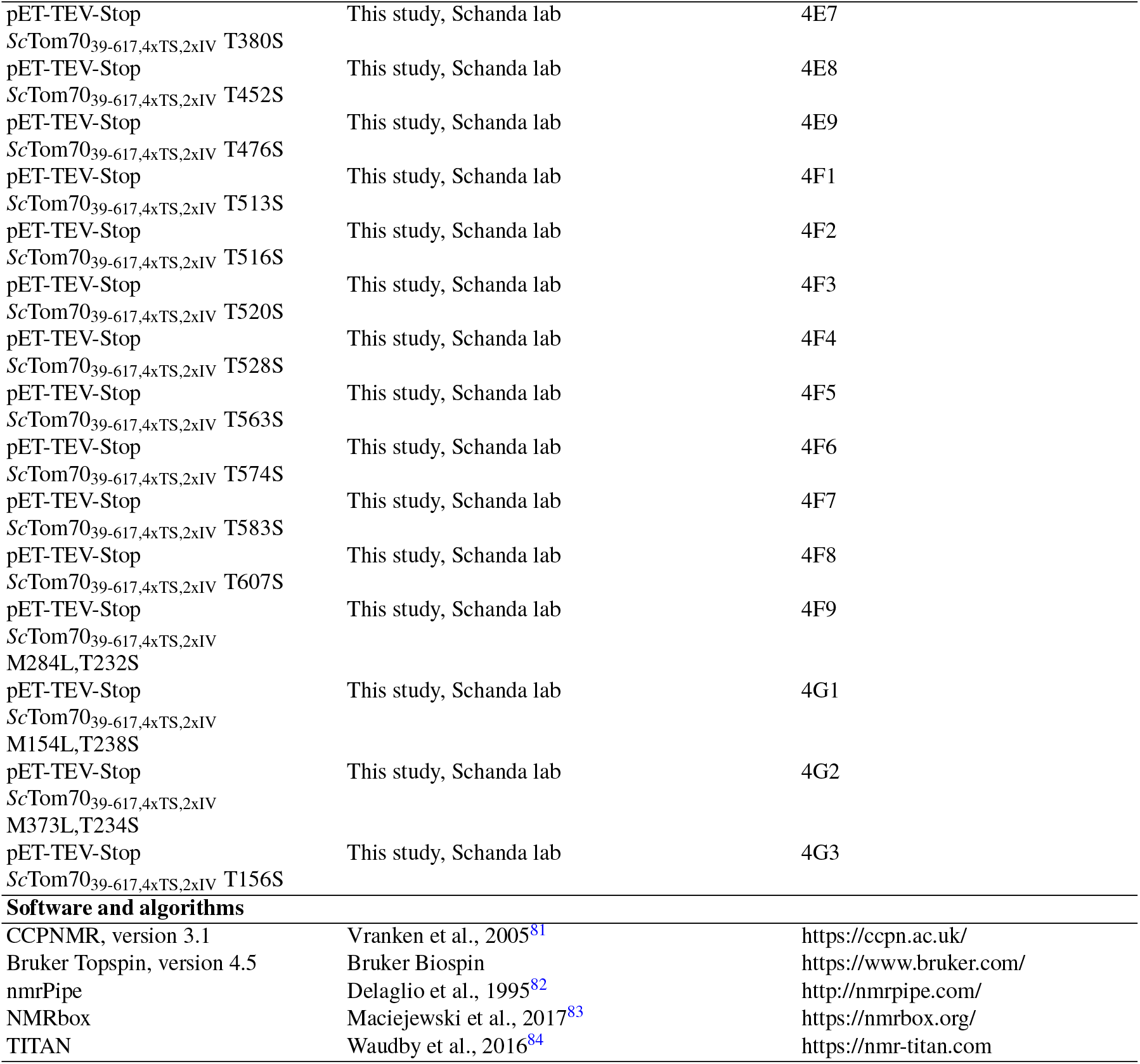
KEY RESOURCES TABLE.

#### Titration experiments with Tom20, Tom22 and the presequences

Separate samples were prepared for each titration point. The peptides were dissolved in pure DMSO and by step-wise addition of NMR buffer (1:1 in each step), the DMSO concentration was reduced to 6%. The NMR buffer contained 20 mM K-Pi pH 6.5 and 20 mM KCl. After adding Tom22 or Tom20, the final DMSO concentration was ≤2.5%, while the Tom20 or Tom22 concentrations were 0.1 and 0.15 mM, respectively. Samples containing recombinantly expressed proteins (rather than peptides) did not contain DMSO. Combined ^1^H-^15^N chemical shift-perturbations (CSPs) were calculated from the weighted difference in the chemical shifts obtained from the ^1^H-^15^N BESTTROSY spectra of the complex samples, in comparison to the apo-receptor spectrum, whereby the contribution of each of the nuclei was weighted by its gyromagnetic ratio.

For calculating the affinities from the NMR titration experiments, the TITAN software package was used (TITration ANalysis; www.nmr-titan.com;^84^, using NMRbox (https://nmrbox.org/)^83^. The 2D HN spectra of the titration series were processed with NMRPipe using the same processing parameters for all spectra. The experimental and processing parameters are read by the TITAN to simulate spectra in the presence of chemical exchange. The TITAN software fits the 2D lineshapes and intensities of selected peaks; errors were estimated with a bootstrap method implemented in TITAN. The fitting was performed assuming a two-state ligand binding model (P + L ⇌ PL) and in each analysis 10-20 residues were selected for the fitting (regions of interest, ROI).

#### NMR relaxation experiments

^15^N R_2_ relaxation experiments for characterization of Tom22_cyt_ backbone protein dynamics, acquired at a ^1^H Larmor frequency of 600 MHz (14.1 T magnetic field strength), consisted of multiple 2D ^15^N-^1^H BEST-TROSY HSQC experiments with variable relaxation delay. Relaxation delays for R_2_ experiments ranged from 0 to 85 ms. HetNOE values, which inform on ps-ns motions, were derived from the ratio of peak integrals in two experiments, with and without ^1^H saturation.

#### Biochemical methods for paramagnetic relaxation enhancement experiments (PRE)

The solvent accessible functional groups of NMR-inactive proteins were labeled with PRE spin labels. For labeling the ε-amino group of lysines, OXYL-1-NHS (1-Oxyl-2,2,5,5-tetramethylpyrolline-3-carboxylate-N-hydroxysuccinimide ester, Toronto Research Chemicals) was used. For sulfhydryl group labeling of cysteines MTSL ((1-Oxyl-2,2,5,5-tetramethyl-Δ3-pyrroline-3-methyl) methanethiosulfonate, Toronto Research Chemicals) was used. Stock solutions of the PRE-label compounds were prepared by dissolving the chemical into 100% DMSO. Using NAP-5 desalting columns, the protein solution was buffer exchanged to labeling buffer (for lysine labeling 10 mM Na_2_CO_3_, for cysteine labeling 50 mM Tris, 50 mM NaCl, pH 7.8). Lysine labelling was chosen for Tom22_1-74_; as this protein contains only four lysines (positions 8, 29, 66, 73). As residues 8 and 29 are not bound to Tom70 (Fig. 3D-F), the Tom70-detected PRE effects (Fig. 3G and Fig. S11C) can be ascribed to arise from paramagnetic tags at positions 66 and/or 73, i.e. within or very close to the PPDE.

For OXYL-1-NHS-lysine labeling of Tom22 (construct 1-74), a 6-fold molar excess of OXYL-1-NHS (3 mM) was added to the protein solution (0.5 mM). In the final reaction 0.8% DMSO was present. The reaction was incubated for 1 hour at room temperature, followed by 4 °C incubation. The excess of spin label was removed by dialysis in 50 mM HEPES pH 7.4, 100 mM NaCl. The lysine labeling was verified by mass spectrometry (166 Da added molecular weight per labeled lysine). For Tom22_1-74_ (4 lysines), a mixture of differently labeled species was observed by mass spectrometry; under typical reaction conditions, the most populated species had three tags, followed by species with 4 labels (ca. 50-60% of the former) and 2 labels (ca. 50%); the species with only 1 label was a minority, and the non-tagged species was barely detectable.

Tom70_39-617_ mutants (C141S/E204C, C141S/H350C and C141S/T513C) were PRE-labeled with MTSL at the incorporated cysteine residues by the addition of a 10-fold excess of spin label compound. After 15 min incubation at room temperature, another 20-fold excess MTSL was added and the mixture incubated at room temperature for 16 h. Excess spin label was removed by dialysis in 20 mM Tris, 100 mM NaCl, pH 7.0. The efficient labeling was verified by mass spectrometry.

#### NMR methods for paramagnetic relaxation enhancement experiments (PRE)

We measured PRE values to provide qualitative information on the interactions between the isotope-labeled protein and the PRE-labeled interaction partner, by comparing NMR signal intensities of samples containing paramagnetic species with those of a sample in which the paramagnetic tags were quenched (reduced), as follows.

For the interaction of ^15^N-^1^H-labeled Tom22_1-97_ with the PRE-labeled Tom70_39-617_ mutants, isotope labeled Tom22_1-97_ was mixed with a 1.5-fold excess of each MTSL-labeled Tom70_39-617_ mutant, respectively, and the peak intensities from the 2D HN-SOFAST NMR experiments compared with the reduced samples as described above (using a relaxation delay of 0.5 s).

For the interaction of OXYL-1-NHS-labeled Tom22_1-74_ with MIT-labeled Tom70_39-617_, methyl-labeled Tom70_39-617_ was mixed with a 2-fold excess of OXYL-1-NHS–labeled Tom22_1-74_ (paramagnetic tagging of lysine groups). The 2D HC-SOFAST HMQC NMR experiments of the oxidized and reduced samples were recorded at 25 °C in 20 mM K-Pi pH 6.5, 20 mM KCl buffer. A short relaxation delay (0.5 s) was used for the Tom22-detected PRE experiments in which Tom70 was paramagnetically tagged at either position 204, 350 or 513; the short recycle delay leads to enhanced intensity due to accelerated longitudinal relaxation in the paramagnetic sample.

### Single-molecule tracking experiments

#### Cell Cultures

HeLa cells were purchased from the Leibniz Institute DSMZ-German Collection of Microorganisms and Cell Cultures. HeLa cells were cultured in T25 flasks and maintained at 5% CO_2_ and 37 °C in supplemented MEM-medium (Minimum essential medium Eagle, Sigma-Aldrich, M2279; 10 % FBS supreme, PAN BioTech P30-3031; 1% HEPES, Sigma-Aldrich H0887-100ML, Ala-Gln, Sigma-Aldrich G8541-100ML, 1%; MEM non-essential amino acid solution, Sigma-Aldrich, M7145-100ML, 1%). The cells were split every 2-3 days and were not allowed to exceed 90% confluence. For imaging, medium without phenol red was prepared.

#### Plasmids

A cell line stably expressing Tom20-HaloTag was generated based on the pSEMS-Halo backbone (Addgene number 111136) (Appelhans, Richter et al. 2012). For testing the effect of Tom22 and mutants, the SNAP-Tag-Tom22, SNAP-Tag-Tom22-L44E-L48E and SNAP-Tag-Tom22-L48E-L51E were generated and transiently expressed in the Tom20-HaloTag cell line. Transfection was done with PEI (Polyethylenimine, Linear MW 25,000 by Polyscience). The following day the media was exchanged and the cells were kept at 37 °C and 5% CO_2_ before further processing. All genes were of human origin. The plasmids contained genes for the resistances against ampicillin, neomycin/kanamycin, a cytomegalovirus (CMV) promoter and enhancer for gene expression, as well as a T7 promoter. In addition, a mutated Tom20-HaloTag was generated (Tom20-L72K-I75K), targeting the putative substrate binding site.

#### Fluorescence labeling of specimen

Depending on the microscopy technique (confocal Laser Scanning Microscopy (cLSM) or Total Internal Reflection Fluorescence (TIRF)) varying concentrations of dyes were used. To check localization of Tom22 and Tom20, common mitochondrial membranes marker MitoTracker™green (MTG) was used. To covalently label the HaloTag for single particle labeling the Janelia Fluor 646 Halo Tag Ligand (JF646-HTL, Promega), which has a high signal to noise ratio was used. For labeling SNAP-Tom22 variants, Snap-Cell TMR-Star by New England Biolabs was used. For confocal imaging, 30 nM JF646-HTL and 30 nM TMR-Star were added to the cells for 30 min. Unbound dye was washed out. For single molecule imaging, 0.5-1 nM JF646-HTL and 3 nM TMR-Star were used.

#### LSM imaging

For localization studies a confocal microscope (Leica SP8) equipped with a 63x objective (HC PL APO CS2, NA 1.2 water immersion), a tunable white light laser, two hybrid detectors (GAsP) and three photon multiplier detectors (PMT) were used. The imaging was conducted in an environment control chamber with adjusted CO_2_ and temperature. Image view: 1024 x 1024 px, pixel size: 489,24 nm. The excitation wave length for MitoTracker™ Green was 488 nm and the emission was recorded by a hybrid detector (emission: 499-530 nm). Snap-Cell TMR-Star was excited with a wavelength of 561 nm and emission range was 575-630 nm. JF646-HTL was excited with 646 nm wavelength, emission range was 656-695 nm. The detection of fluorescence was performed sequentially to minimize bleed through of signal.

#### TIRF microscopy

The main factor for single particle tracking is a good signal to noise ratio. To minimize background signal, single fluorophores were excited by a highly inclined laminar optical sheet (HiLo)^90^, which operates just below the critical angle for TIRF microscopy. It provides a penetration depth sufficient to excite fluorophores within peripheral mitochondria. For the Tracking and Localization Microscopy (TALM)^91^, an inverse TIRF microscope (Olympus IX83, IX3-TIR-C4-MITICO) equipped with an UApo N 100x/1.49 NA (oil) objective was used. For excitation, 488 nm (200 mW), 561 nm (300 mW) and 646 nm (140 mW) laser diodes were used. The microscope was equipped with a Multisplit (Cairn) with the following filters: 520/35 Brightline HC, 438/24 Brightline HC, 600/37 Brightline HC and a dichroid H 643 LPXR superflat with a TIRF Quad Filter set for all laser lines, and a CMOS camera (Hamamatsu C14440-20UP). Pixel size was 130 nm edge length (2x binning). All measurements were performed at 22 °C. Cells displaying single Tom20 molecules in the JF646-HTL channel (646 nm laser excitation) were recorded for 85 s (5000 images, with 17 ms/frame). For each cell, the angle of the HiLo incident beam was adjusted to ensure a homogeneous signal distribution along the mitochondrial network. The laser output intensity at 646 nm was set between 20% and 60% to ensure homogeneous illumination of single particle signals.

#### Post processing of single molecule data

For data post processing, the Slimfastfaster v1.2A macro, written for MATLAB (v2024b), was used^92^. The pixel size was 130 nm, the photon conversion factor 0.24 and the frame rate 17 ms.

#### Single particle tracking and localization

Single particles of Tom20 have been monitored until bleaching. In the first round of evaluation, particles are localized using a 2D Gaussian fit taking into account the point spread function (PSF). The uncertainty of static localization *ω* is a function of the signal-to-noise ratio, as 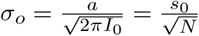, where *N* is the number of photons collected from the PSF (dimension *s*_0_), *a* is the pixel size, *I*_0_ is the peak intensity, when we assume that *ω*_*x*_ = *ω*_*y*_ = *ω*_0_^93^. Finally, trajectories were calculated as described^92, 94^. Trajectories are constructed by linking particles detected within a specified radius of a particle’s previous-frame position. Next, the mean square displacement (MSD) is calculated from single trajectories. The MSD is a statistic measure of the average displacement of a particle in a given time interval (*ε*). The general course of the MSD (t) provides information about the type of diffusion: active transport, anomalous, Brownian, or confined/corralled diffusion^95, 96^. The observed MSD plateau over time suggests anomalous diffusion. Nonetheless, these values represent population averages; individual trajectories may display heterogeneous behavior, such as transitioning from free to confined diffusion or the reverse. The formula for determining the diffusion coefficient *D* for anomalous diffusion from MSD is as follows: MSD(*τ*)= ⟨α*r*(*τ*)^2^ ⟩= 2*dDτ* ^*α*^, where *α<* 1. Here Δ*r*(*τ*) is the displacement, *d* is the spatial dimension, and *γ* is the anomalous sub-diffusion parameter. In case of diffusion in a membrane, *d*=2.

### Methods for yeast experiments

#### Cloning

A yeast plasmid (see Key Resources Table – Recombinant DNA) encoding Tom22 was used as template for quick-change polymerase chain reaction (PCR) to introduce mutations encoded within the primers (see Key Resources Table – Oligonucleotides) into the cytosolic domain of Tom22. Tom22 truncation/fusion constructs and constructs containing TEV protease cleavage sites were generated using KLD cloning (NEB M0554S) and Gibson assembly (NEB M5510A). Tom22 deletion and fusion constructs were combined into a single plasmid by HindIII restriction digestion and ligation. The sequence for the TEV cleavage site (TCS) flanked by Gly-Ala-Gly spacers to enhance cleavage efficiency encoding N’-GAG ENLYFQS GAG-C’ was inserted after the codon for amino acid 53 or 74 of Tom22. In addition, a cytosolic enhanced TEV protease (uTEV3)^97^ under control of the GAL1 promoter was encoded on the same plasmid. Yeast strains (see Key Resources Table – Organisms) were generated through plasmid shuffling. Briefly, cells were transformed with plasmids encoding the respective Tom22 variants, followed by counterselection of the wild-type Tom22 containing plasmid, using 5-fluoroorotic acid (Melford F10501).

#### Yeast serial dilution growth test

Strains were cultured overnight to stationary phase in YPD (10 g/L yeast extract, 20 g/L peptone and 2% glucose) or YPG media (10 g/L yeast extract, 20 g/L peptone and 3% glycerol), diluted and grown to mid log phase. Cells were washed with deionized water, resuspended to an OD_600_ of 1 per mL equivalent cells, serially diluted 10-fold, plated on indicated agar (3% (w/v)) media (YPGal, 10 g/L yeast extract, 20 g/L peptone and 2% galactose; YPG, 10 g/L yeast extract, 20 g/L peptone and 3% glycerol; YPG + Gal, 10 g/L yeast extract, 20 g/L peptone and 3% glycerol, 0.25% galactose), and incubated at indicated temperatures.

#### Whole cell protein extracts

The desired strains were grown overnight at 30°C in YPG, then diluted into the indicated media and grown to mid-log phase. For precursor accumulation assays,^72^ strains were grown at 19°C in YPG to OD_600_ = 1 and shifted to 37°C for 4 h. Cells corresponding to 1 mL of culture at OD_600_ = 1.5 were harvested by centrifugation and washed in 100 µL of H_2_O. Cells were pelleted again, resuspended in 80 µL of H_2_O, and 80 µL of 0.2 M NaOH was added. Following centrifugation, the supernatant was removed and the cells were resuspended in 100 µL of 0.2 M NaOH. The cell suspension was incubated for 5 min at room temperature. Cells were then pelleted again and resuspended in 90 µL 2× Laemmli sample buffer (2α SB: 4% SDS, 20% glycerol, 120 mM Tris-HCl pH 6.8, 0.04% bromophenol blue, 2% β-mercaptoethanol, 2 mM PMSF). Samples were boiled for 5 min at 95°C and separated by Tris-Tricine SDS-PAGE.

#### Isolation of Mitochondria

Yeast were harvested at mid-log phase after growth in YPGal or YPG at 19–23 °C. Mitochondria were isolated as described previously^98^. Briefly, spheroplasts were generated by successive treatments of yeast cells with DTT and zymolyase. Spheroplasts were manually homogenized and mitochondria were isolated by differential centrifugation, quantified by Bradford assay, snap frozen and stored at −80 °C in SEM (250 mM sucrose, 1 mM EDTA, 10 mM MOPS pH 7.2) buffer.

#### Polyacrylamide gel electrophoresis and immunodecoration

Isolated mitochondria were analyzed by (i) Tris-Tricine SDS-PAGE or (ii) blue native-PAGE^99^. (i) Mitochondria were resuspended in 2x Laemmli sample buffer, boiled 5 min at 95 °C and separated by Tris-Tricine SDS-PAGE. (ii) Mitochondria were solubilized in digitonin buffer (20 mM Tris pH 7.4, 0.1 mM EDTA, 50 mM NaCl, 10% glycerol, 1% digitonin) at a 1/10 protein/digitonin ratio for 15 min on ice. Samples were pelleted to remove insoluble material, supplemented with 1x blue native loading dye (10 mM Bis-Tris pH 7.0, 50 mM ε-amino n-caproic acid, 0.5% Coomassie blue G) and loaded onto 6–16.5% blue native-PAGE gels, which were run for 2 h at 20 mA. Gels were incubated 5 min in 1% SDS prior to transfer to PVDF membranes (Immobilon-P, Millipore) by semidry blotting. The membranes were sliced horizontally prior to blocking in TBST (20 mM Tris-HCl pH 7.5, 125 mM NaCl, 1% Tween-20) containing 5% skim milk. Next, membranes were decorated with primary antibody diluted in TBST + 5% milk (see Key Resources Table – Antibodies). Following several washing steps with TBST, membranes were incubated with HRP-conjugated anti-rabbit (1:5000, Jackson ImmunoResearch) secondary antibody in TBST + 5% milk. All membranes were washed several times again in TBST and developed with ECL solution. A LAS-4000 imager (Fujifilm) was used to detect chemiluminescence.

### Molecular dynamics simulations

#### Coarse-grained MD simulations

Coarse-grained (CG) molecular dynamics simulations were employed to study the aggregation of the transmembrane helix of Tom20 on the membrane-exposed surface of the Tom core complex. Starting from the cryo-EM structure of *Sc*TOM_core_ (PDB: 6JNF), we positioned eight copies of the Tom20 TM helix (AF2 model, residues 7–28) evenly distributed at approximately 20 Å around TOM_core_ (see Fig. S1J). The initial CG protein models were generated from the corresponding all-atom structures using Martinize2^100^; the soluble region of Tom22 monomers missing in the cryo-EM structure was not added. The TOM core complex was modeled as a single entity with an elastic network (force constant of 500 kJ·mol^−1^·nm^−2^, cutoff of 0.9 nm) to preserve its tertiary and quaternary structure throughout the simulations. A similar elastic network was applied to maintain the secondary structure of each individual Tom20 helix. Proteins were embedded in a POPC lipid bilayer using the INSANE script^101^ and solvated with Martini water with 0.15 M NaCl. The final simulation box (24 × 24 α 16 nm^3^) contained 1,687 POPC molecules and 42,558 water beads. All CG simulations were performed using GROMACS (version 2021.4)^102^. The Martini 3 force field was employed for proteins, lipids, and water^103^. Electrostatic interactions were treated using a reaction-field approach with a dielectric constant of 15 and a Coulomb cutoff of 1.1 nm. Van der Waals interactions were handled with a potential-shift modification and a cutoff of 1.1 nm. The temperature was maintained at 300 K using the velocity-rescale thermostat^104^ with separate coupling groups for protein, membrane, and solute, each with a coupling time constant of 1.0 ps. The pressure was controlled using a semi-isotropic Berendsen barostat with a coupling time of 5.0 ps, a compressibility of 3×10^−4^ bar^−1^ in the xy plane and the z direction, and a reference pressure of 1 bar.

Each system underwent an initial energy minimization of 5,000 steps using the steepest descent algorithm to remove steric clashes and relax the starting configuration. A first equilibration phase was run for 10 ns with an integration time step of 2 fs, keeping all backbone beads restrained to their initial positions with a harmonic potential (force constant: 1,000 kJ·mol^−1^·nm^−2^). Using the same conditions, a second equilibration step of 500 ns was performed with a time step of 10 fs. Positional restraints on the backbone beads of each Tom20 transmembrane helix were then removed, and ten independent trajectories of 10 µs were generated with a time step of 20 fs.

#### All-atom MD simulations

*TOM complexes*. Starting from the coarse-grained (CG) data, we constructed two all-atom molecular models of the full TOM complex interacting with either Tom20 or Tom70. As a starting point, we selected a conformation from the CG trajectories in which a Tom20 transmembrane helix was stably associated with Tom (cluster 3 in Fig. 6B). This CG structure was then backmapped to atomistic resolution using CG2AT2^105^. To reduce the computational cost while preserving the overall curvature of the membrane along the longitudinal axis of the complex, the initial bilayer was trimmed to a patch of 24 × 16 nm”. The Tom70 assay was generated by manually mutating the residues of the Tom20 transmembrane helix to match the corresponding Tom70 sequence. Missing cytosolic regions of Tom22, Tom20, and Tom70 in both systems were reconstructed in three steps. (1) AlphaFold2^57^ was used to predict the structured segments of the Tom22:Tom20 (Figure 2E,F) and Tom22_1-97_:Tom70 (Figure 3G and S12) complexes. For the latter, AF2 produced two possible orientations of Tom22 with respect to Tom70, both consistent with the NMR data. We ran 0.5 µs-long MD simulations for both models; for this simulation of the whole TOM complex, the C-ter down structure (see Fig. S12) was used. (2) To build the full TOM model, the Tom22:Tom20 and Tom22:Tom70 complexes were positioned above the Tom core in orientations that enabled the generation of the disordered loops linking these complexes to their membrane anchors using MODELLER^106^. (3) The N-terminal disordered regions of Tom22 in complex with Tom20, as well as the complete soluble region of the second Tom22 monomer, were built as random-coil chains with the program ENSEMBLE^73^. This procedure yielded two complete all-atom reconstructions of the TOM complex interacting either with Tom20 or Tom70 embedded in a fully hydrated POPC bilayer. The complete molecular assays (~510,000 atoms, 24 ×24 ×16 nm^3^) resulted in 954 POPC molecules and ~121,000 water molecules. All molecular-dynamics simulations were performed using NAMD 3.0^107^ at 320 K and 1 atm. Water molecules were described by the TIP3P^108^ model and CHARMM36^109^ was employed to model proteins, ions and lipids. The pressure and temperature were maintained using a Langevin barostat^110^ and Langevin dynamics, respectively. Hydrogen mass repartitioning^111^ and the r-RESPA multiple time-step integration algorithm^112^ were employed to integrate the equations of motion with time steps of 4 fs for short-range and 8 fs for long-range interactions. Covalent bonds involving hydrogen atoms were constrained using the SHAKE/RATTLE^113^ and SETTLE^114^ algorithms. Long-range electrostatic interactions were treated using the particle-mesh Ewald method^115^. A 12-Å cutoff was applied to truncate both van der Waals and short-range Coulombic interactions. After 1,000 minimization steps, four repeats of the following protocol were conducted for both assays. First, systems were equilibrated for 80 ns in the presence of soft harmonic restraints maintaining every backbone atom at its initial position. Next, all harmonic restraints applied to extracellular regions of Tom22, Tom20 and Tom70 were removed, and the dynamics of these segments were sampled for 250 ns.

*HsTom22cyt (PDB:7VDD) and RnTom20 cyt (PDB:3AWR)*. Two additional all-atom assays were constructed to investigate the structural stability of the previously proposed Tom22_cyt_ conformation^23^, and the conformation of Tom20_cyt_ (Fig. S1C-F): the proposed Tom22_cyt_ was extracted in the cryo-EM structure of the *Homo sapiens* TOM_core_ complex (PDB:7VDD, residues 1-87). The *Rn*Tom20 structure had been obtained by NMR structure (PDB:3AWR). Each of these proteins were embedded into a cubic water box at 0.15 M NaCl of 9 x 9 x 9 nm^3^ (20,700 water molecules; for Tom22) and 7.2 x 7.2 x 7.2 nm^3^ (10,700 water molecules; for Tom20). Using the same MD parameters as reported above, and following 500 minimization steps, both systems were first thermalized at 300 K for 1.5 ns, with harmonic restraints on every protein heavy atom. The restraints were subsequently removed stepwise, and the dynamics of each of the two proteins were sampled for 0.8 µs.

**Table S1.**
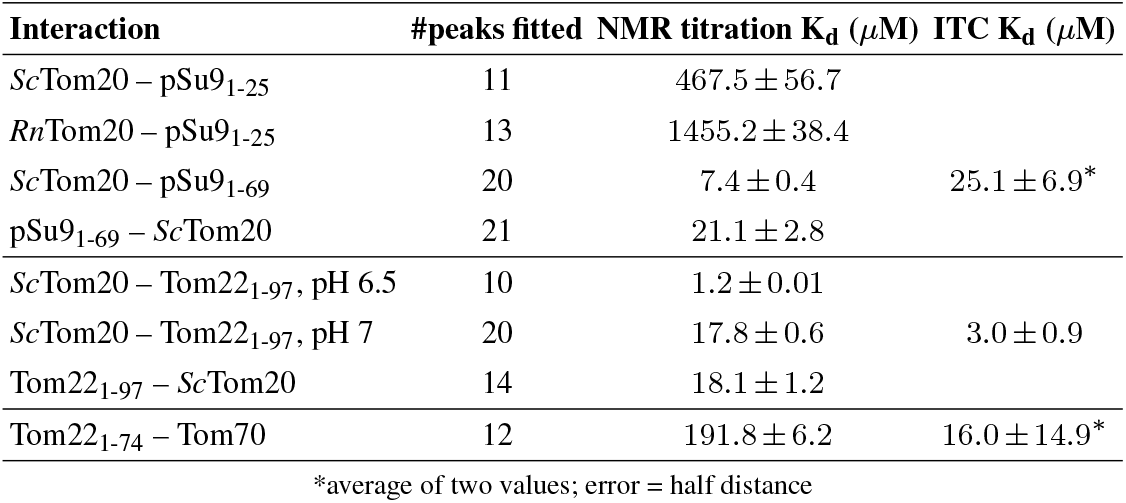
Dissociation constants (K_d_) obtained from NMR titration experiments and from ITC. For the NMR analysis (with TITAN) a two-state ligand binding model was used for the fitting of 2D ^1^H-^15^N NMR spectra. A global analysis, using a number of peaks (as reported here) was used. ITC K_d_ values shown are weighted averages of three (*Sc*Tom20–Tom22) independent ITC measurements or simple averages (*Sc*Tom20–pSu9 and Tom22–Tom70) of two independent measurements. See also the ITC data in Fig. S10.

| **Interaction** | **#peaks fitted** | **NMR titration K<sub>d</sub> (*μ*M)** | **ITC K<sub>d</sub> (*μ*M)** |
|---|---|---|---|
| *Sc*Tom20 – pSu9<sub>1-25</sub> | 11 | 467.5 ± 56.7 |  |
| *Rn*Tom20 – pSu9<sub>1-25</sub> | 13 | 1455.2 ± 38.4 |  |
| *Sc*Tom20 – pSu9<sub>1-69</sub> | 20 | 7.4 ± 0.4 | 25.1 ± 6.9* |
| pSu9<sub>1-69</sub> – *Sc*Tom20 | 21 | 21.1 ± 2.8 |  |
| *Sc*Tom20 – Tom22<sub>1-97</sub>, pH 6.5 | 10 | 1.2 ± 0.01 |  |
| *Sc*Tom20 – Tom22<sub>1-97</sub>, pH 7 | 20 | 17.8 ± 0.6 | 3.0 ± 0.9 |
| Tom22<sub>1-97</sub> – *Sc*Tom20 | 14 | 18.1 ± 1.2 |  |
| Tom22<sub>1-74</sub> – Tom70 | 12 | 191.8 ± 6.2 | 16.0 ± 14.9* |

\*average of two values; error = half distance

