## Supplementary Figures for "A dynamic displacement mechanism drives protein import into mitochondria"

**Supplementary Movie 1.** Molecular dynamics simulations of the *Sc*TOM$_{core}$ complex with Tom20, corresponding to Fig. 6C. The movie corresponds to four trajectories, each 250 ns long, i.e. the total duration of the simulation shown here is 1 microsecond. The same set of trajectories is shown on the left and right, seen from the top and side, respectively.

**Supplementary Movie 2.** Molecular dynamics simulations of the *Sc*TOM$_{core}$ complex with Tom70, corresponding to Fig. 6D. The movie corresponds to four trajectories, each 250 ns long, i.e. the total duration of the simulation shown here is 1 microsecond. The same set of trajectories is shown on the left and right, seen from the top and side, respectively.

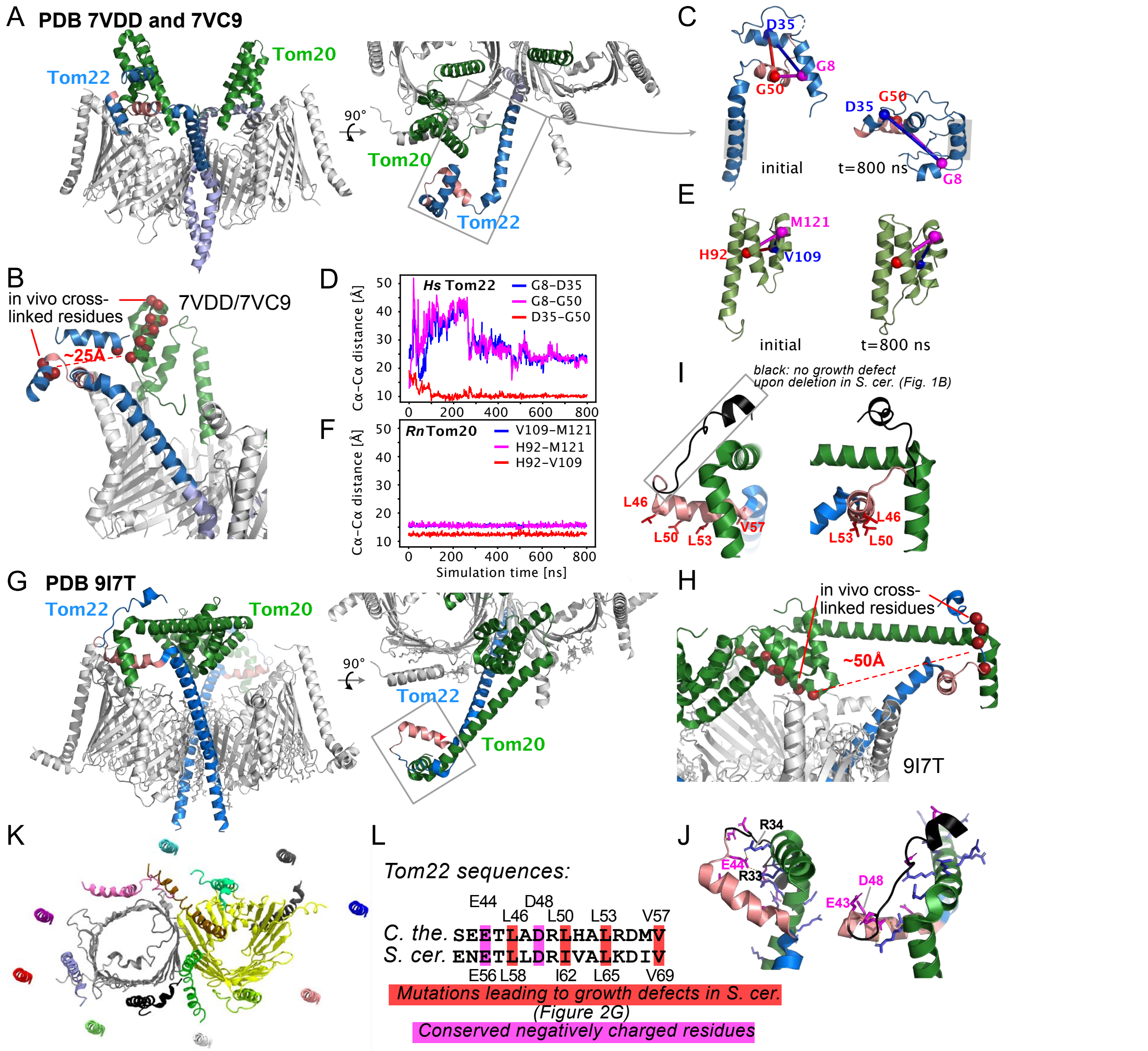


**Figure S1. Analysis of previously proposed TOM conformations, and molecular dynamics investigations**

**A**, Cryo-EM model obtained with the soluble cross-linking agent glutaraldehyde[23] (PDB entries 7VDD and 7VC9) seen from the side (left) and top (right). In addition to Tom22's extended TM helix, the model comprises a compact arrangement of three helices in the cytosolic part. Tom22 is shown in blue, and the Tom22$_{PPDE}$ element in pink; Tom20 is shown in green.

**B** Tom20 and Tom22 residues resulting in cross-linking products [29] are mapped on the structure reported by Su *et al.* (red spheres). The dashed arrow indicates the distance between the two cross-linked regions. With a distance of 25 Å, it is not likely that these residues could cross-link in vivo.

**C**, Investigation of the stability of the proposed compact three-helix fold of *Hs*Tom22$_{cyt}$ from panel **A** through all-atom MD simulations. The structures shown on the left and right are the conformations at the beginning of the simulation and after 800 ns, respectively, aligned to the helical part indicated by a grey box.

**D**, Evolution of the distances of the three helices of this Tom22 model (panel C) over the simulation time, highlighting the melting of the tertiary structure. Shown are the distances between the Cα atoms of the three indicated residues. This simulation strongly suggests that the proposed model[23] is not a stable conformation.

**E,F**, Equivalent simulations of *Rn*Tom20, using the NMR structure[31] as starting point, showing that the fold is highly stable.

**G**, Cryo-EM structure of the TOM$_{core}$ complex[26] of *Chaetomium thermophilum* TOM$_{core}$ (PDB 9I7T). In this structure, Tom22 interacts via its Tom22$_{PPDE}$ element with Tom20 at a kink formed after the TM helix of the latter. Note the dramatic differences in the Tom22 architecture in the two models (a) and (b).

**H** Similar to panel B, but mapped on the structure reported by Agip *et al.*. With distance of 50 Å, it is not likely that these residues could cross-link in vivo.

**I**, Zoom onto the Tom20-Tom22 interface in the structure shown in (B); PDB 9I7T), highlighting the conserved hydrophobic residues (red). Mutations of these residues showed yeast growth defects (Figs. 1B,G and 2E). None of these hydrophobic residues of Tom22 is within 6 Å from any residue of Tom20 in this structure, despite the functional importance shown by the growth defect data. Deletion of the part shown in black, proposed in the study by Agip *et al.* to be important, did not result in growth defects (Fig. 1B), suggesting that the interaction surface proposed in this model is not essential.

**J**, The report by Agip *et al.* suggested that charge interactions are important for the Tom20-Tom22 interaction. This panel shows the location of charged residues in the Tom20-Tom22 interface. The two intermolecular distances between oppositely charged residues shorter than 3 Å are indicated (E44 of Tom22). Of note, in *N. crassa* the simultaneous replacement of up to 15 negatively charged (D, E) residues by neutral ones (N, Q), including the position equivalent to E44, did not result in growth defects[64]. A doubly positive mutation in *S. cerevisiae* Tom22 (N55R, E56K, where E 56 corresponds to E44 of *C. thermophilum* Tom22), led to growth defects[33]; we hypothesize that these two positive charges may interact with negative ones of Tom22$_{cyt}$, hampering its interactions with binding partners. The structure of the human TOM/VDAC/PINK1 array points to a similar arrangement as shown here, but the part corresponding to the pink helix was not resolved[27] and the binding was proposed to be due to hydrophobic contacts rather than electrostatic ones.

**K**, Positions of the Tom20 TM helices around the TOM$_{core}$ complex used as starting configuration of the coarse-grained simulations of Fig. 6B. Ten simulations from this configuration with different initial velocities were performed. The positions of the 14 Tom20$_{TM}$ helices that remained stably TOM$_{core}$-bound are reported in Fig. 6B.

**L**, Alignment of Tom22 sequences of *Chaetomium thermophilum* and *Saccharomyces cerevisiae*, highlighting the conserved hydrophobic residues (red) and conserved negative charges (magenta), indicated in (G) and (H), respectively.

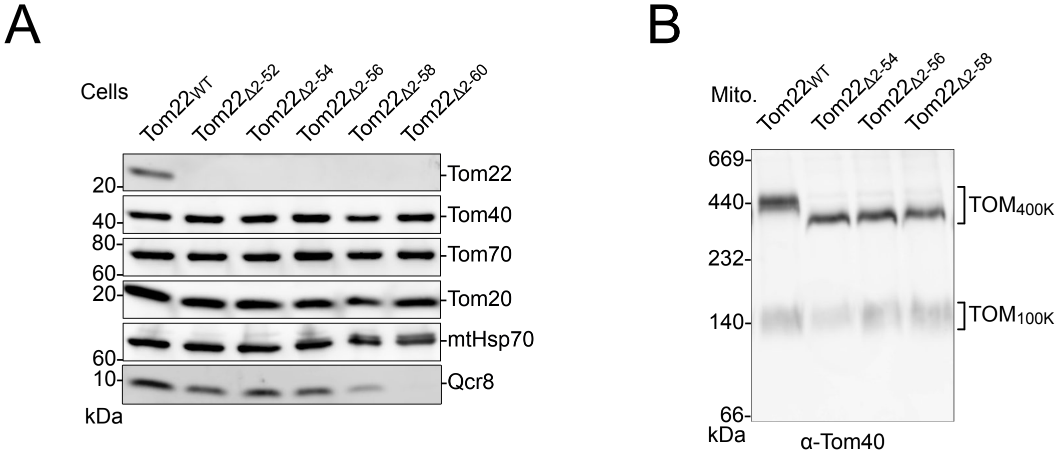


**Figure S2. Tom protein and TOM complex levels in strains with N-terminal truncation of Tom22, related to Figure 1B.**

**A**, Whole cell extract/immunodecoration reveals comparable levels of Tom40, Tom70 and Tom20 in yeast strains with N terminal truncations of Tom22. The antibody directed against the N-terminus of Tom22 confirms the N terminal truncations. The available Tom22 antibodies recognize an N terminal part which is missing in the truncation mutants. The decreasing levels of Qcr8 (subunit 8 of the mitochondrial respiratory chain complex III/cytochrome $bc_1$ complex) indicate progressive mitochondrial defects caused by longer N-terminal truncations of Tom22.

**B**, Isolated mitochondria were lysed with digitonin and subjected to blue-native PAGE/immunodecoration. The Tom22 variants assemble efficiently into mature $TOM_{400K}$ complexes of lowered molecular mass reflecting the N terminal truncations with a $TOM_{400K}$ (TOM with Tom22)/$TOM_{100K}$ (TOM without Tom22) ratio comparable to wild type (WT).

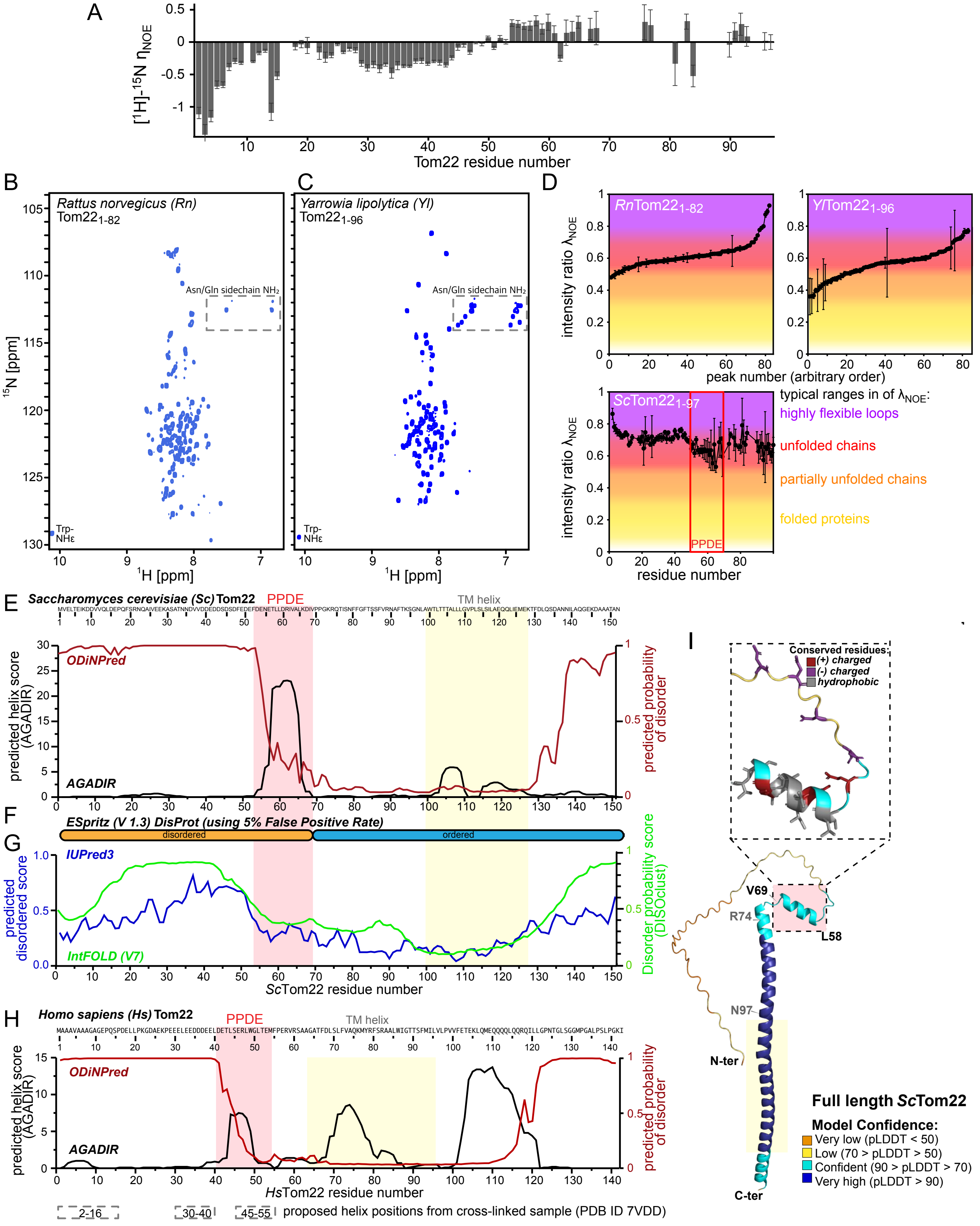
A
[1H]-15N ηNOE
Tom22 residue number
B
Rattus norvegicus (Rn) Tom22 1-82
Asn/Gln sidechain NH₂
Trp-NHε
15N [ppm]
1H [ppm]
C
Yarrowia lipolytica (Yl) Tom22 1-96
Asn/Gln sidechain NH₂
Trp-NHε
1H [ppm]
D
RnTom22 1-82
YlTom22 1-96
intensity ratio λNOE
peak number (arbitrary order)
ScTom22 1-97
PPDE
residue number
typical ranges in of λNOE:
highly flexible loops
unfolded chains
partially unfolded chains
folded proteins
E
Saccharomyces cerevisiae (Sc) Tom22
PPDE
TM helix
ODiNPred
AGADIR
predicted helix score (AGADIR)
predicted probability of disorder
F
ESpritz (V 1.3) DisProt (using 5% False Positive Rate)
disordered
ordered
G
IUPred3
IntFOLD (V7)
predicted disordered score
Disorder probability score (DISOclust)
ScTom22 residue number
H
Homo sapiens (Hs) Tom22
PPDE
TM helix
ODiNPred
AGADIR
predicted helix score (AGADIR)
predicted probability of disorder
HsTom22 residue number
2-16
30-40
45-55
proposed helix positions from cross-linked sample (PDB ID 7VDD)
I
Conserved residues:
(+) charged
(-) charged
hydrophobic
V69
R74
L58
N97
N-ter
C-ter
Full length ScTom22
Model Confidence:
Very low (pLDDT < 50)
Low (70 > pLDDT > 50)
Confident (90 > pLDDT > 70)
Very high (pLDDT > 90)

**Figure S3. (Preceding page) Intrinsic disorder and transient $\alpha$-helix formation in Tom22 from different organisms**

**A**, Residue-wise [$^1$H]-$^{15}$N heteronuclear-NOE relaxation values in Tom22$_{1\text{-}97}$. Negative values, typically found in disordered proteins[116] are observed for residues 1-53, while positive values for E54 to D68 (average value 0.3) reveal residual structure in the Tom22$_{PPDE}$.

**B,C**, $^1$H-$^{15}$N correlation spectra of *Rn*Tom22$_{1\text{-}82}$ (B) and *Yl*Tom22$_{1\text{-}96}$ (C). These spectra show that the Tom22$_{cyt}$ segments of these two organisms are disordered.

**D**, Semi-quantitative assessment of the local structural compactness of these two Tom22 constructs using the HET-SOFAST NMR approach[117]. The parameter $\lambda_{NOE}$ reports on the local compactness, at the location of each amide site, averaged over all conformations present. As the NMR spectra of RnTom22$_{1\text{-}82}$ and *Yl*Tom22$_{1\text{-}96}$ have not been assigned sequence-specifically, these values, obtained for all amide signals, are reported in an arbitrary order (increasing $\lambda_{NOE}$). The lower left panel in **D**, shows the data for *Sc*Tom22$_{1\text{-}97}$, as a function of the sequence position. The Tom22$_{PPDE}$ element has lower values, in line with the fact that it forms a transient helix. The typical ranges of $\lambda_{NOE}$ for different types of structural order are indicated in colors, as described[117]. This data shows more quantitatively that the three Tom22$_{cyt}$ constructs are largely disordered.

**E**, Sequence-based predictions of local helix propensity using AGADIR[55] (black) and of disorder using ODINPred[56] for full-length *Sc*Tom22.

**F,G**, Disorder predictions of full-length *Sc*Tom22 using ESpritz[118], IUPred3[119] and IntFold[120].

**H**, As **E**, for *Hs*Tom22. The dashed boxes below indicate the location of $\alpha$-helices proposed in the model of chemically cross-linked TOM complex[23].

**I**, AlphaFold2 model of *Sc*Tom22. The Tom22$_{PPDE}$ element is highlighted in the insert.

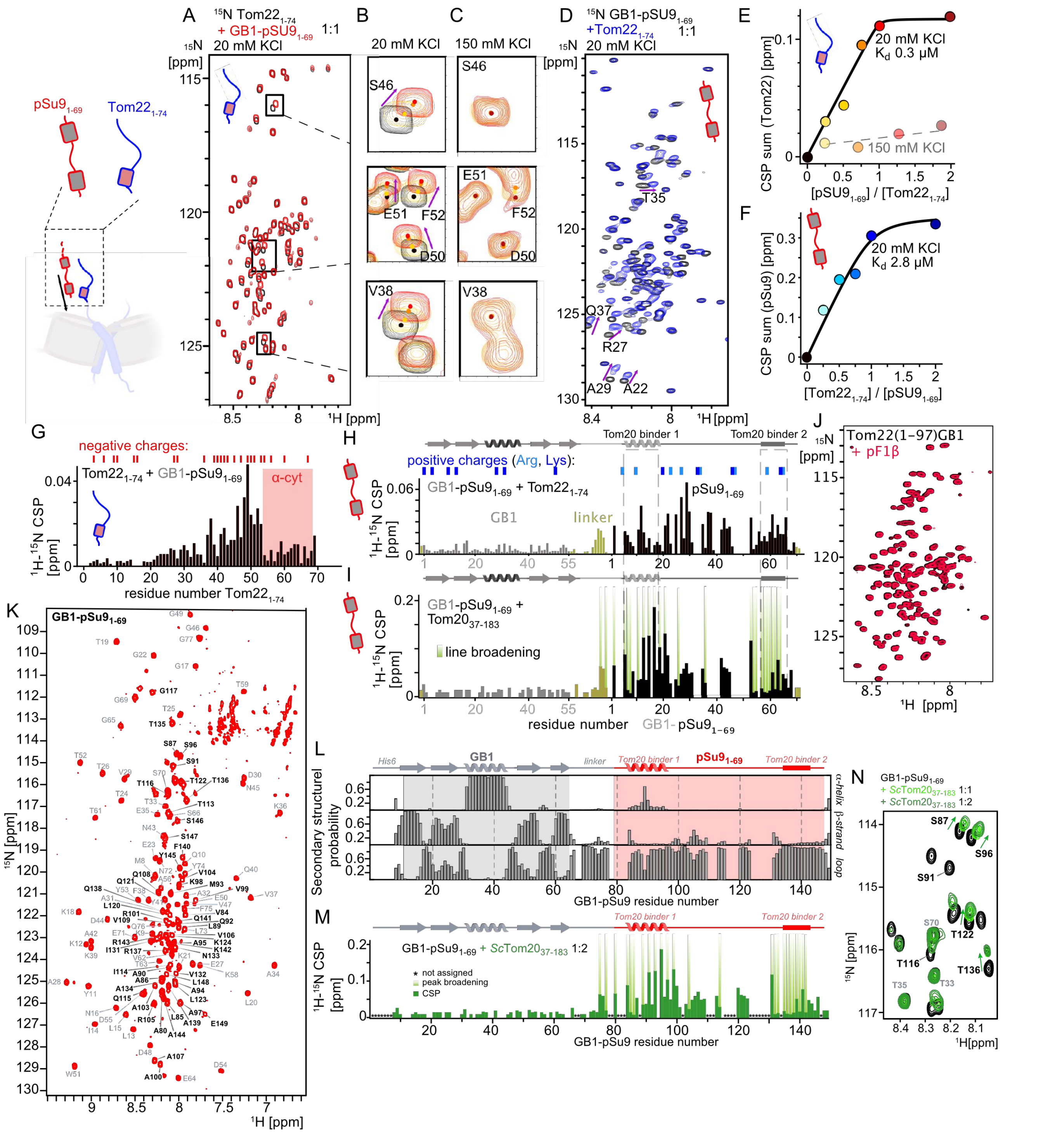

A
B
C
D
E
F
G
H
I
J
K
L
M
N
pSu9₁₋₆₉
Tom22₁₋₇₄
¹⁵N Tom22₁₋₇₄ + GB1-pSU9₁₋₆₉ 1:1
20 mM KCl
150 mM KCl
¹⁵N GB1-pSU9₁₋₆₉ +Tom22₁₋₇₄ 1:1
20 mM KCl Kd 0.3 µM
150 mM KCl
20 mM KCl Kd 2.8 µM
CSP sum (Tom22) [ppm]
CSP sum (pSu9) [ppm]
[pSU9₁₋₆₉] / [Tom22₁₋₇₄]
[Tom22₁₋₇₄] / [pSU9₁₋₆₉]
negative charges:
Tom22₁₋₇₄ + GB1-pSu9₁₋₆₉
α-cyt
¹H-¹⁵N CSP [ppm]
residue number Tom22₁₋₇₄
positive charges (Arg, Lys):
Tom20 binder 1
Tom20 binder 2
GB1-pSu9₁₋₆₉ + Tom22₁₋₇₄
GB1
linker
pSu9₁₋₆₉
GB1-pSu9₁₋₆₉ + Tom20₃₇₋₁₈₃
line broadening
residue number GB1- pSu9₁₋₆₉
Tom22(1-97)GB1 + pF1β
GB1-pSu9₁₋₆₉
¹⁵N [ppm]
¹H [ppm]
His6
Secondary structure probability
α-helix
β-strand
loop
GB1-pSu9 residue number
GB1-pSu9₁₋₆₉ + ScTom20₃₇₋₁₈₃ 1:2
not assigned
peak broadening
CSP
GB1-pSu9₁₋₆₉ + ScTom20₃₇₋₁₈₃ 1:1 + ScTom20₃₇₋₁₈₃ 1:2

**Figure S4. (Preceding page) Unfolded Tom22$_{cyt}$ interacts with a presequence through charge interactions at low salt concentration**

**A**, $^{1}$H-$^{15}$N spectra of *Sc*Tom22$_{1\text{-}74}$ (80 µM) in the absence (black) and presence (red) of the 69-residue presequence of *N. crassa* subunit 9 of the F0-ATPase (GB1-pSu9$_{1\text{-}69}$, 70 µM, unlabeled) in low-salt buffer (20 mM KCl, 20 mM KPi, pH 6.5). The observed chemical-shift perturbations (CSPs) show that these two proteins interact, and allow identifying which residues are involved (see panel G).

**B**, Zoom onto three regions highlighted in (**A**). Shown are spectra with 0 (black), 40 (yellow), 60 (orange) and 80 µM (red) GB1-pSu9$_{1\text{-}69}$.

**C**, Spectra of NMR-visible ($^{15}$N labeled) *Sc*Tom22$_{1\text{-}74}$ (75 µM) with different concentrations of GB1-pSu9$_{1\text{-}69}$ (ratios 1:0.25 yellow; 1:0.75 orange; 1:2 dark red) at 150 mM KCl. No significant CSPs are detected, showing that at this salt concentrations these two proteins do not interact. The full spectra are displayed in Fig. S5D.

**D**, $^{1}$H-$^{15}$N spectrum of GB1-pSu9$_{1\text{-}69}$ (70 µM; black) with *Sc*Tom22$_{1\text{-}74}$ (70 µM; unlabeled). Arrows indicate examples of residues with large chemical-shift perturbations. See panel H for a residue-wise plot of these CSPs.

**E**, Fit of combined CSPs from residues with large CSPs (41,46,49,51,52,53); the solid line corresponds to a $K_d$ of 0.3 µM. The summed CSP for the same residues in the presence of 150 mM salt is indicated with a dashed line, showing the absence of detectable interaction.

**F**, Fit of the combined CSPs from the 12 residues of GB1-pSu9$_{1\text{-}69}$ with largest CSP; the solid line corresponds to a $K_d$ of 2.8 µM.

**G**, Residue-wise plots of the CSP effects in *Sc*Tom22$_{1\text{-}74}$ in the presence of GB1-pSu9$_{1\text{-}69}$ at a 1:1 molar ratio. Negative charges (Asp, Glu) are indicated above (red bars) and on a conformation of Tom22$_{cyt}$. (Red spheres denote C$\alpha$ atoms of Asp and Glu residues.)

**H**, Residue-wise plots of the CSP effects in GB1-pSu9$_{1\text{-}69}$ in the presence of *Sc*Tom22$_{1\text{-}74}$ at a 1:1 molar ratio. Positively charged residues are indicated above (blue bars).

**I**, CSP effect in GB1-pSu9$_{1\text{-}69}$ upon addition of *Sc*Tom20$_{37\text{-}183}$. Residues with significant line broadening are indicated by green gradient-colored bars.

**J**, $^{1}$H-$^{15}$N spectrum of *Sc*Tom22$_{1\text{-}74}$ (80 µM) in the absence (black) and presence (red) of the presequence F1$\beta$ (160 µM) at 100 mM KCl concentration, showing no significant change. Additional spectra with other presequences are shown in Fig. S5.

**K**, $^{1}$H-$^{15}$N spectrum of GB1-pSu9$_{1\text{-}69}$. To obtain the numbering that starts with 1 for residue 1 of the pSu9 sequence, 78 shall be subtracted from the numbers displayed. The resonance assignment is deposited in the BioMagResBank under accession number 51093.

**L**, TALOS-N derived populations of $\alpha$-helical (top), $\beta$-strand (middle) and disordered/loop (bottom) conformations of the GB1-pSu9 construct. A small population of $\alpha$-helical conformation is detected for the first presequence (corresponding to ca. residues 7-20; 85-98 here), while the second presequence does not show $\alpha$-helical conformations.

**M**, Combined $^{1}$H,$^{15}$N chemical-shift perturbation in GB1-pSu9 upon addition of a 2-fold excess of *Sc*Tom20$_{37\text{-}183}$.

**N**, Zoom of the spectra of GB1-pSu9$_{1\text{-}69}$ in the absence (grey) and presence of *Sc*Tom20$_{37\text{-}183}$.

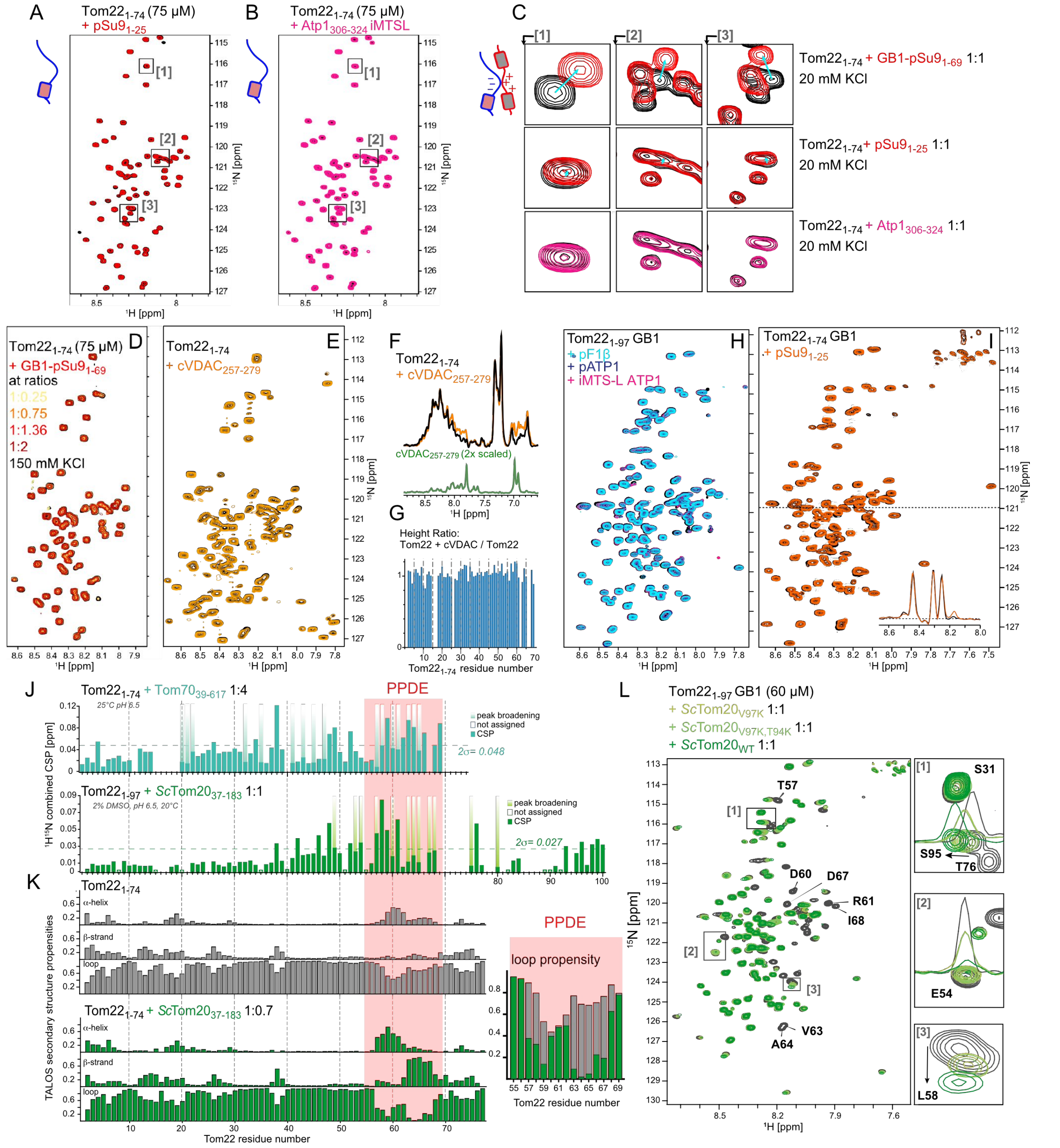

A
Tom22 1-74 (75 µM)
+ pSu9 1-25
B
Tom22 1-74 (75 µM)
+ Atp1 306-324 iMTSL
C
Tom22 1-74 + GB1-pSu9 1-69 1:1
20 mM KCl
Tom22 1-74 + pSu9 1-25 1:1
20 mM KCl
Tom22 1-74 + Atp1 306-324 1:1
20 mM KCl
D
Tom22 1-74 (75 µM)
+ GB1-pSu9 1-69
at ratios
1:0.25
1:0.75
1:1.36
1:2
150 mM KCl
E
Tom22 1-74
+ cVDAC 257-279
F
Tom22 1-74
+ cVDAC 257-279
cVDAC 257-279 (2x scaled)
G
Height Ratio:
Tom22 + cVDAC / Tom22
Tom22 1-74 residue number
H
Tom22 1-97 GB1
+ pF1β
+ pATP1
+ iMTS-L ATP1
I
Tom22 1-74 GB1
+ pSu9 1-25
J
Tom22 1-74 + Tom70 39-617 1:4
25°C pH 6.5
PPDE
peak broadening
not assigned
CSP
2σ= 0.048
Tom22 1-97 + ScTom20 37-183 1:1
2% DMSO, pH 6.5, 20°C
2σ= 0.027
1H15N combined CSP [ppm]
K
Tom22 1-74
α-helix
β-strand
loop
Tom22 1-74 + ScTom20 37-183 1:0.7
TALOS secondary structure propensities
Tom22 residue number
PPDE
loop propensity
Tom22 residue number
L
Tom22 1-97 GB1 (60 µM)
+ ScTom20 V97K 1:1
+ ScTom20 V97K,T94K 1:1
+ ScTom20 WT 1:1
T57
D60
D67
R61
I68
V63
A64
S31
S95
T76
E54
L58
1H [ppm]
15N [ppm]

**Figure S5. (Preceding page) $Tom22_{cyt}$ does not show significant interaction with preprotein peptides at physiologically relevant salt concentration, but interacts with Tom20.**

Spectra of $^{15}$N-labeled $Tom22_{1\text{-}74}$ in its apo state (always in black) and upon adding different presequence peptides:

**A**, $pSu9_{1\text{-}25}$

**B**, $Atp1_{306\text{-}324}$

**C**, Close-up of Tom22 spectra with GB1-$pSu9_{1\text{-}69}$. The shown parts correspond to those indicated with rectangles in panels **A** and **B**.

**D**, Spectra of *Sc*$Tom22_{1\text{-}74}$ with different concentrations of GB1-$pSu9_{1\text{-}69}$.

**E**, Spectra of $^{15}$N-labeled $Tom22_{1\text{-}74}$ without and with a cyclic peptide corresponding to human $VDAC1_{257\text{-}279}$ (unlabeled). The addition of the peptide is shown by the additional $^{1}$H signals (orange in **F**; the green spectrum is a sample of the peptide alone), but the lack of any spectral changes in the $^{1}$H-$^{15}$N spectrum (**E**), including the absence of sizable intensity changes (**G**), shows that there is no interaction.

**H, I**, Equivalent analyses to panel B, using either the presequence of the ATP synthase subunit β (pF1$\beta_{21\text{-}33}$; APLLSTSWKRSM; Uniprot ID P00830; note a replacement of C by S), the presequence of the α-subunit of the ATP synthase ($pATP1_{1\text{-}20}$), the internal matrix targeting signal of ATP1, $Atp1_{306\text{-}324}$, and the presequence of the *N. crassa* subunit 9 of the F0-ATPase ($pSu9_{1\text{-}25}$). In all cases, the protein concentration was 80 μM, and the peptide was in 2-fold molar excess; the buffer was 50 mM KPi, 100 mM KCl, pH 6.5.

**J**, Chemical-shift perturbation of Tom22 upon addition of either Tom20 or Tom70, as indicated. Bars with gradient color show significant line broadening in the presence of the binding partner.

**K**, Secondary-structure analysis of Tom22 apo (grey) and in the presence of Tom20 (green). Shown is the per-residue propensity to form α-helix, β-strand or unfolded (loop) conformations, as indicated. A clear increase in secondary structure (and decrease in unfolded conformations) upon binding Tom20 indicates a conformational-selection process, in which the population of unfolded state is shifted toward a state with secondary structure, as expected from the structural models (Fig. 2C,D).

**L**, $Tom22_{1\text{-}97}$ spectra in the presence of different Tom20 mutants, as indicated.

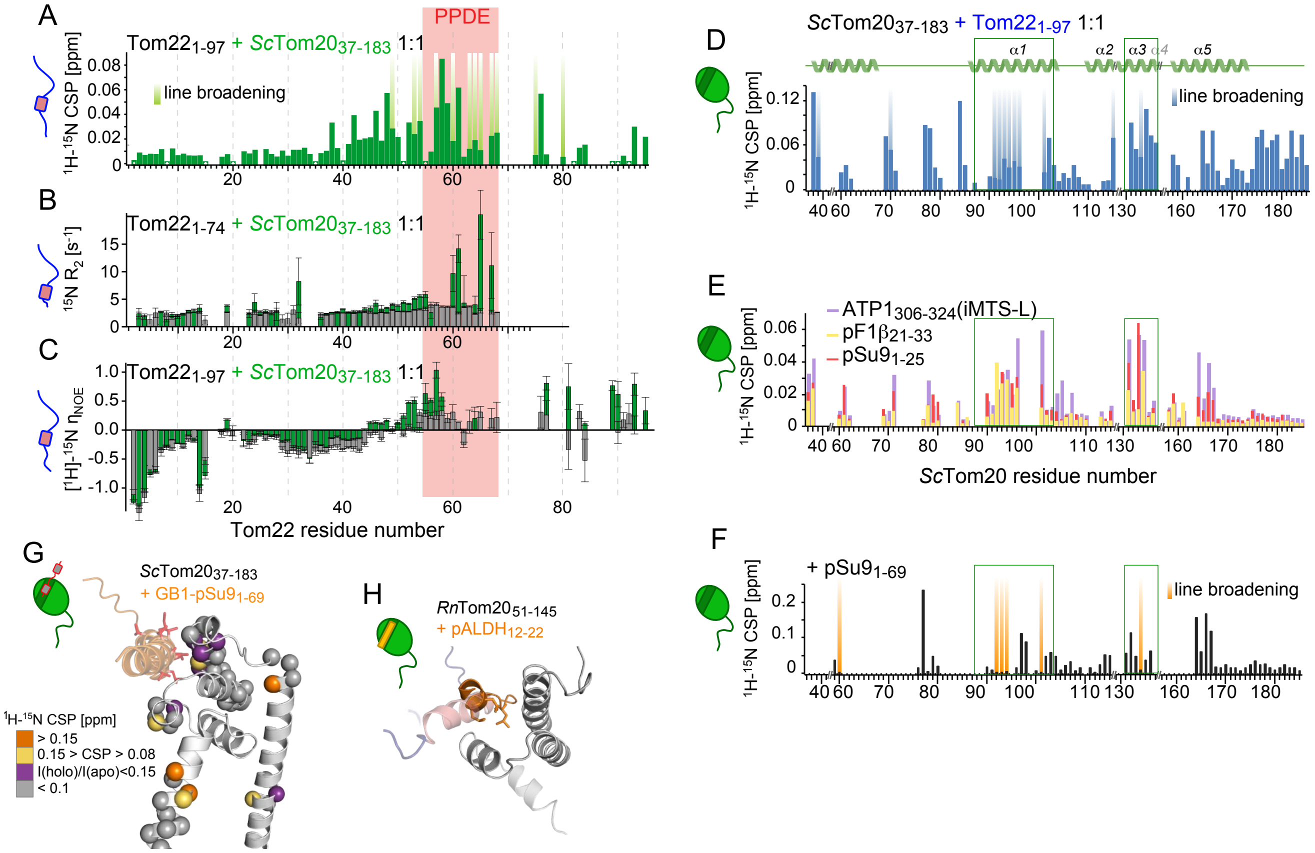


**Figure S6. NMR data of interactions of *Sc*Tom22 with *Sc*Tom20, and of *Sc*Tom20 with *Sc*Tom22, pSu9 and targeting sequences**

**A**, $^{1}$H-$^{15}$N chemical shift perturbations (CSP) in *Sc*Tom22$_{1\text{-}97}$ induced by addition of equimolar *Sc*Tom20$_{37\text{-}183}$.

**B**, $^{15}$N R$_2$ relaxation rate in *Sc*Tom22$_{1\text{-}74}$ in the absence and presence of *Sc*Tom20$_{37\text{-}183}$ (molar ratio 1:1), acquired at 14 T B$_0$ field strength. The elevated $^{15}$N R$_2$ relaxation rates observed for several residues in *Sc*Tom22$_{\text{PPDE}}$ – D60, R61, L65, D67 – are characteristic of binding/release dynamics on the µs-ms timescale, typical of micromolar affinity complexes[121].

**C**, [$^{1}$H]-$^{15}$N hetNOE values in *Sc*Tom22$_{1\text{-}97}$ without (grey) and with (green) *Sc*Tom20, indicating a decrease in local mobility for Tom22 residues within Tom22$_{\text{PPDE}}$.

**D**, $^{1}$H-$^{15}$N CSP in *Sc*Tom20$_{37\text{-}183}$ induced by the addition of *Sc*Tom22$_{1\text{-}97}$ (1:2). Bars with a color gradient indicate peaks with strongly (>10-fold) diminished intensity in the complex. The numbering of helices above is according to Abe *et al.* 2000, and in line with the numbering of *Rn*Tom20; the additional N-terminal α-helix is predicted by AlphaFold.

**E**, As in D, but with different targeting peptides, as indicated. The largest CSP or intensity reduction effects are observed in the same region as with *Sc*Tom22 (D). The molar ratio *Sc*Tom20$_{37\text{-}183}$ : pSu9$_{1\text{-}69}$ is 1:2, while the ratio of *Sc*Tom20$_{37\text{-}183}$ to pSu9$_{1\text{-}25}$, ATP1$_{306\text{-}324}$ or pF1$_{21\text{-}33}$ is 1:5.

**F**, $^{1}$H-$^{15}$N CSPs in *Sc*Tom20$_{37\text{-}183}$ in the presence of the presequence of subunit 9 of F$_0$-ATPase protein (pSu9$_{1\text{-}69}$).

**G**, CSP from panel F plotted onto the AlphaFold model of the Tom20-pSu9 complex.

**H**, Previously reported NMR structure of the *Rn*Tom20-pALDH$_{12\text{-}22}$ interaction. Additionally, the position of Tom22 is indicated (as in Fig. 2E,F).

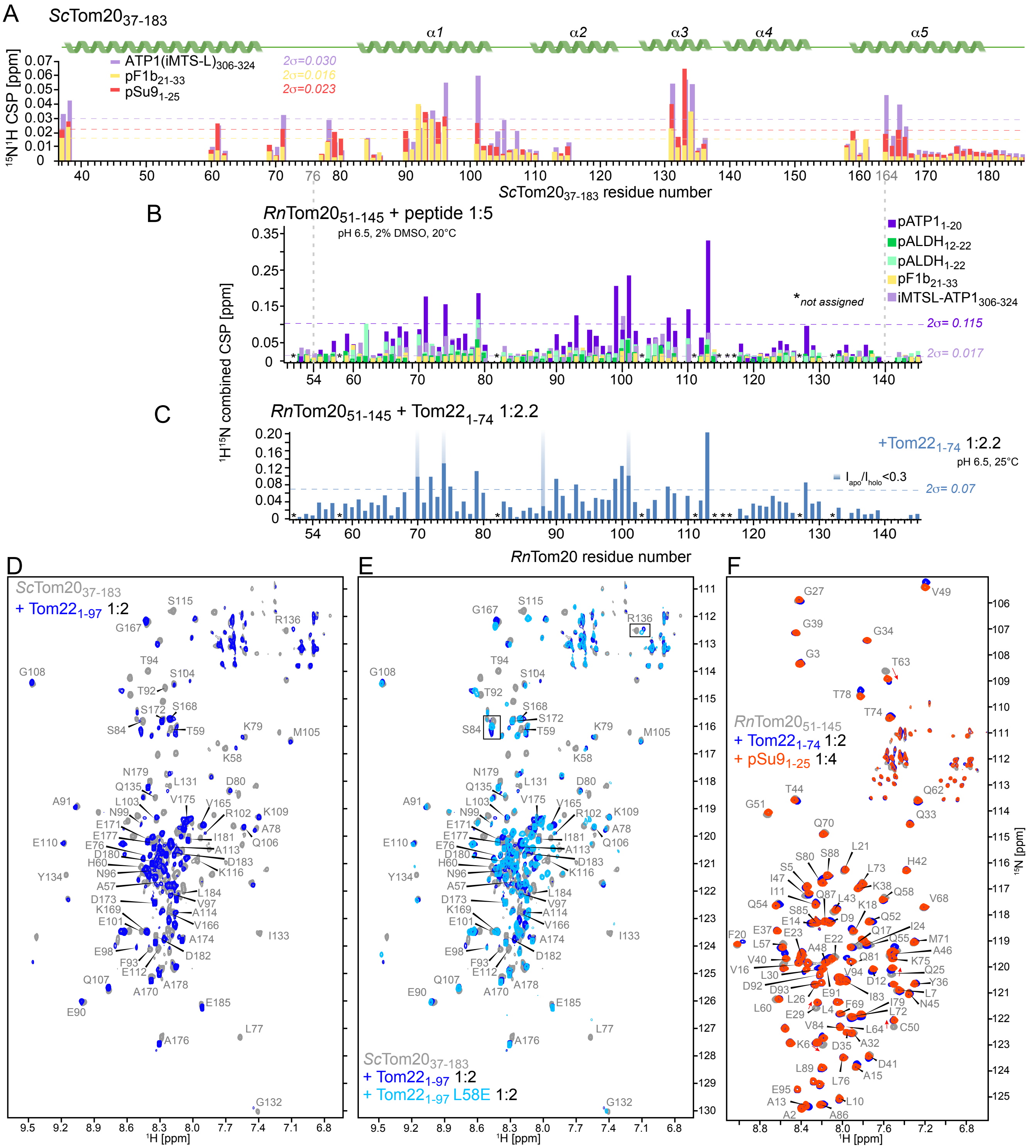
A
ScTom20 37-183
α1
α2
α3
α4
α5
ATP1(iMTS-L)306-324
pF1b21-33
pSu91-25
2σ=0.030
2σ=0.016
2σ=0.023
15N1H CSP [ppm]
ScTom20 37-183 residue number
B
RnTom20 51-145 + peptide 1:5
pH 6.5, 2% DMSO, 20°C
pATP1 1-20
pALDH 12-22
pALDH 1-22
pF1b 21-33
iMTSL-ATP1 306-324
*not assigned
2σ= 0.115
2σ= 0.017
1H15N combined CSP [ppm]
C
RnTom20 51-145 + Tom22 1-74 1:2.2
+Tom22 1-74 1:2.2
pH 6.5, 25°C
Iapo/Iholo<0.3
2σ= 0.07
RnTom20 residue number
D
ScTom20 37-183
+ Tom22 1-97 1:2
E
ScTom20 37-183
+ Tom22 1-97 1:2
+ Tom22 1-97 L58E 1:2
F
RnTom20 51-145
+ Tom22 1-74 1:2
+ pSu9 1-25 1:4
1H [ppm]
15N [ppm]

**Figure S7. (Preceding page) Tom20 proteins from rat (*Rn*) and yeast (*Sc*) bind presequences and Tom22 in the client binding pocket**

**A**, Combined $^1$H-$^{15}$N chemical shift perturbations of yeast Tom20$_{37\text{-}183}$ in presence of presequence peptides pF1$\beta$, pSu9 and internal matrix targeting sequence of ATP1. Approximately 50% of the backbone resonances of *Sc*Tom20$_{37\text{-}183}$ were assigned (see Methods). The numbering of $\alpha$-helices, indicated above, follows the numbering of the *Rn*Tom20 by Abe *et al.*.

**B**, $^1$H-$^{15}$N CSPs of *Rn*Tom20$_{51\text{-}145}$ with five times molar excess of indicated presequence peptides. Vertical dashed lines between panels A and B represent equivalent protein sequence between Tom20 from two organisms, derived from multiple sequence alignment.

**C**, $^1$H-$^{15}$N CSPs of *Rn*Tom20$_{51\text{-}145}$ induced by the presence of a 2.2-fold excess of *Sc*Tom22$_{1\text{-}74}$.

**D**, $^1$H-$^{15}$N resonance assignment of *Sc*Tom20$_{37\text{-}183}$ spectrum. Shown are spectra of apo *Sc*Tom20 (gray) and in the presence of *Sc*Tom22$_{1\text{-}97}$.

**E**, *Sc*Tom20 spectra without (gray) and with WT (blue) and mutant L58E (light blue) *Sc*Tom22$_{1\text{-}97}$.

**F**, Assigned $^1$H-$^{15}$N resonances of *Rn*Tom20 protein. Shown are spectra of apo *Rn*Tom20 protein (gray) and upon adding either *Sc*Tom22$_{1\text{-}74}$ (blue) or presequence peptide pSu9$_{1\text{-}25}$ (red).

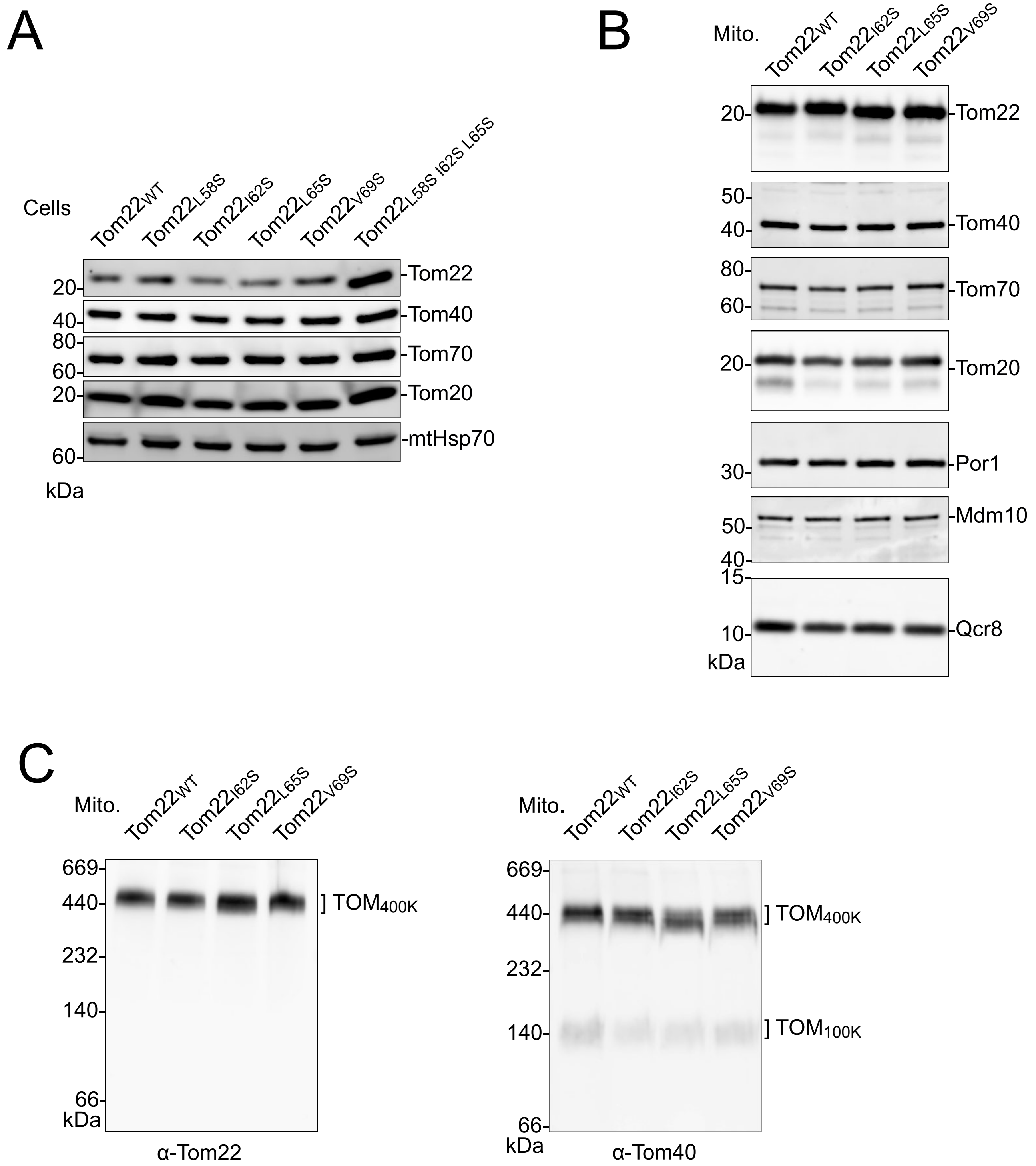


**Figure S8. Tom protein and TOM complex levels in strains expressing Tom22$_{PPDE}$ point mutants, related to Figure 2G.**

**A**, Whole cell extract/immunodecoration reveals similar levels as in wild-type.

**B**, Isolated mitochondria subjected to SDS-PAGE/immunodecoration show protein levels comparable to wild type (WT).

**C**, Isolated mitochondria were lysed with digitonin and subjected to blue-native PAGE/immunodecoration. The Tom22$_{PPDE}$ point mutant variants are efficiently assembled into the mature TOM$_{400K}$ complex.

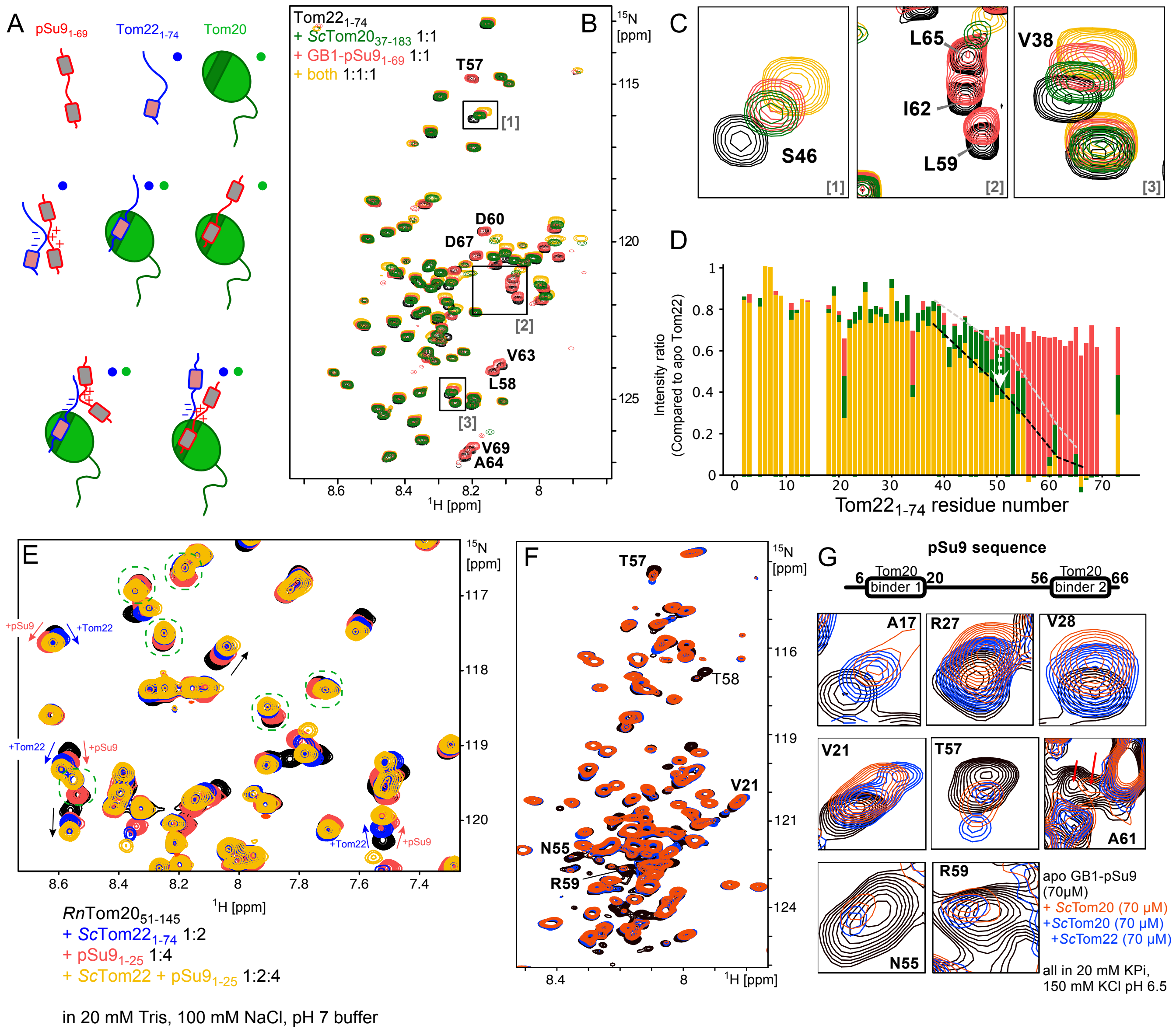

**Figure S9. Monitoring interactions and ternary-complex formation of Tom22, pSu9 and Tom20 using NMR.**

**A**, Possible species present in a sample containing $pSu9_{1\text{-}69}$, $Tom22_{cyt}$ and $Tom20_{cyt}$. Data obtained with samples comprising $^{15}$N-labeled $Tom22_{1\text{-}74}$ are shown in B-D; only the species marked with a blue dot in (**A**) are visible in these data. Spectra of $^{15}$N-labeled *Rn*$Tom20_{51\text{-}145}$ are depicted in E, and report on species marked with a green dot in A.

**B, C**, NMR spectra and peak intensities of Tom22 apo (black), in the presence of Tom20 (green), in the presence of pSu9 (red) and both (yellow).

**D**, Intensity ratios of Tom22 signals from the spectra in (**B**), showing the intensity with binding partner(s) divided by the intensity in the apo spectrum.

The addition of pSu9 only (red) leads to modest CSPs, but no significant intensity changes. This shows that Tom22 remains highly flexible in this interaction. Addition of Tom20 only leads to strong intensity reduction around the $Tom22_{PPDE}$, e.g. L59, I62, 65 (highlighted in C). This intensity reduction is expected for the binding of a highly flexible protein to a larger folded protein (due to the slower overall tumbling).

The simultaneous presence of pSu9 and Tom20 (yellow) leads to a further intensity reduction in Tom22. This can only be explained by the fact that a higher population of Tom22 is bound to Tom20 when pSu9 is present. This observation can be understood by the fact that the dynamic interactions of the three proteins essentially increase the local concentration, and thus the population of ternary (transient) complex. The pattern of intensity and chemical-shift perturbations show that $Tom22_{1\text{-}74}$ must have interactions with both proteins.

**E**, A complementary view of these interactions is provided through samples in which *Rn*Tom20 is NMR visible. Chemical-shift changes induced by pSu9 or Tom22 are indicated with green and blue arrows. Importantly, in many cases the chemical shift perturbation induced by the simultaneous presence of both Tom22 and pSu9 is larger than the one induced by only one protein, showing that the population of complex is increased. This data shows that Tom20 binds both proteins (either as a ternary complex or as a binary complex). In the latter case, the data clearly show that this binding and release is fast, i.e. that both compete for the same site.

Collectively, although indirect, these data strongly suggest that ternary complexes are formed when all three proteins are present. A different interaction pattern of Tom22 – undergoing hydrophobic interactions between $Tom22_{PPDE}$ and electrostatic interactions with its negatively charged residues (see Fig. S4G) – is indicated from CSPs and intensity changes. **F,G**, Ternary complex formation observed by $^{15}$N-labeled GB1-pSu9 showing the full spectrum (F) and zooms onto selected peaks (G). The spectra were acquired at 150 mM KCl concentration. The color code of the spectra is as indicated (apo, with Tom20, with both Tom20 and Tom22); the sequence of $pSu9_{1\text{-}69}$ is indicated above. There are different cases: in some, the addition of Tom22 to the Tom20/pSu9 sample leads to further shifts (e.g., A17, R27, V28); most of these are in the linker region between the two binders; in some, the addition of Tom22 leads to a reduction of the Tom20-induced effect, presumably due to competition of binding to Tom20 (e.g., V21, T57, N55, R59, A61). Some residues (not shown as zooms) do not significantly change upon addition of Tom22. These data support the idea of a ternary complex formation, with on/off dynamics on fast time scales and slower (µs-ms) time scales (for the binder 2 region).

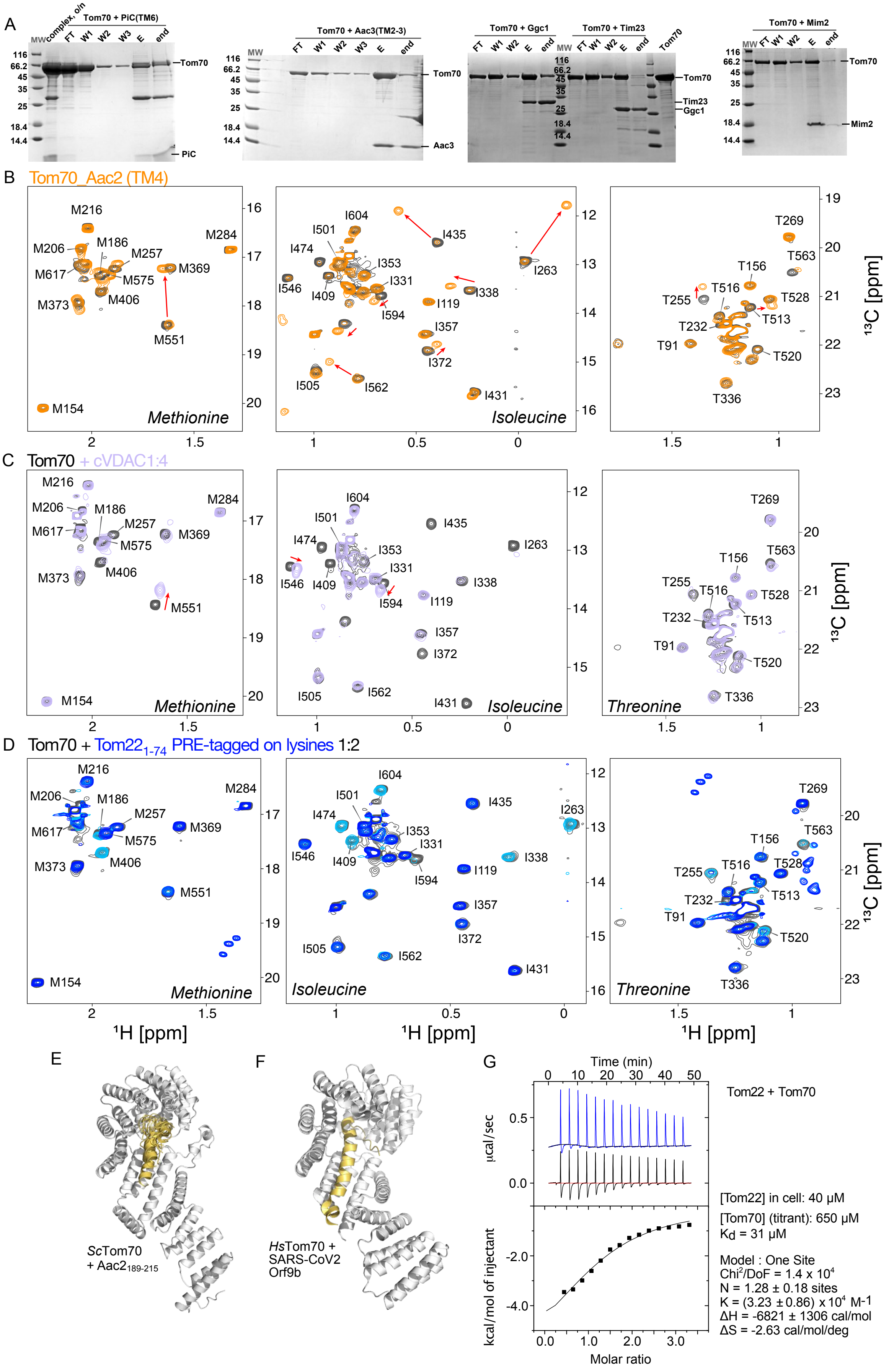
A
MW
complex, o/n
Tom70 + PiC(TM6)
FT
W1
W2
W3
E
end
Tom70
PiC
Tom70 + Aac3(TM2-3)
Aac3
Tom70 + Ggc1
Tom70 + Tim23
Tim23
Ggc1
Tom70 + Mim2
Mim2
116
66.2
45
35
25
18.4
14.4
B
Tom70_Aac2 (TM4)
Methionine
Isoleucine
C
Tom70 + cVDAC1:4
Threonine
D
Tom70 + Tom22 1-74 PRE-tagged on lysines 1:2
M216
M206
M186
M257
M284
M617
M575
M369
M373
M406
M551
M154
I604
I501
I474
I353
I331
I546
I409
I594
I435
I263
I338
I119
I357
I372
I505
I562
I431
T269
T563
T156
T255
T516
T528
T513
T232
T91
T520
T336
¹³C [ppm]
¹H [ppm]
E
ScTom70
+ Aac2 189-215
F
HsTom70 +
SARS-CoV2
Orf9b
G
Time (min)
0
10
20
30
40
50
µcal/sec
0.5
0.0
kcal/mol of injectant
-2.0
-4.0
0.0
1.0
2.0
3.0
Molar ratio
Tom22 + Tom70
[Tom22] in cell: 40 µM
[Tom70] (titrant): 650 µM
Kd = 31 µM
Model : One Site
Chi²/DoF = 1.4 x 10⁴
N = 1.28 ± 0.18 sites
K = (3.23 ± 0.86) x 10⁴ M-1
ΔH = -6821 ± 1306 cal/mol
ΔS = -2.63 cal/mol/deg

**Figure S10. (Preceding page) NMR study of *Sc*Tom70 and its complexes.**

**A**, SDS-PAGE analysis of $Tom70_{39\text{-}617}$ complexes with different client proteins. Transmembrane helix 6 (TM6, residue 251-299) of the mitochondrial phosphate carrier protein 2 (PiC; UniProt ID: P23641) was expressed as a fusion construct with Tom70. The fusion protein included a TEV protease recognition site between Tom70 and PiC(TM6), allowing for site-specific cleavage. Following TEV protease treatment, the cleaved PiC fragment was isolated via reverse Ni-NTA purification, as shown in the gel. Full-length mitochondrial GTP/GDP carrier protein 1 (GGC; UniProt ID: P38988), mitochondrial inner membrane translocase subunit Tim23 (UniProt ID: P32897), mitochondrial import protein 2 (Mim2; UniProt ID: Q3E798) and transmembrane helices 2 and 3 (TM2-3) of mitochondrial ADP/ATP carrier protein 3 (AAC3; UniProt ID: P18238) were purified in denaturing conditions and complex with Tom70 was formed using pull-down approach.

**B**, Spectrum of Met, Ile (Cδ1) and Thr methyl groups of $Tom70_{39\text{-}617}$ in complex with $Aac2_{189\text{-}215}$ (TM4), shown in orange. The complex sample was produced as a fusion construct, and TEV-cleaved prior to NMR analysis. The spectrum shown in black is from the corresponding apo sample, produced as a Tom70-TEV-$His_6$ construct, and equally subjected to TEV cleavage, such that the entire sequence, including its C-terminus, is identical.

**C**, Met, Ile (Cδ1) and Thr methyl spectrum of $Tom70_{39\text{-}617}$ with a four-fold molar excess of $cVDAC_{257\text{-}279}$ peptide. In both cases (purple: with cVDAC peptide; black: apo), the same *Sc*$Tom70_{39\text{-}617}$ protein was used, ending cleanly with the native "GLM" C-terminus. Note that the strongest effects are in the intensities, more than in CSPs.

**D**, Spectrum of *Sc*$Tom70_{39\text{-}617}$ in the presence of non-isotope-labeled *Sc*$Tom22_{1\text{-}74}$ in its untagged form (black), with MTSL tags on lysines in paramagnetic form (dark blue) and after addition of ascorbic acid to the MTSL-tagged sample (light blue).

**E**, AlphaFold2 model of *Sc*$Tom70_{89\text{-}617}$ and *Sc*$Aac2_{189\text{-}215}$.

**F**, Structure *Hs*$Tom70_{89\text{-}617}$ and SARS-CoV2 Orf9b (PDB 7DHG).

**G**, Isothermal titration calorimetry of the Tom70-Tom22. Shown are the raw thermograms (top) and the binding isotherms (bottom). The concentrations of the protein in the cell (Tom22) and of the titrant Tom70 is indicated in the legend. Titration of Tom70 into Tom22 is represented in black while titration into buffer (negative control) is shown in blue.

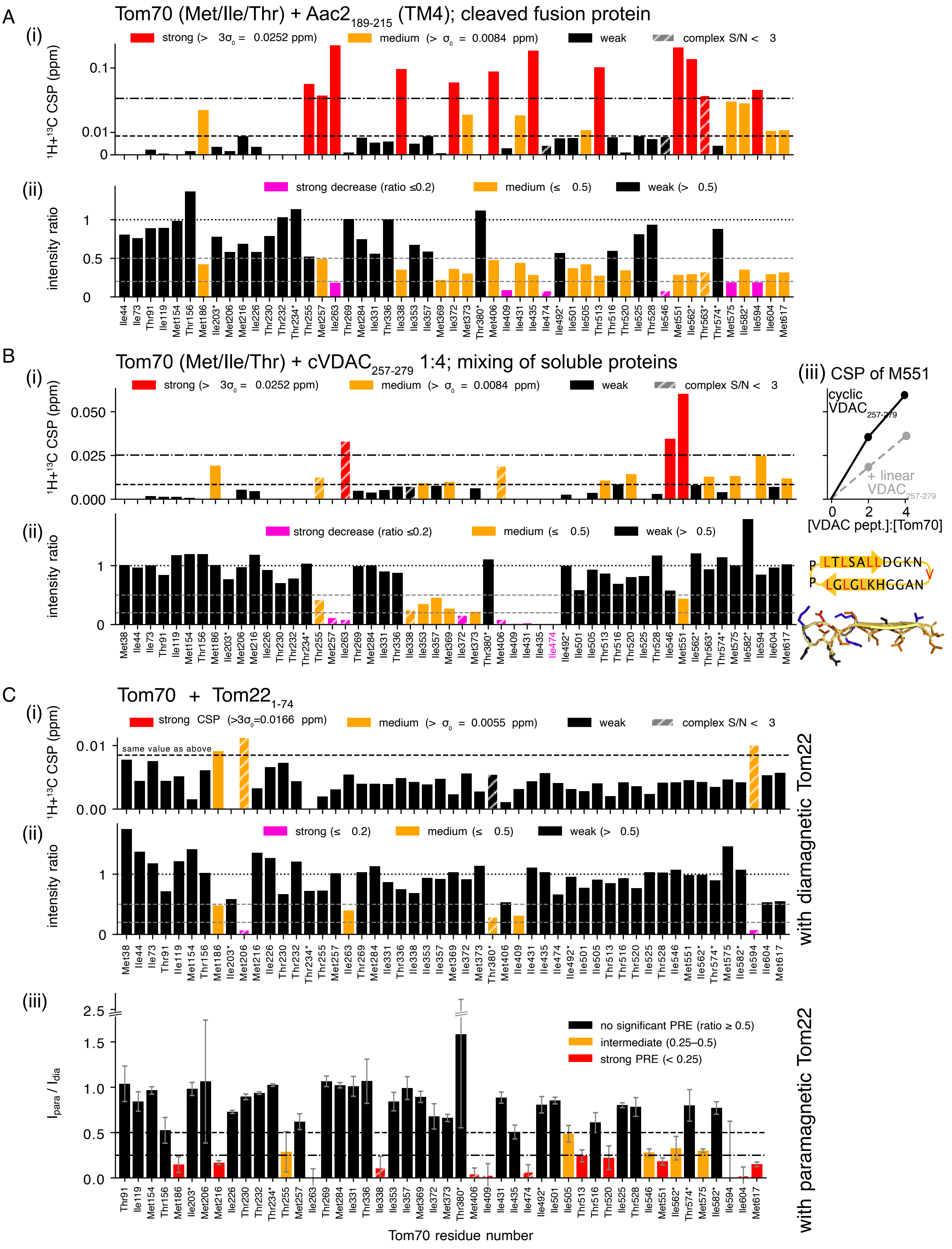
A
Tom70 (Met/Ile/Thr) + Aac2 189-215 (TM4); cleaved fusion protein
(i)
strong (> 3σ0 = 0.0252 ppm)
medium (> σ0 = 0.0084 ppm)
weak
complex S/N < 3
1H+13C CSP (ppm)
(ii)
strong decrease (ratio ≤0.2)
medium (≤ 0.5)
weak (> 0.5)
intensity ratio
B
Tom70 (Met/Ile/Thr) + cVDAC 257-279 1:4; mixing of soluble proteins
(iii) CSP of M551
cyclic VDAC 257-279
+ linear VDAC 257-279
[VDAC pept.]:[Tom70]
P LTLSALLDGKN
P LGLGLKHGGAN
C
Tom70 + Tom22 1-74
strong CSP (>3σ0=0.0166 ppm)
medium (> σ0 = 0.0055 ppm)
same value as above
strong (≤ 0.2)
with diamagnetic Tom22
(iii)
no significant PRE (ratio ≥ 0.5)
intermediate (0.25–0.5)
strong PRE (< 0.25)
Ipara / Idia
Tom70 residue number
with paramagnetic Tom22

**Figure S11. (Preceding page) Quantitative analysis of the interaction of *Sc*Tom70 with two preprotein fragments and with *Sc*Tom22 by methyl-NMR.** In (A), (B), and (C), the sub-panel (i) shows the combined $^1H/^{13}C$ chemical-shift perturbation $\mathrm{CSP} = \sqrt{\Delta\delta_H^2 + (\alpha\,\Delta\delta_C)^2}$ with residue-type-specific $\alpha$ (Ile 0.171, Met 0.22, Thr 0.19), and the sub-panel (ii) shows the intensity ratio $I_{complex}/I_{apo}$.

**A**, CSP and intensity effects induced by the presence of $Aac2_{189\text{-}215}$. The complex was obtained by cleavage of a *Sc*$Tom70_{39\text{-}617}$-TEV-$Aac2_{189\text{-}215}$ construct. The colors of the bars are a visual guide only; in (i), red and orange bars refer to CSPs larger than the indicated values (determined jointly for both Aac and cVDAC). In (ii), color codes refer to the ratio of peak intensity in the complex compared to apo.

**B**, CSP and intensity effects induced by the presence of a cyclic $VDAC_{257\text{-}279}$ construct, which was added at a 1:4 ratio. Sub-panel (iii) shows that addition of a linear version of the peptide leads to significantly smaller CSPs than the cyclic version, as illustrated with M551, the residue with the largest CSP. A model of the cyclic peptide shows segregation of hydrophobic and hydrophilic residues on the opposing faces of the β-turn.

**C**, Effects induced by *Sc*$Tom22_{1\text{-}74}$, which was added in a 1:2 ratio (150 µM MIT-labeled Tom70, 300 µM Tom22). Sub-panels (i) and (ii) show the effects of a diamagnetic version of Tom22, while sub-panel (iii) shows the paramagnetic effects on Tom70 methyl groups by a paramagnetically tagged Tom22 (attached at lysines, using 1-Oxyl-2,2,5,5-tetramethylpyrroline-3-carboxylate N-hydroxysuccinimide ester). In the construct *Sc*$Tom22_{1\text{-}74}$ there are only four lysines: K8 and K29 are in the flexible N-terminal part which our Tom22-detected data show to not be involved in the interaction; K66 and K73 are both within or close to the PPDE. The PRE effects are a proxy of the location residues 66 and 73.

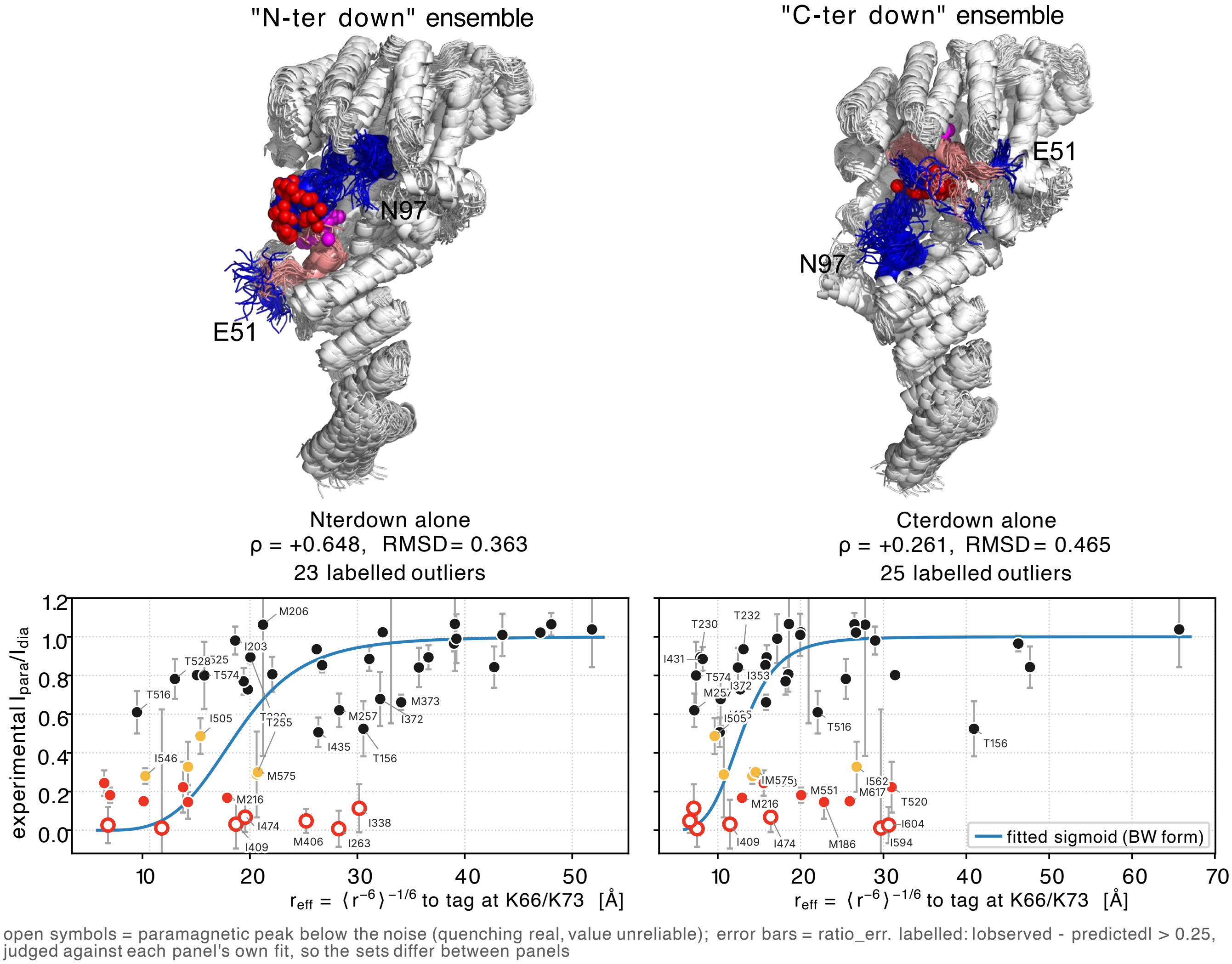


**Figure S12. Structural insights into the ensembles of conformations of the Tom70-Tom22 complex from MD and experimental PRE data.** Two different ensembles of conformations of the complex have been generated using all-atom MD simulations, starting from two different AlphaFold2 structural models of Tom70-Tom22$_{51\text{-}97}$; residues 1-50 have not been used, because Tom22-detected data show that this part remains free and flexible in the complex (Figure 3 D-F). In these two models, the orientation of Tom22 differs, i.e., the N-terminal side including the PPDE is either on the lower side or deeply inserted into the binding pocket. From each of these two models, a 0.5 µs long MD simulation was performed; along these trajectories, the binding poses remain overall stable, and the two ensembles are called here "N-ter down" and "C-ter down" ensemble, to reflect the different orientations. (The magenta and red spheres indicate the CA positions of the lysine residues 66 and 73, respectively; the other two lysines of the 1-74 construct are in the highly flexible N-terminal part that does not interact with Tom70.) We took one snapshot every 10 ns and built the OXYL-1-NHS tag onto the lysine side chain, using a 54-member rotamer set (lysine $\chi 3$, $\chi 4$ and the two linker torsions; backbone and $\chi_1$, $\chi_2$ from the MD). Rotamers were Boltzmann-weighted by a soft steric potential against the protein rather than hard-rejected, in the spirit of MMM [122]. The weights $w_j^{(t,k)}$ are these populations. Then the effective distances of each methyl group to the radical position (placed in the middle of the N-O bond of the tag) was calculated as $r_{\text{eff}} =$

$$r_{\text{eff},i} = \left[ \frac{P}{N_s} \sum_{k=1}^{N_s} \sum_{t\in\{66,73\}} \sum_{j=1}^{N_r} w_j^{(t,k)} \left\| \mathbf{r}_{i,k} - \mathbf{R}_j^{(t,k)} \right\|^{-6} \right]^{-1/6}$$

The three sums are not equivalent. The rotamer weights are normalised per $(t, k)$ so that sum is a weighted average; the sum over tag sites is a true sum because the two centres' relaxation rates add; and the $1/N_s$ makes the structure sum an average. The symbols are: $i$, methyl group and $\mathbf{r}_{i,k}$ its position in space; $k$ structure, $N_s = 50$; $t$ tag site, Lys66 / Lys73; $j$ rotamer, $N_r = 54$; $\mathbf{R}_j^{(t,k)}$ spin centre = midpoint of the nitroxide N–O; $P$ per-site labelling occupancy, 0.757, derived from mass spectrometry-estimated populations of Tom22 tagged with 0, 1, 2, 3 or 4 tags; $w_j^{(t,k)}$ rotamer weights. These effective distances are plotted against the PRE data from Figure S11 (C,iii). The line indicates a sigmoidal form, fitted to the data, in the spirit of Battiste–Wagner [123]. Outlier methyls are labelled. This data suggests that the "N-ter down" ensemble is in better agreement with the experimental data. We also considered the possibility that the experimental data is explained by a mixture of these two ensembles. The best-fit mixture is dominated by the N-ter down ensemble (98%), and slightly improves the agreement with the experimental data ($\rho$=0.739, RMSD=0.304). Of note, this analysis only seeks to distinguish between two AlphaFold/MD derived models, and we do not imply that these ensemble models are experimentally-grounded structures. Even the better of the two models only incompletely explains the experimental data, which suggests that the complex is more heterogeneous than any of these two states. Moreover, while the experiment used PRE-tagged Tom22$_{1\text{-}74}$, the structural model uses the construct Tom22$_{51\text{-}97}$.

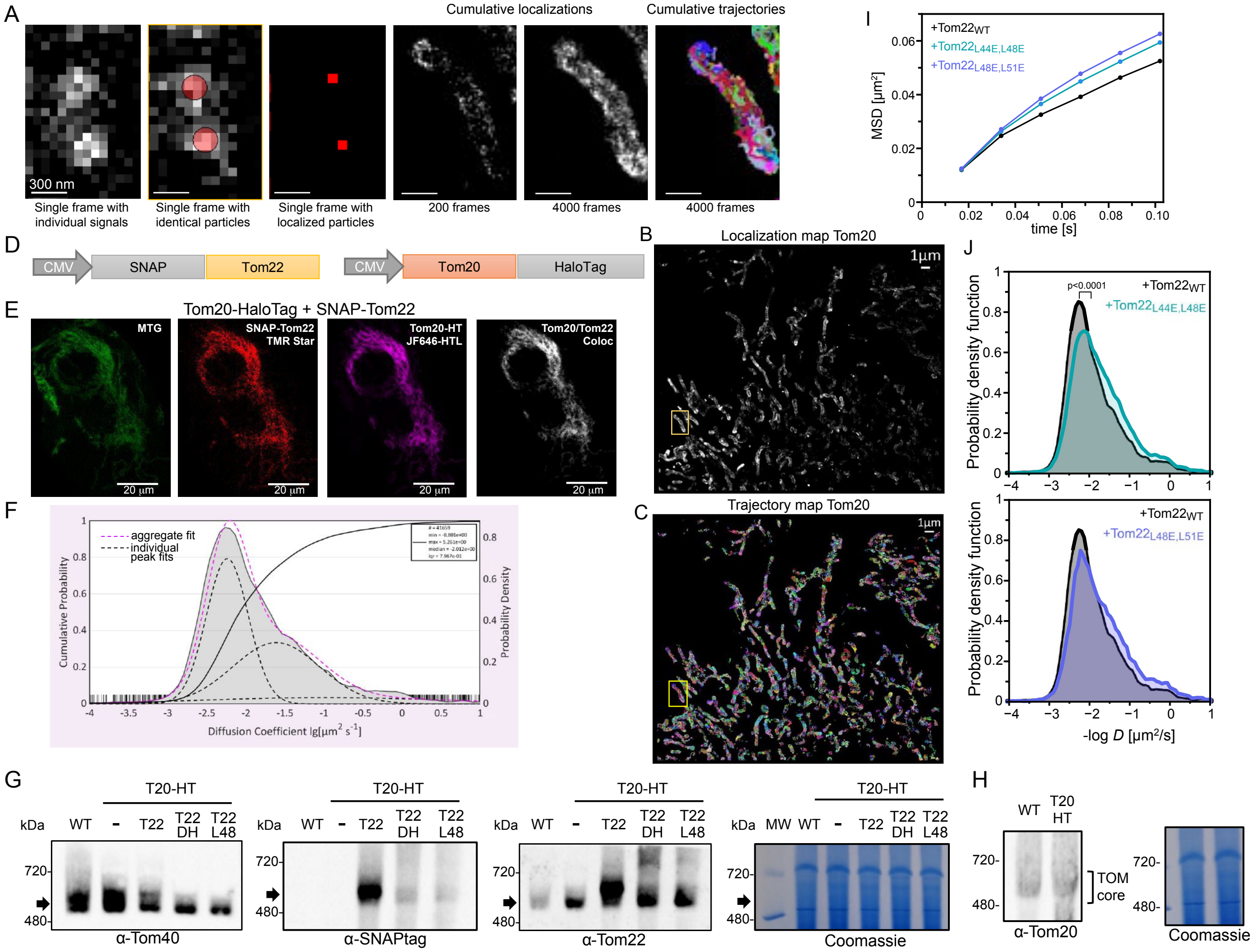

**Figure S13. Supplementary data for the single-particle tracking experiments.**

**A**, Process of single molecule localization and generation of cumulative localization and trajectory maps.

**B**, Cumulative images of localizations and **C**, individual trajectories, each trajectory in a different color. 4000 frames with 17 ms/frame recorded.

**D**, Constructs used for tracking experiments.

**E**, Mitochondrial localization of Tom20-HaloTag and SNAP-Tom22. Double transfected cervical HeLa cells were stained with MitoTracker Green (MTG) and supplied with substrates for the HaloTag: JF646-HTL and for the SNAP-tag: TMR-star.

**F**, Example of a Gaussian fit to estimate the population level of free and bound states.

**G**, Assembly of SNAP-tagged Tom22 variants into the TOM core complex. BN-PAGE, immunostaining of Tom40, SNAP-tag, Tom22 and Tom20, and Coomassie-stained gel. Molecular weight marker: MW (kDa). L48: Tom20-L48E-L51E; DH: Tom20-ΔHelix (kinetic data not shown, and data not used in this study).

**H**, Tom20 level in wt and Tom20-HaloTag (T20-HT) expressing cells.

**I**, Mean-square displacement (MSD) of fluorescently labeled Tom20 in HeLa cells expressing Tom22 either in its wild-type form, or with the mutations L44E,L48E or L48E,L51E.

**J**, Probability density function (PDF) of diffusion coefficients, *D*, of Tom20 in cells expressing different forms of Tom22. Statistical comparisons of all diffusion constants of all trajectories were done with non-parametric Kolmogorov-Smirnov test. Only trajectories with a minimum of 4 steps were analysed. Number of Tom20 trajectories analysed in the presence of Tom22 full length (41,659 trajectories, N=16 cells, n=4 technical replicates) Tom22-L44E-L48E (121,991; N=18, n=3), Tom22-L48E-L51E (56,401, N=19, n=3).
